# The adenosine analogue prodrug ATV006 is orally bioavailable and has potent preclinical efficacy against SARS-CoV-2 and its variants

**DOI:** 10.1101/2021.10.13.463130

**Authors:** Liu Cao, Yingjun Li, Sidi Yang, Guanguan Li, Qifan Zhou, Jing Sun, Tiefeng Xu, Yujian Yang, Tiaozhen Zhu, Siyao Huang, Yanxi Ji, Feng Cong, Yinzhu Luo, Yujun Zhu, Hemi Luan, Huan Zhang, Jingdiao Chen, Xue Liu, Ping Wang, Yang Yu, Fan Xing, Bixia Ke, Huanying Zheng, Xiaoling Deng, Wenyong Zhang, Chun-Mei Li, Yu Zhang, Jincun Zhao, Xumu Zhang, Deyin Guo

## Abstract

Severe acute respiratory syndrome coronavirus 2 (SARS-CoV-2), which causes the COVID-19 pandemic, is rapidly evolving. Due to the limited efficacy of vaccination in prevention of SARS-CoV-2 transmission and continuous emergence of variants of concern (VOC), including the currently most prevalent Delta variant, orally bioavailable and broadly efficacious antiviral drugs are urgently needed. Previously we showed that adenosine analogue 69-0 (also known as GS-441524), possesses potent anti-SARS-CoV-2 activity. Herein, we report that esterification of the 5’-hydroxyl moieties of 69-0 markedly improved the antiviral potency. The 5’-hydroxyl-isobutyryl prodrug, ATV006, showed excellent oral bioavailability in rats and cynomolgus monkeys and potent antiviral efficacy against different VOCs of SARS-CoV-2 in cell culture and three mouse models. Oral administration of ATV006 significantly reduced viral loads, alleviated lung damage and rescued mice from death in the K18-hACE2 mouse model challenged with the Delta variant. Moreover, ATV006 showed broad antiviral efficacy against different mammal-infecting coronaviruses. These indicate that ATV006 represents a promising oral drug candidate against SARS-CoV-2 VOCs and other coronaviruses.

## Introduction

The outbreak of COVID-19 pandemic, caused by SARS-CoV-2, has been continuing for over one year and has resulted in over 228 million confirmed infections and over 4 million reported deaths worldwide as of 20 September 2021 (WHO 2021b). SARS-CoV-2 is a positive-sense, single-stranded RNA virus belonging to the genus *Betacoronavirus* of the family *Coronaviridae* (Chen et al. 2020). Two other members of the same genus, namely severe acute respiratory syndrome coronavirus (SARS-CoV) and Middle East respiratory syndrome coronavirus (MERS-CoV), have caused outbreaks with substantial fatality rates in 2002 and 2012, respectively (Al-Tawfiq et al. 2014; Perlman et al. 2009). Given the repeated and accelerating emergence of highly pathogenic coronaviruses, it is increasingly important to develop broadly effective anti-coronaviral agents to combat the pandemics of COVID-19 and the future emerging CoVs.

Although the coronavirus has a certain proofreading ability (Robson et al. 2020), it still has a high mutation rate. Progressive mutational change in the virus is therefore inevitable after a number of replicative cycles, which leads to the emergence of new variants. At present, the WHO has classified many variants of the SARS-CoV-2 (WHO 2021a), including the variants of concern (VOC), such as the Alpha variant (B.1.1.7), Beta variant (B.1.351) and Delta variant (B.1.671.2). Delta variant is highly contagious, and has rapidly become the prevalent variant, contributing to the current wave of pandemic in India and worldwide and leading to vaccine breakthrough infections associated with higher viral load and long duration of shedding (Reardon 2021). These SARS-CoV-2 variants reduced the vaccine effectiveness and antibody protection (Harvey et al. 2021; Liu et al. 2021; Tegally et al. 2021). Currently available directly-acting antiviral drugs repurposed for treatment of COVID-19 patients showed limited efficacy in large-scale clinical trials (Consortium et al. 2021). Therefore, it is in urgent need to develop orally available and broadly efficacious anti-coronaviral agents against SARS-CoV-2 and its variants.

The non-structural protein 12 (nsp12) of SARS-COV-2 acts as the RNA-dependent RNA polymerase (RdRp), which catalyzes RNA template-dependent formation of phosphodiester bonds between ribonucleotides using ribonucleoside triphosphates as substrates, serving as the key component of the replication/transcription machinery (Hillen et al. 2020; Wang et al. 2020b). RdRps are considered as the primary targets for antiviral drug development in a wide variety of viruses (Kabinger et al. 2021; Picarazzi et al. 2020). Several nucleoside or nucleotide analogues, including remdesivir, AT-527, favipiravir and molnupiravir (EIDD-2801), originally developed by targeting the RdRp of other RNA viruses, have been repurposed for SARS-CoV-2 since the COVID-19 outbreak (Ghasemnejad-Berenji et al. 2021; Goldman et al. 2020; Good et al. 2021; Wahl et al. 2021; Wang et al. 2020a). Until now, remdesivir is the only FDA-approved antiviral drug for the treatment of COVID-19 patients (Beigel et al. 2020). However, remdesivir is administered by intravenous (IV) injection and thus has a clinical application limited to hospitalized patients with relatively advanced disease (Alanazi et al. 2019; Beigel et al. 2020; Consortium et al. 2021). We and others reported that the parent nucleoside of remdesivir, with CAS registry number 1191237-69-0 (named as 69-0, also known as GS-441524), potently inhibits SARS-CoV-2 infection in cell culture and mouse models infected with SARS-CoV-2 (Li et al. 2021; Pruijssers et al. 2020). 69-0 is a 1’-cyano-substituted adenosine analogue with broad-spectrum antiviral activities across multiple virus families (Cho et al. 2012; Huang et al. 2021; Murphy et al. 2018; Pedersen et al. 2019). However, 69-0 suffers from its disadvantage of poor solubility (water solubility of < 1 μg/mL) and poor oral pharmacokinetic (PK) profile (Li et al. 2021). In non-human primates, 69-0 has an oral bioavailability of 8.3% and poor plasma exposure, preventing it from further development into an oral drug (NCATS 2021).

In this study, we reported a series ester prodrugs of adenosine analogues 69-0 with improved antiviral potency. The isobutyryl nucleoside derivative of 69-0, named as ATV006, with improved oral PK profiles, potently inhibited the replication of SARS-CoV-2 and other CoVs. In three different animal models, ATV006 could efficaciously suppress the infection and pathogenesis of SARS-CoV-2 and its variants including the most prevalent Delta variant. These results indicate that ATV006 represents a promising drug candidate for the further clinical development against COVID-19 and other CoV diseases.

## Results

### Design and synthesis of prodrugs of adenosine analogs that target SARS-CoV-2 RNA polymerase

Although the adenosine analogue 69-0 was effective against SARS-CoV-2 in vitro and in mouse models, it suffers from poor oral bioavailability, which hampers its further development as oral drug as shown in our previous study(Li et al. 2021). To overcome this limitation, we devoted our efforts to synthesize 69-0 prodrugs by employing short chain fatty acid (SCFA) or amino acid to mask the polar hydroxyl- or animo-groups. For this, 21 compounds with different substitutions at the positions of **R^1^**, **R^2^**, **R^3^** and **R^4^ of 69-0** were designed and chemically synthesized (Fig 1; supplementary materials). In brief, the compounds ATV001-004 were synthesized from 69-0 via one-step acylation reaction with related acid anhydride in the presence of dimethylaminopyridine (DMAP) and ethylene glycol dimethacrylate (EDMA). To synthesize 5’-hydroxyl-acetylated compounds, the 2’,3’-hydroxyl moieties of 69-0 were firstly protected with acetonide in the presence of sulfuric acid. Then, different aliphatic acids, amino acids or benzoic acid were selectively coupled with the free hydroxyl group on C5’ position to produce the corresponding esters. Final deprotection of the acetonide with 6N hydrochloride acid (HCl) afforded the target compounds ATV005-024. All the compounds were purified by high performance liquid chromatography (HPLC), reaching a purity of above 95%.

**Figure 1.**
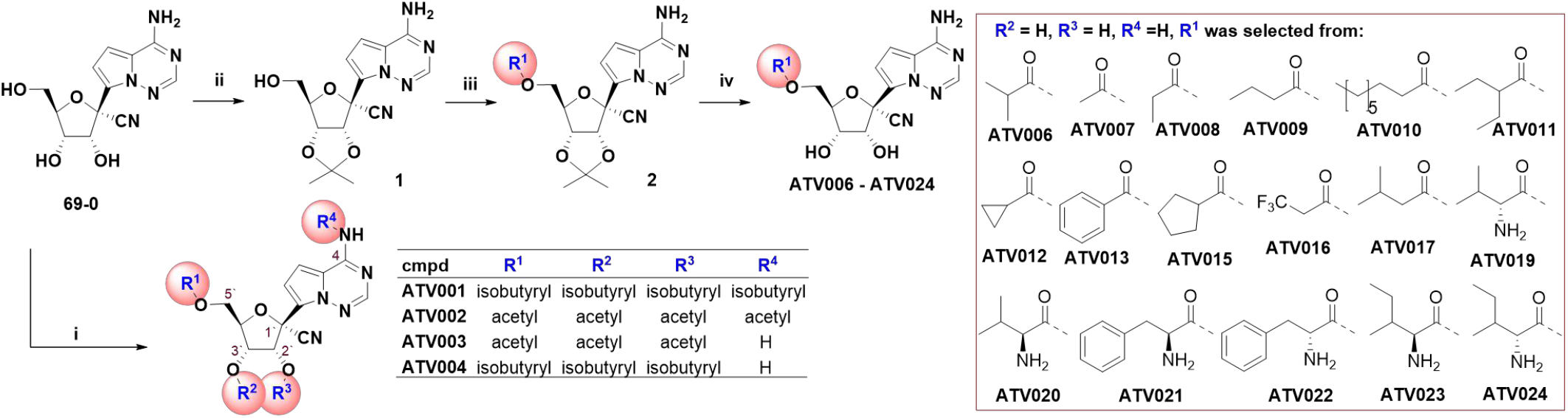
The chemical structure and synthesis of 69-0 prodrugs. Reagent and condition: i) anhydrides, DMAP, EDMA, ACN, 40 °C, 0.5 h; ii) 2,2-Dimethoxypropane, Conc. H_2_SO_4_, Acetone, rt~45 °C, 4 h; iii) carboxylic acid, DCC, DMAP, DCM, rt, 12 h; iv) 6 N HCl, THF, 0 °C, 7 h.

The antiviral effect of the compounds was initially evaluated by using a biosafe SARS-CoV-2 replicon system (pBAC-SARS-CoV-2-Replicon-Luc), established in our previous work (Jin et al. 2021), which carries all the genes essential to genome replication including that for RdRp and the luciferase reporter gene but does not produce infectious virus particles. We first tested the percentage inhibition of the replicon replication at 10 μM of each compound (Fig S1A) and then selected 17 compounds to measure their concentration for 50% of maximal effect (EC_50_) value, which ranged from 0.217 to 2.351 μM (Fig S1B). As shown in Fig S1A and S1B, the compounds ATV001 and ATV002 with isobutyryl amide or acetyl amide at the base moiety showed decreased antiviral activities probably due to the biostable amide group that obstructs the hydrogen bond formation between inhibitor and RNA template (Kokic et al. 2021; Wang et al. 2020b). Tri-esterification of the hydroxyl groups on C5’ (**R^1^**), C2’ (**R^2^**) and C3’ (**R^3^**) positions (ATV003-004) did not significantly change the activity while the mono-isobutyryl-modification of 5’-hydroxyl group (ATV006) markedly improved the inhibitory activities in the replicon system (EC_50_ value of 0.52 μM, about one-fold more potent compared to parent 69-0). We then kept the 2’ and 3’ hydroxyl group unchanged, while the R^1^ group was replaced with straight cyclic, branched SCFA, benzyl acyl-group or amino acid-group (ATV007-024). We found that some of the SCFA ester prodrug compounds displayed an improvement in potency relative to 69-0. Six compounds (ATV019-024) bearing L- or D-amino acid ester were designed to improve drug absorption by targeting the peptide transporter family 1 (PepT1) (Zhang et al. 2013). However, these compounds did not show improved activities against the replicon or improved permeabilities in Caco-2 cells (Table S1).

Together, SCFA esterification on C5’ position could generally improve the potency relative to 69-0. Then we selected six compounds from the SCFA group for further analysis of anti-SARS-CoV-2 activity with the live viruses of different variants in cell culture and animal models (Table 1).

**Table 1.** Anti-SARS-CoV-2 activity and cytotoxicity of the adenosine analogue prodrugs in comparison with remdesivir and 69-0

| Compound | EC<sub>50</sub> (μM) in Vero E6 cells |  |  | CC<sub>50</sub> (μM) |
| --- | --- | --- | --- | --- |
|  | SARS-CoV-2 (B.1) | SARS-CoV-2 (Beta, B.1.351) | SARS-CoV-2 (Delta, B.1.617.2) |  |
| Remdesivir | 2.279 | 1.780 | 1.645 | >50 |
| 69-0 | 1.709 | 1.354 | 0.957 | >50 |
| ATV006 | 1.360 | 1.127 | 0.349 | 128.00 |
| ATV009 | 1.329 | 1.484 | 0.492 | >50 |
| ATV010 | 0.696 | 1.002 | 0.457 | 44.62 |
| ATV011 | 2.117 | 2.302 | 0.408 | >50 |
| ATV013 | 2.262 | 2.434 | 0.965 | >50 |
| ATV017 | 2.188 | 2.847 | 0.428 | >50 |

### The adenosine analog prodrug ATV006 potently inhibits the replication of SARS-CoV-2 and its variants of concern

The antiviral efficacy of the six SCFA prodrugs (ATV006, ATV009-011, ATV013 and ATV017) was first evaluated in a cell culture model infected with different strains of SARS-CoV-2, including the early strain B.1, and two prevalent SARS-CoV variants of concern (VOC), the Beta (B.1.351) and Delta (B.1.671.2) variants (Fig 2, Table 1). The Vero E6 cells were treated with the compounds and then infected with SARS-CoV-2 at a multiplicity of infection (MOI) of 0.05, and the copy number of viral genome RNA in the cell culture supernatant was measured by the quantitative real-time polymerase chain reaction (qRT-PCR) 48 h post infection (hpi). As shown in Figure 2 and Table 1, the compounds showed improved potency against SARS-CoV-2 relative to remdesivir and 69-0, which was in consistency with the results of the SARS-CoV-2 replicon system. Among them, ATV010 exhibited low micromolar EC_50_ value with early strain B.1 and Beta variant, while ATV006 had an overall >4-fold potency improvement in inhibiting the replication of Delta variant, with EC_50_ reaching 0.3485 μM. We repeated the antiviral experiment of the compounds in Huh7 cells with B.1 strain, and the results showed that these compounds exhibited similar antiviral activity to that in Vero E6 cells (Fig S2). Together, these results indicate that SCFA-esterified compounds could potently inhibit the replication of SARS-CoV-2 and its variants.

**Figure 2.**
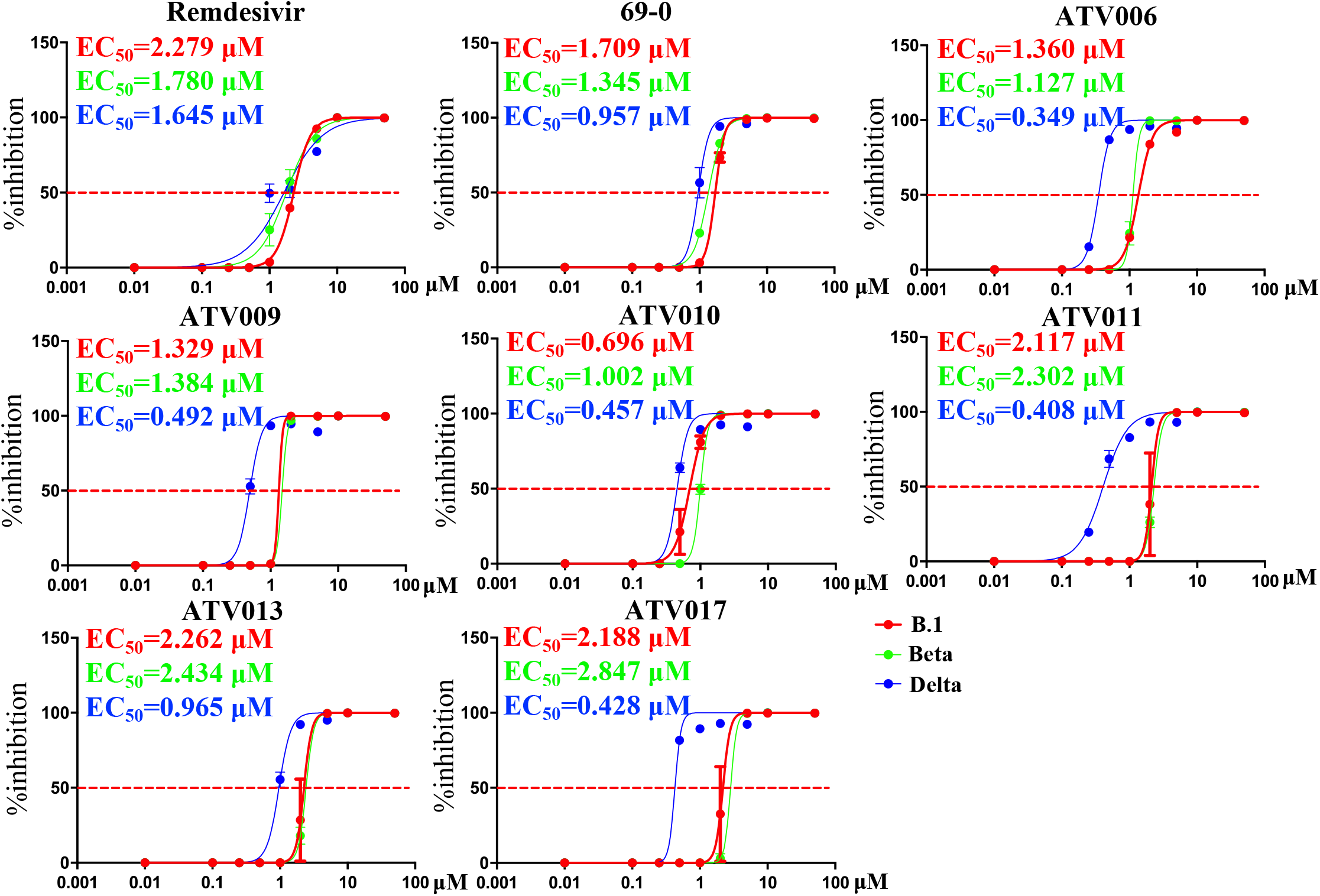
Antiviral activity of compounds against SARS-CoV-2 variants (B.1, Beta and Delta) in Vero E6 cells. Vero-E6 cells were infected with different strains of SARS-CoV-2 variant (B.1, Beta and Delta) at an MOI of 0.05 and treated with dilutions of compounds (0, 0.01, 0.1, 0.5, 1, 2, 5, 10 and 50 μM) for 48 h. Viral yield in the cell supernatant was then quantified by qRT-PCR. The values of EC_50_ of each compound were analyzed.

The cytotoxicity of the compounds was evaluated on Vero E6 cells with the CCK8 assays (Fig S3). The results showed that most of the compounds had low toxicity with CC_50_ > 50 μM, except for ATV010 with CC_50_ of 44.62 μM, suggesting that most compounds had excellent safety. The therapeutic index (CC_50_/ EC_50_) of ATV006 was as high as 367 in Vero E6 cells. Considering the potent inhibition against Delta variant and high selectivity against cell proliferation, ATV006 was selected for further studies.

### Pharmacokinetic properties of ATV006 in rats and cynomolgus monkeys

To assess the oral absorption of ATV006, the pharmacokinetic (PK) studies were conducted in rats and monkeys. Following oral dosing of 20 mg/kg in rats, ATV006 displayed high oral bioavailability (F%) of 98%, using the parent nucleoside 69-0 as an analyte (Table 2). The C_max_ of 11445 μg/L was achieved 0.83 h after the oral administration, indicating its effective blood exposure. Plasma concentration decayed with a half-life of 3.62 h (Fig 3A). In cynomolgus monkeys, C_max_ of averaged 2715 μg/L was reached in 1.5 h after the IG administration of 10 mg/kg ATV006. Plasma concentration decayed with a T1/2 of 4 h and the oral bioavailability was about 30% (Fig 3B). As a compound with an oral bioavailability of >10% has the potential for development as an oral drug (Martin 2005), ATV006 well met such standard as an oral drug candidate for further testing in animal models. Next, we explored the tissue distribution of ATV006 in the mouse model by measuring its parent nucleoside 69-0. The results revealed that oral administration of ATV006 achieved a broad distribution in liver and kidney as well as in lung, the major target organ to SARS-CoV-2 infection, indicating its potential of an oral drug for the treatment of COVID-19 (Fig 3C).

**Table 2.** Pharmacokinetic profile of ATV006 in Sprague-Dawley (SD) rat and cynomolgus monkey ^a^

| parameters | rat |  | monkey |  |
| --- | --- | --- | --- | --- |
|  | IV (4mg/kg) | IG (20mg/kg) | IV (5mg/kg) | IG (10mg/kg) |
| AUC<sub>last</sub> (μg/L*h) | 2347±354 | 11445±813 | 5960±490 | 3560±245 |
| T<sub>1/2</sub> (h) | 1.30±0.24 | 3.62±0.61 | 1.78±0.6 | 4.08±0.94 |
| T<sub>max</sub> (h) | 0.083 | 0.83±0.29 | 0.083 | 1.50±2.20 |
| C<sub>max</sub> (μg/L) | 3030±451 | 4017±359 | 3730±709 | 1080±651 |
| F (%) |  | 98% |  | 30.08% |

<sup>a</sup> n=3, ATV006 was administrated using the indicated route, and the parameters were calculated based on the LC-MS-MS analysis of parent nucleoside 69-0. IV, intravenous; IG, intragastric.

**Figure 3.**
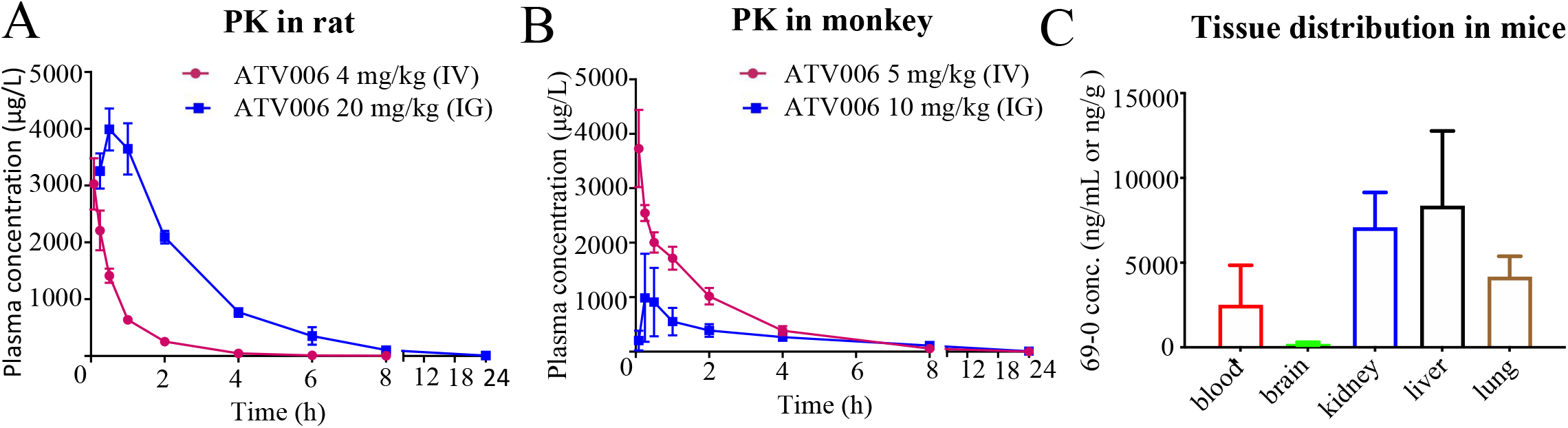
Pharmacokinetic profile of ATV006 in SD rats and cynomolgus monkeys. (A) Time-plasma concentration curve of the nucleoside following single IV (4mg/kg) or IG (20 mg/kg) administration of ATV006 to SD rats (n = 3, Mean ± SD). (B) Plasma concentration of the nucleoside following single IV (5 mg/kg) or IG (10 mg/kg) administration to cynomolgus monkeys (n = 3, Mean ± SD). (C) Tissue distribution of the parent nucleoside followed a single oral dose of 100 mg/kg ATV006 to C57BL/6 mice (n = 5, Mean ± SD).

### Orally administered ATV006 could effectively suppress SARS-CoV-2 replication in mouse models

We next investigated the in vivo antiviral activity of orally administered ATV006 in two different mouse models of SARS-CoV-2 (strain B.1) infection, one with humanized angiotensin-converting enzyme 2 (hACE2) (Sun et al. 2020b)and the other with adenovirus-delivered human ACE2 (Ad5-hACE2) (Sun et al. 2020a). The hACE2 transgenic mice were intranasally inoculated with SARS-CoV-2 (2×10^5^ plaque forming units (PFU) per mouse) and were treated with vehicle (control, n=6), ATV006 (500 mg/kg, IG, once daily, n=6) or ATV006 (250 mg/kg, IG, once daily, n=6), starting at 2 h prior to virus inoculation (Fig 4A) and continuing until 4 days post-infection. To better determine the replication levels of SARS-CoV-2, we detected both the genomic RNA (gRNA) and subgenomic RNA (sgRNA), the latter being produced by discontinuous synthesis and representing a biomarker of coronavirus replication (Hussain et al. 2005; Kim et al. 2020) (Fig S4). In the control group, both gRNA and sgRNA of SARS-CoV-2 reached high levels in the lung, indicating that the mouse infection model was well established. In contrast, SARS-CoV-2 RNAs were hardly detectable at day 4 in the ATV006 treatment groups (Fig 4B and 4C), demonstrating robust inhibition of SARS-CoV-2 replication by ATV006.

**Figure 4.**
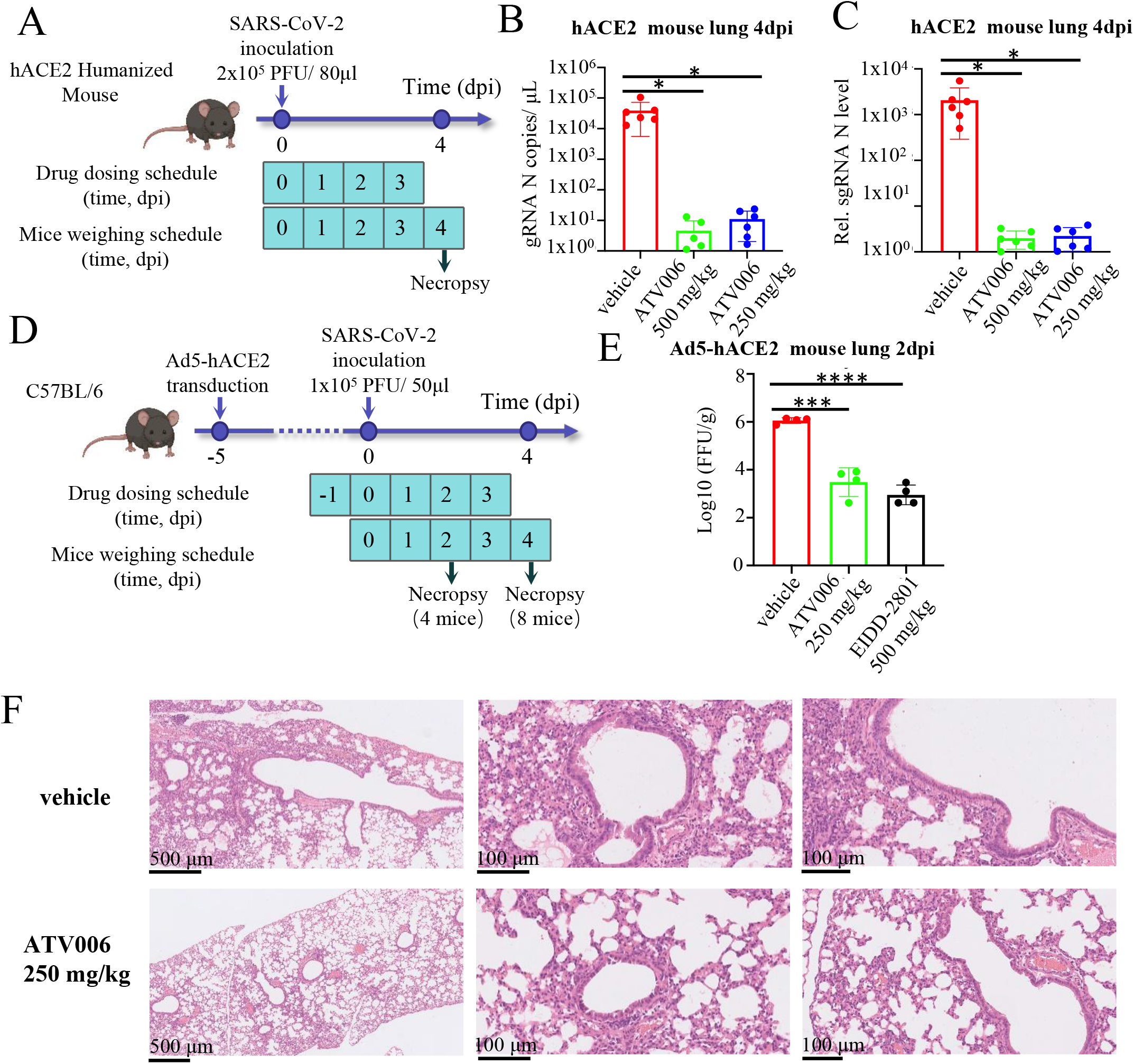
Anti-SARS-CoV-2 efficacy of ATV006 in hACE2 humanized and Ad5-hACE2 mouse model. (A) Schematic of the experiment viral infection in hACE2 humanized mice. hACE2 humanized mice were intranasally inoculated with B.1 original strain of SARS-CoV-2 (2×10^5^ PFU virus per mouse) and were administered with vehicle (control), ATV006 (250 mg/kg, IG, once daily), ATV006 (500 mg/kg, IG, once daily). Viral titers in the lungs at 4 dpi were measured by qRT-PCR analysis of gRNA N (B) and sgRNA N (C) of SARS-CoV-2. (D) Schematic of the experiment viral infection in Ad5-hACE2 mice. B.1 original strain of SARS-CoV-2 (1 × 10^5^ PFU virus per mouse) infected Ad5-hACE2 mice were administered with vehicle (control), ATV006 (250 mg/kg, IG, once daily), EIDD-2801 (500 mg/kg, IG, once daily) beginning at −2 hpi. (E) Viral titers in the lungs of treated or untreated Ad5-hACE2 mice at 2 dpi were measured by plaque assay. (F) Histopathology analysis of lung from vehicle group and ATV006 (250 mg/kg) group at 4 dpi. *p values ≤ 0.05; **p values ≤ 0.005; ***p values ≤ 0.0005; ****p values ≤ 0.0001.

We further tested the antiviral potency of ATV006 in the Ad5-hACE2 mouse model, which supports SARS-CoV-2 infection and pathogenesis in mouse lung (Sun et al. 2020a). The mice were inoculated intranasally with 1 × 10^5^ PFU virus per mouse and were then treated with vehicle (control, n=8) or ATV006 (250 mg/kg, IG, once daily, n=8) and EIDD-2801 (500 mg/kg, IG, once daily, n=8) starting at 1 day prior to virus inoculation (Fig 4D). EIDD-2801 was previously shown to effectively inhibit SARS-CoV-2 replication at 500 mg/kg dosage (Wahl et al. 2021) and used as a positive control. The virus titers were measured, and results showed that ATV006 (250 mg/kg) and EIDD-2801 (500 mg/kg) could significantly reduce the viral load and pathological damage of the lung (Fig 4E and 4F). Together, ATV006 showed potent anti-SARS-CoV-2 efficacy in different mouse models.

### ATV006 reduces lung damage and protects mice from death by infection of the Delta variant in the K18-hACE2 transgenic mice

We next tested the antiviral potency in the K18-hACE2 transgenic mice, which are susceptible to SARS-CoV-2 and can lead to death of infected mice (Oladunni et al. 2020). As the Delta variant of SARS-CoV-2 is the most prevalent variant, we specifically tested the efficacy of ATV006 against the Delta variant. The K18 hACE2 mice were intranasally inoculated with SARS-CoV-2 Delta variant (1×10^4^ PFU virus per mouse) and then treated with vehicle (control, n=11), ATV006 (250 mg/kg, IG, once daily, n=11), ATV006 (100 mg/kg, IG, once daily, n=8) or EIDD-2801 (500 mg/kg, IG, once daily, n=8) starting at 2 h prior to virus inoculation (Fig 5A) and continuing until 5 days post-infection (dpi). During the 9-day observation period, the mice in the control group gradually lost weight from the fourth day and died from the sixth day, and all died on the seventh day, but all mice of treatment groups survived (Fig 5B and 5C). At 3 dpi, we evaluated the abundance of both the viral gRNA of SARS-CoV-2 in the mouse lung and brain tissue by qPCR. The amount of viral RNAs of the treatment groups was significantly lower than that of the control group, with ATV006 (250 mg/kg) group having the strongest potency with reduction of viral RNA for more than 10,000 times in the lung (Fig 5D).

**Figure 5.**
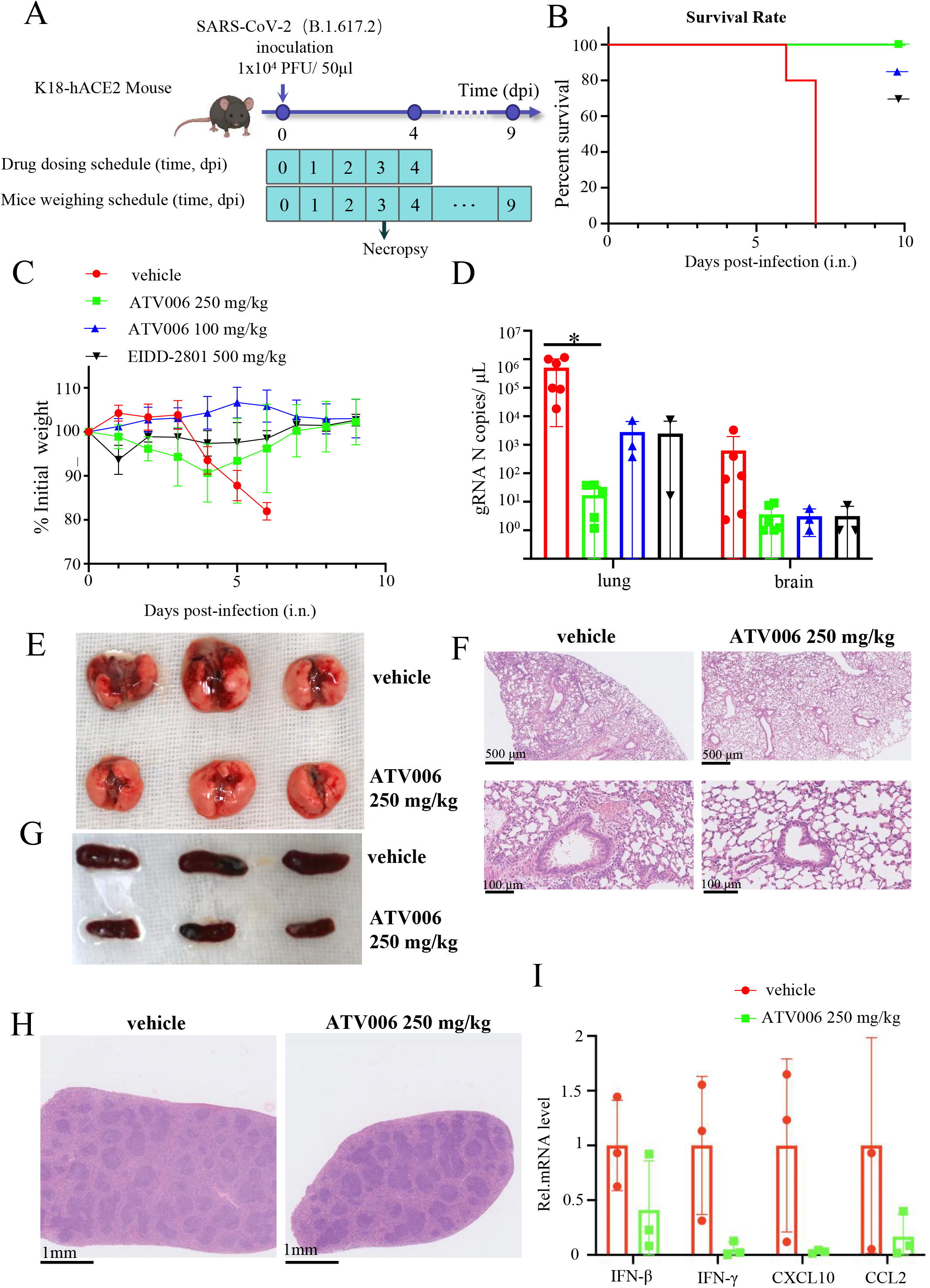
Anti-SARS-CoV-2 Delta variant efficacy of ATV006 in K18 hACE2 mouse model. (A) Schematic of the experiment. K18 hACE2 mice were intranasally inoculated with SARS-CoV-2 Delta variant (1 × 104 plaque forming units (PFU) virus per mouse) and were treated with vehicle (control, n=11), ATV006 (250 mg/kg, IG, once daily, n=11), ATV006 (100 mg/kg, IG, once daily, n=8) or EIDD-2801 (500 mg/kg, IG, once daily, n=8). (B) Survival curve. (C) Body weight curve. (D) Viral titers from lungs and brains tissue were harvested at 3 dpi and analyzed by qRT-PCR. Histopathology (F, H) and gross pathology (E, G) of lungs and spleens from vehicle group and ATV006 (250 mg/kg, IG, once daily). (I) Representative chemokines and cytokines assessment of the lung tissues harvested at 3 dpi of the vehicle group and 250 mg/kg ATV006 group. Total RNA were extracted from lung homogenates and IFN-β, IFN-γ, CXCL10 and CCL2 were analyzed by qRT-PCR. *p values ≤ 0.05; **p values ≤ 0.005; ***p values ≤ 0.0005; ****p values ≤ 0.0001.

Histopathological analysis and observation were performed with the lungs of the mice infected with SARS-CoV-2 at 3 dpi. The vehicle-treated mice showed multiple injuries, including inflammatory cell infiltration ranging from the trachea, peri-alveolar space, to the interstitium whereas ATV006-treated animals had markedly alleviated symptoms in the lungs (Fig 5E and 5F). Compared with the ATV006 treatment group, the spleen of the mice in the control group was significantly enlarged, and the white pulp was atrophied to varying degrees (Fig 5G and 5H). Furthermore, ATV006 markedly reduced the production of inflammatory cytokines and chemokines in the lung tissues (Fig 5I).

Together, our results showed that intragastric administration of ATV006 could efficiently inhibit SARS-CoV-2 replication, ameliorate SARS-CoV-2–induced lung lesions in vivo and prevent death of the mice infected by the Delta variant of SARS-CoV-2. These demonstrated the potential of ATV006 as an orally bioavailable anti–SARS-CoV-2 drug.

### ATV006 possesses broad antiviral activities against diversified coronaviruses

To explore the broad-spectrum antiviral activity of ATV006, we further tested it against other coronaviruses of the genera *Alphacoronavirus* and *Betacoronavirus* that are known to infect humans (Chen et al. 2020; Fung et al. 2019). Six coronaviruses were selected, including mouse hepatitis virus (MHV), feline infectious peritonitis virus (FIPV), porcine epidemic diarrhea virus (PEDV), canine coronavirus (CCoV), transmissible gastroenteritis virus (TGEV) and swine acute diarrhea syndrome coronavirus (SADS-CoV). Among these coronaviruses, MHV is a beta-coronavirus that is distantly related to human coronaviruses SARS-CoV-2, SARS-CoV and MERS-CoV.

We first compared the antiviral activities of ATV006 with that of remdesivir and 69-0 in MHV cell culture. The mouse L2 cells were infected with MHV-A59, a strain that infects the liver and brain of mice and causes acute hepatitis, encephalitis, and chronic demyelinating disease (Weiss et al. 2011), at a MOI of 0.1 and treated with different dilutions of the compounds. Antiviral activities were evaluated by qRT-PCR quantification of the viral copy number in the culture supernatant and intracellular fraction after 16 hpi. ATV006 showed robust anti-MHV activity (EC_50_ = 0.265 μM) compared to remdesivir (EC_50_ = 1.338 μM) and 69-0 (EC_50_ = 0.874 μM) in L2 cells (Fig 6A). Then we tested the antiviral activity of ATV006 against FIPV, CCoV, PEDV, TGEV and SADS-CoV, and the EC_50_ was 1.040 μM, 0.186 μM, 1.040 μM, 3.045 μM and 2.490 μM, respectively, in cell culture models (Fig 6B-6F). These results indicate that ATV006 has a broad-spectrum anti-coronavirus efficacy.

**Figure 6.**
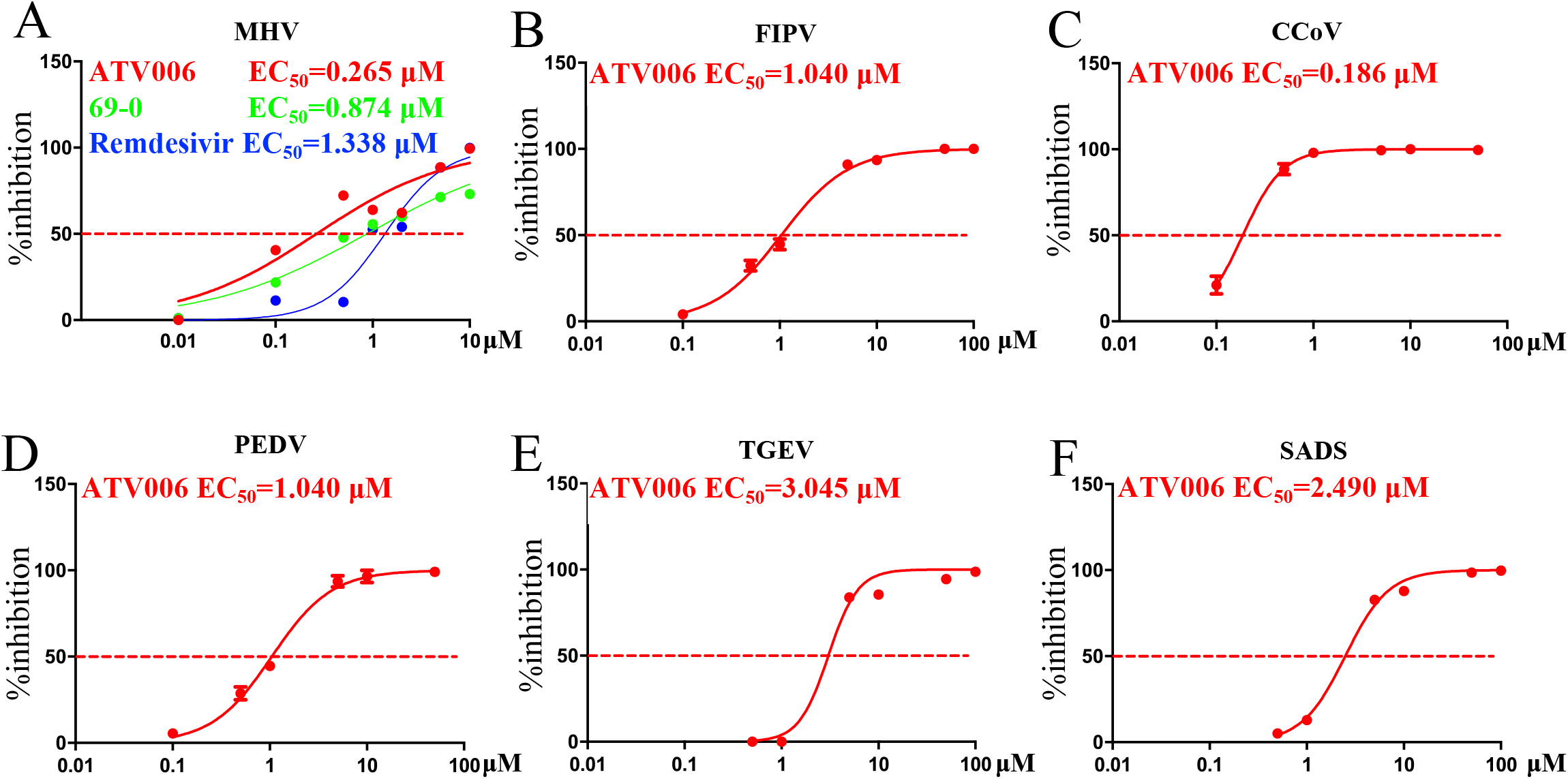
ATV006 has broad-spectrum antiviral activity among different coronaviruses. (A) L2 cells were infected with MHV-A59 at a multiplicity of infection (MOI) of 0.1 and treated with dilutions of remdesivir, 69-0 and ATV006. Antiviral activities were evaluated by qRT-PCR quantification of a viral copy numbers in the cultured supernatant after 16 h post infection. (B-F) The values of EC_50_ of ATV006 in (FIPV, CCoV, PEDV, TGEV and SADS) were analyzed.

We further evaluated the anti-coronavirus activity of ATV006 in MHV infection mouse model. The Balb/c mice were inoculated intranasally with 1 × 10^6^ PFU per mouse of MHV-A59, and treated with vehicle (control, n=5) or ATV006 (500, 250, 100, 50 mg/kg, IG, twice daily, n=5), starting at 3 hours prior to virus inoculation and continuing until 2 days post-infection (Fig 7A). We found that ATV006 treatments at different dosages could prevent the weight loss of the mice (Fig 7B). At 2 dpi, we measured the viral gRNA and sgRNA copy number and virus titer in mouse lung and liver (Fig S4 and Fig 7). It was found that the treatments of high dosages of ATV006 (250, 500 mg/kg) could effectively inhibit virus replication both in the lung and liver, and the 500 mg/kg group had 99% inhibition of virus replication in the lung and liver (Fig 7C-7F) while the treatment of low dosages of ATV006 (50, 100 mg/kg) significantly inhibited viral replication in the liver, but not significantly in the lung (Fig 7C-7F). Histopathological analysis demonstrated that the vehicle-treated mice showed inflammatory cell infiltration, whereas ATV006 (500 mg/kg)-treated mice had alleviated symptoms in the lungs at 2 dpi (Fig 7G). In addition, ATV006 significantly reduced the production of inflammatory cytokines and chemokines, such as IL-6, IL-1β and IFN-γ (Fig 7H). Due to the suppression of virus replication, IFN-β and ISGs (CXCL10) also significantly decreased (Fig 7H).

**Figure 7.**
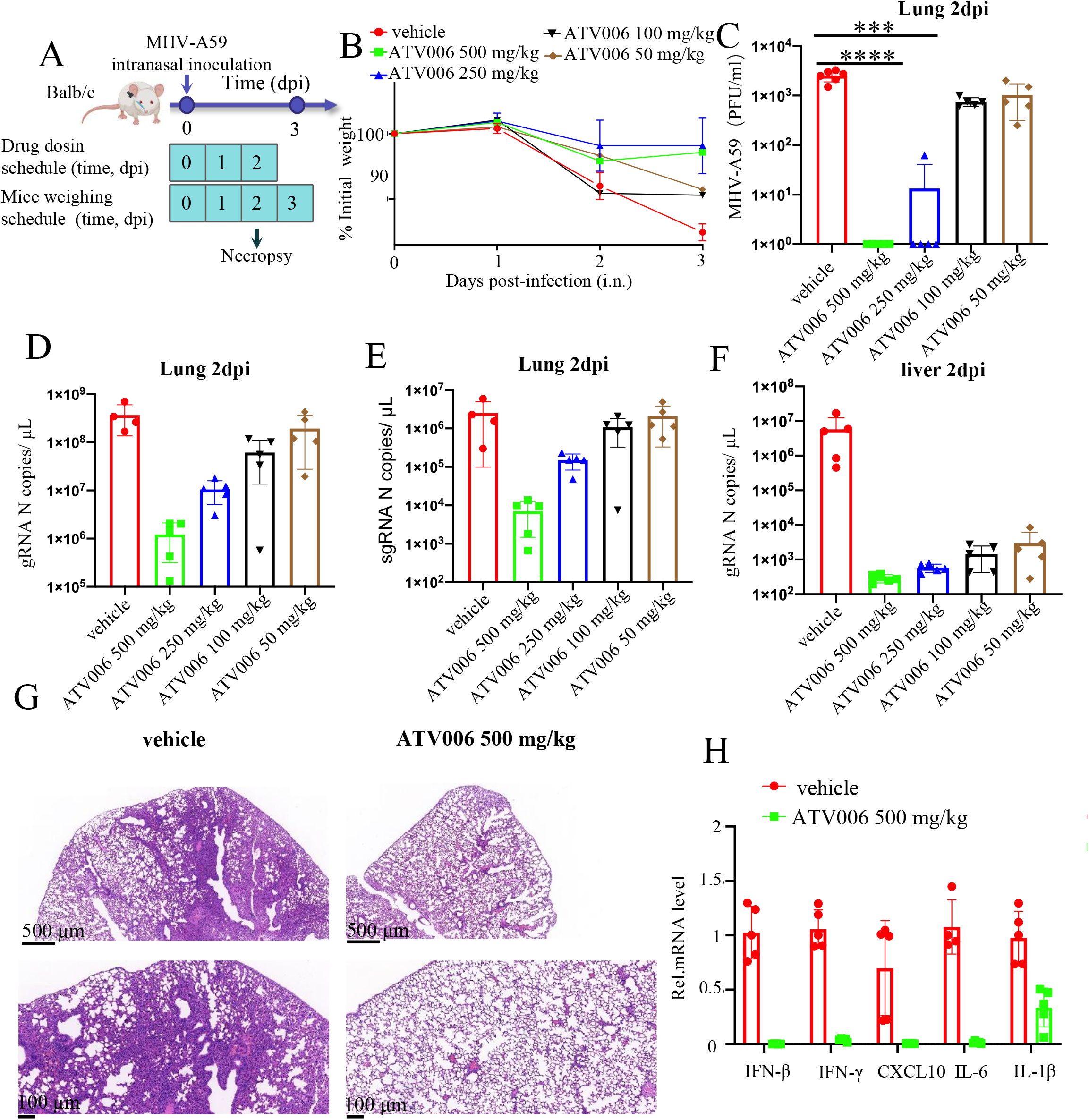
Anti-viral efficacy of ATV006 against MHV in vivo. (A)Schematic of the experiment. 5 weeks Balb/c mouse were intranasally inoculated with 1 × 10^6^ PFU per mouse of MHV-A59, and treated with vehicle (control, n=5) and ATV006 (500, 250, 100, 50 mg/kg, IG, twice daily, n=5 per group). (B) Body weight. (C, D, E, F) Viral titers from lungs and livers tissue were harvested at 2 dpi and analyzed by plaque assay and qRT-PCR. (G) Histopathology analysis of lung from vehicle group and ATV006 (500 mg/kg, IG, twice daily). (H) Representative chemokine and cytokine assessment of the lung tissues of the indicated groups, as detected in lung tissue homogenate at 2 dpi. *p values ≤ 0.05; **p values ≤ 0.005; ***p values ≤ 0.0005; ****p values ≤ 0.0001.

We next explored the lowest oral dosages of ATV006 that could protect MHV-A59-infected mice from death and weight loss. As shown in Figure S5, ATV006 could prevent mouse death and weight loss of mice at dosages of 5-50 mg/kg while the mouse treated with 2 mg/kg of ATV006 began to die 4 days post infection (dpi) and all mice died till 10 dpi. In comparison, the mice of the control group all died at 8 dpi and even the treatment with 2 mg/kg ATV006 prolonged the survival time and reduced the viral load in the liver by about six times (Fig S5B-S5D). IP administration of remdesivir (20 mg/kg) and IG administration of 69-0 (50 mg/kg) could also prevent death of the MHV-infected mice (Fig. S5) but the body weight of the mice was reduced in comparison with the mice treated with ATV006.

MHV mainly infects the liver of mouse and causes hepatitis. Therefore, we also performed intrahepatic (IH) inoculation to directly observe whether ATV006 had the effect of inhibiting virus replication in the liver and alleviating the symptoms of hepatitis (Fig S6A). Intriguingly, compared with the vehicle group, mice treated with the lowest concentration (2 mg/kg) had a 100-fold reduction in virus replication by measurement of the viral RNA load by qPCR, or the virus titer by plaque assay (Fig S6B, S6C and S6D). The serum ALT and AST values of ATV006-treated mouse were significantly lower than that of the control group (Fig S6E and S6F). The results of histopathological analysis also showed that after ATV006 treatment, the inflammatory cell infiltration in the liver was significantly reduced (Fig S6G). ATV006 could reduce the production of various inflammatory cytokines (Fig S6H). The above results demonstrate that ATV006 could also inhibit the replication of MHV-A59 in both the lung and liver in the mouse model and prevent the death of MHV-infected mice, indicating that ATV006 has a broad anti-coronavirus activity.

## Discussion

COVID-19, caused by SARS-CoV-2, is currently spreading globally, threatening human health and economic development. The vaccine-induced or naturally acquired protective herd immunity to interruption of transmission chains had been hampered by the rapid evolution and recurrent emergence of SARS-CoV-2 variants, such as Delta variant, one of the major variants of concern (VOC) (Liu et al. 2021; McCallum et al. 2021). Therefore, effective and broad-spectrum anti-SARS-CoV-2 drugs are desperately needed. Until now, remdesivir is the only FDA-approved small molecule antiviral drug for the treatment of COVID-19. However, the obligatory IV administration of remdesivir limits its clinical application only for hospitalized patients with advanced symptoms and its efficacy is limited in large scale clinical trials (Beigel et al. 2020). As COVID-19 is an acute infectious disease, antiviral treatment can exert its best effect at the early stage of the infection. In contrast, at the late stage of COVID-19 with hospitalized patients, anti-inflammatory therapy may play a major role in lessening the symptoms. Therefore, orally bioavailable anti-SARS-CoV-2 drugs suitable for outpatients are superior to injectable drugs applied to hospitalized patients.

Previously, we reported 69-0, the major metabolite and the parent nucleotide of remdesivir that targets the RdRp of SARS-CoV-2, has a better inhibitory activity on SARS-CoV-2 and MHV-A59 in vitro and in vivo (Li et al. 2021). However, the unfavorable oral PK prevents the further development of 69-0 (Li et al. 2021; NCATS 2021). To address this issue, herein, we synthesized a series of SCFA and amino acid prodrug of 69-0 aiming at overcoming its limitations. Among the compounds synthesized, the isobutyryl adenosine analogue ATV006 had improved oral absorption and potently inhibited the replication of SARS-CoV-2, especially the Delta variant (Fig 2). Compared to remdesivir, ATV006 is structurally simpler and easier to synthesize via a three-step transformation with 69-0 as starting material, which would reduce the cost and accelerate mass production. In addition, the orally active ATV006 is potentially more useful for the management of SARS-CoV-2 infection at the early stage.

After oral administration of ATV006, it is rapidly hydrolyzed by cellular esterases to produce the parent nucleoside 69-0 (Hsu et al. 2003; Lavis 2008), which then undergoes three steps of phosphorylation and is transformed to the active triphosphate form (Fig 3C), the same active component as that of remdesivir. Therefore, ATV006 shares the same mechanism of stalling SARS-CoV-2 polymerase as that of remdesivir (Kokic et al. 2021; Mackman et al. 2021; Wei et al. 2021; Yin et al. 2020a).

Several other small-molecule anti-SARS-CoV-2 antivirals are also under development, including the ones that block viral entry, inhibit Mpro, and target host immunity (Faheem et al. 2020; Good et al. 2021; Kabinger et al. 2021; Li et al. 2021; Minghua et al. 2021; Pruijssers et al. 2020; Sabbah et al. 2021; Vuong et al. 2021; Wahl et al. 2021; Zhang et al. 2021). Among them, EIDD-2801 was an orally available prodrug of nucleoside analog at phase 3 clinical trials and it was granted provisional approval in Australia in August, 2021. Clinical studies showed that a 5-day treatment of EIDD-2801 has a 100% SARS-CoV-2 clearance rate in the non-hospitalized patient (ClinicalTrials.gov NCT04405570) (Fischer et al. 2021). EIDD-2801 was reported to induce a two-step mutagenesis of viral RNA that is different to the inhibitory mechanism of ATV006 (Kabinger et al. 2021). We compared the antiviral efficiency of ATV006 with that of EIDD-2801 in the same experimental settings, and the results demonstrated that ATV006 possesses similar or higher anti-SARS-CoV-2 potency in different mouse models (Fig. 4 and Fig. 5).

Delta variant recently causes the sharp rise in SARS-CoV-2 cases worldwide. Compared to the original strain, Alpha and Beta variant, the Delta variant is more infectious and pathogenic (Motozono et al. 2021; Reardon 2021; Teyssou et al. 2021). Intriguingly, ATV006 showed an improved antiviral potency against Delta variant and the EC_50_ of ATV006 was about 3-4 times lower against Delta than the original strain and Beta variant (Fig 2). Oral treatments of ATV006 250 mg/kg, 100 mg/kg) as well as EIDD-2801 (500 mg/kg) effectively protected mice from severe weight loss and death induced by the infection of Delta variant (Fig 5C), indicating the high efficacy of ATV006 against the prevalent VOC. The signature mutations of the variants reside mainly in the receptor binding domain of spike protein (McCallum et al. 2021; Salleh et al. 2021) while the key residues of the nucleoside analog-binding and RdRp catalytic sites are 100% conserved (Fig S7). Therefore, we speculate that the increased sensitivity of Delta to ATV006 may be related to its high replication rate but not to the mutations in RdRp. The detailed mechanism needs to be further investigated in the future work. The analysis of 5,600 SARS-CoV-2 genomes also indicated that the adenosine analog, the parent nucleoside of remdesivir, does not seem to exert high selective pressure (Mari et al. 2021), suggesting that ATV006 and similar adenosine analogs have low risk to induce escape mutation and drug resistance.

The RdRp-encoding nsp12 is the most conserved protein in the coronaviruses. The key residues of nsp12 that are essential for RdRp enzymatic activity (Jin et al. 2021) and the binding with the parent nucleoside of remdesivir (Yin et al. 2020b) are 100% conserved throughout the coronaviruses of *Alphacoronavirus* and *Betacoronavirus* (Fig). Indeed, our current study demonstrated that ATV006 has a broad antiviral activity against different coronaviruses, including MHV, FIPV, CCoV, PEDV, TGEV and SADS (Fig 2 and Fig 6) (Haake et al. 2020; Izes et al. 2020; Korner et al. 2020; Laude et al. 1990; Lee 2015; Weiss et al. 2011; Zhou et al. 2018). The parent nucleoside 69-0 was previously reported to be broadly active against viruses that belong to the families *Paramyxoviridae, Coronaviridae and Filoviridae* (Cho et al. 2012; Huang et al. 2021; Lo et al. 2017; Murphy et al. 2018; Pedersen et al. 2019), indicating the potential for more broad antiviral application of ATV006. Collectively, our results demonstrated that ATV006 has potent and broad efficacy against SARS-CoV-2 and its variants of concern as well as other coronaviruses, thus representing a promising orally available drug candidate for the treatment for COVID-19 and emerging coronavirus diseases in the future.

## Materials and Methods

### Compounds, cells, and viruses

The preparation of novel compounds, ^1^H NMR, ^13^C NMR, HRMS analysis and HPLC purity are supplied in the Supplementary Information.

HEK 293T cells were obtained from American Tissue Culture Collection (ATCC). Rat lung epithelial cells (L2) and wild-type MHV-A59 were kindly provided by Rong Ye (Shanghai Medical School of Fudan University). African green monkey kidney Vero E6 cell line (Vero E6) was kindly provided by Dr. Hui Zhang (Sun Yat-sen University). Feline kidney cells (CRFK cells) and swine testicle cells (ST cells) were provided from Guangdong Province Key Laboratory of Laboratory Animals. HEK 293T, Vero E6, L2, CRFK and ST cells were cultured in DMEM supplemented with 10% FBS, 100 U/mL penicillin and streptomycin at 37 °C in a humidified atmosphere of 5% CO2.

SARS-CoV-2 (B.1, hCoV-19/CHN/SYSU-IHV/2020 strain, Accession ID on GISAID: EPI_ISL_444969) was isolated from a sputum sample from a woman admitted to the Eighth People’s Hospital of Guangzhou.

SARS-CoV-2 (B.1.351, SARS_CoV-2_human_CHN_20SF18530_2020, Accession ID on GWH: WHBDSE01000000) was isolated from a throat swab sample from a man admitted to the Eighth People’s Hospital of Guangzhou.

SARS-CoV-2 (B.1.617.2, GDPCC 2.00096) was isolated from a patient infected with SARS-CoV-2 Delta variant admitted in the Guangzhou Eighth People’s Hospital by Center for Disease Control and Prevention of Guangdong Province (Wang et al. 2021).

Canine coronavirus (CCoV), Porcine epidemic diarrhea virus (PEDV), Transmissible gastroenteritis virus (TGEV), Swine acute diarrhea syndrome coronavirus (SADS-CoV) and Infectious peritonitis virus (FIPV) were stored at Guangdong Province Key Laboratory of Laboratory Animals.

SARS-CoV-2 infection experiments were performed in the BSL-3 laboratory of Sun Yat-sen University or Guangzhou Customs District Technology Center.

MHV-A59 infection experiments were performed in the Biosafety Level 2 (BSL 2) laboratory of Guangdong Laboratory Animals Monitoring Institute. All animal studies protocols were approved by the Animal Welfare Committee and all procedures used in animal studies complied with the guidelines and policies of the Animal Care and Use Committee.

### SARS-CoV-2 replicon assays

The assays were performed following the manufacturer’s instructions (Promega Corporation, Fitchburg, WI, USA). In brief, the cells in 24-well plate transfected with 500 ng pBAC-SARS-CoV-2-Replicon-Luciferase plasmid and 10 ng RL-TK plasmid. After 6-8 h, the cells are transfected, the supernatant was discarded and replaced with fresh DMEM medium, followed by adding each compound (described in Table 1) to the media with the final concentration of 50 μM, 10 μM, 5 μM, 2 μM, 1 μM, 0.1 μM or 0.01 μM. After 60 h, cells were lysed in 200 μL Passive Lysis Buffer (PLB). Each lysate (20 μL) was transferred into 96-well white plate and then mixed with 20 μL Luciferase Assay Reagent II, followed by 20 μL of Stop & Glo solution. The luminescence values of the two-step reaction were recorded using a luminescence detector in Synergy H1 Hybrid Multi-Mode Reader.

### Anti-SARS-CoV-2 activity assays

Vero E6 and Huh7 cells were seeded at 2 × 10^4^ cells per well in 48-well plates. Cells were allowed to adhere for 16-24 h and then infected at MOI of 0.05 with SARS-CoV-2 for 1 h at 37°C. Then viral inoculum was removed, and cells were washed 2 times with pre-warmed PBS. Medium containing dilutions of compounds, or DMSO was added. At 48 hpi, supernatants or cells were harvested for qRT-PCR analysis. The dose-response curves were plotted from viral RNA copies versus the drug concentrations using GraphPad Prism 6 software.

### Anti-MHV-A59 Activity Assays

L2 cells were seeded at 1 × 10^5^ cells per well in 6-well plates. Cells were allowed to adhere for 16-24 h and then infected at MOI of 0.1 with MHV-A59 for 1h at 37°C. Then viral inoculum was removed, and cells were washed 2 times with pre-warmed PBS. Medium containing dilutions of compounds (ATV006, 69-0 and remdesivir), or DMSO was added. At 20 hpi, supernatants were harvested for qRT-PCR analysis. The EC_50_ values were calculated from the dose response curve. Viruses and cells are listed in Table S2. qPCR primers are listed in Table S3.

### Anti-FIPV, CCoV, PEDV TGEV and SADS Activity Assays

Cells were seeded in 48-cell plates for 24 h to reach 80% confluence and washed thrice with serum-free medium. Cells were infected with virus (0.01 MOI) at 37°C for 1 h. Medium containing dilutions of ATV006 or DMSO was added. After incubating at 37°C for 48h, cells and supernatants were harvested to determine viral loads using qRT-PCR. The EC_50_ values were calculated from the dose response curve. Viruses and cells are listed in Table S2. qPCR primers are listed in Table S3.

### qRT-PCR analysis

For SARS-CoV-2 RNA quantification, RNA was isolated by Magbead Viral DNA/RNA Kit (CWBIO). SARS-CoV-2 nucleic acid detection kit (Daan Company) is used to detect the virus.

For the detection of cellular viruses and tissue viruses and cytokines, total RNA was isolated from cells or tissue samples with TRIzol reagent under the instruction of the manufacturer. The mRNAs were reverse transcribed into cDNA by PrimeScript RT reagent Kit (Takara). The cDNA was amplified by a fast two-step amplification program using ChamQ Universal SYBR qPCR Master Mix (Vazyme Biotech Co., Ltd) or Taq Pro HS Universal Probe Master Mix (Vazyme Biotech Co., Ltd). GAPDH was used to normalize the input samples via the ΔCt method. The relative mRNA expression level of each gene was normalized to GAPDH housekeeping gene expression in the untreated condition, and fold induction was calculated by the ΔΔCT method relative to those in untreated samples. The qRT-PCR primers are listed in Table S3.

### CCK-8 cell viability assay

To investigate the effect of drugs on cell viability, Vero E6 cells were seeded in 96-well plates at a density of 20,000 cells/well and were treated with drugs at indicated concentrations (0, 0.01, 0.1, 1, 5, 10, 50 μM) for 48 h. Cell viability was tested by using Cell Counting Kit-8 (CCK-8, Bimake, B34302). The figures were plotted from viral RNA copies in supernatants versus the drug concentrations using GraphPad Prism 6 software.

### PK study in rat

Male SD rats (180-220 g, N = 3) were fasted for 12 h before drug administration. ATV006 was administered intravenously at 4 mg/kg or intragastrically at 20 mg/kg. Blood samples were collected from the jugular vein into anticoagulant EDTA-K2 tubes at 0.083, 0.25, 0.5, 1, 2, 3, 4, 6, 8 and 24 h for the IV group, and 0.25, 1, 0.5, 2, 3, 4, 6, 8 and 24 h for the IG group, respectively. All samples were centrifuged under 4000 rpm/min for 10 min at 4°C and the plasma (supernatants) were collected and stored at −65°C for future analysis. An aliquot of 50 μL each plasma sample was treated with 250 μL of acetonitrile. The samples were centrifuged under 4000 rpm/min for 10 min and filtered through 0.2 μm membrane filters. The concentration of analytes in each sample were analyzed by LC/MS/MS.

### PK study in cynomolgus monkeys

Three male cynomolgus monkeys (2 to 5 years of age, weighing 3 to 5 kg) were used were orally received AVT006 of 10 mg/kg on day 1. After administration, the blood samples for plasma were collected from a jugular vein into anticoagulant EDTA-K2 tubes at 0.083, 0.25, 0.5, 1, 2, 4, 8, 24 and 48 h. After a 3-day washout period, each animal was administered intravenously with ATV006 at a dose of 5 mg/kg, followed by bold collection from the jugular vein at the specified time points. All samples were centrifuged under 4000 rpm/min for 10 min at 4°C and the plasma (supernatants) were collected and stored at −65°C for future analysis. The concentration of analytes in each sample were analyzed by LC/MS/MS.

### Tissue distribution study in mice

Five C57BL/6 mice were fasted for 12-16 h before administered orally with a single dose of 100 mg/kg ATV006. After 1 h of dosing, 0.5 mL of the blood sample was taken from heart. Liver, kidney, lung, brain tissues were harvested. The blood sample was processed to the same method as in the rat PK studies. Tissue samples were homogenized and extracted with 70% methanol, after centrifuge under 4000 rpm for 10min, the supernatants were transfer to a clean tube and the concentration of analytes in each sample were analyzed by LC/MS/MS.

### Ad5-hACE2 Mice Study

The experiments were performed as previously described (Sun et al. 2020a). Mice were lightly anesthetized with isoflurane and transduced intranasally with 2.5 × 10^8^ FFU of Ad5-ACE2 in 75 μL DMEM. Five days post transduction, mice were infected intranasally with SARS-CoV-2 (1 × 10^5^ PFU) in a total volume of 50μL DMEM.

### Focus forming assay (FFA)

Vero E6 cells were seeded in 96-well plates one day before infection. Tissue homogenates were serially diluted and used to inoculate Vero E6 cells at 37°C for 1 h. Inoculate were then removed before adding 125 μL 1.6% carboxymethylcellulose per well and warmed to 37°C. After 24 h, cells were fixed with 4% paraformaldehyde and permeabilized with 0.2% Triton X-100. Cells were then incubated with a rabbit anti-SARS-CoV-2 nucleocapsid protein polyclonal antibody (Cat. No.: 40143-T62, Sino Biological), followed by an HRP-labeled goat anti-rabbit secondary antibody (Cat. No.: 109-035-088, Jackson Immuno Research Laboratories). The foci were visualized by TrueBlue Peroxidase Substrate (KPL) and counted with an ELISPOT reader (Cellular Technology). Viral titers were calculated as per gram tissue.

### MHV plaque assay

L2 cells were grown in 60-mm dishes to 70-80% confluence and infected with 1 mL of media containing viruses at dilutions ranging from 10-3 to 10-6. After 1 h at 37°C, the inoculate were removed. Seven ml of 0.95% agar (Amresco) in DMEM with 5% FBS was overlaid onto cells at 1-2h. Plaques were picked between 24 and 36 h. For plaque staining, 3 mL of agar containing 0.02% neutral red (Sigma-Aldrich) was overlaid onto cells. Six to eight hours later, the stained plaques were counted.

### AST/ALT assay

Blood was incubated at room temperature to allow coagulation and was then centrifuged to obtain serum; the serum was used for measurements of alanine transaminase (ALT) and aspartate aminotransferase (AST) levels using Assay Kit (Nanjing Jiancheng Bioengineering Institute).

### H&E Staining

Mice lung, liver and spleen dissections were fixed in zinc formalin and embedded with paraffin. Tissue sections (~4 μm) were stained with hematoxylin and eosin.

### Analysis of RdRp mutations of SARS-CoV-2 and its variants

There were four subtypes high-quality SARS-CoV-2 genomes sequences available in Global Initiative on Sharing All Influenza Data (GISIAD) (Shu et al. 2017). These high-quality genomic sequences of SARS-CoV-2 were screened out under the following criteria: 1) complete genome (>29000bp); 2) high coverage (only entries with <1% Ns and <0.05% unique amino acid mutations and no insertion/deletion unless verified by submitter); 3) low coverage excl (exclude entries with >5% NNNs). In total, we collected Alpha (B.1.1.7) (2021.5.1-6.10), Beta (B.1.351) (2019-2021.6.10), Gamma (P.1) (2019-2021.6.10), Delta (B.1.617.2) (2019-2021.6.10) were 108791, 13871, 18808, 29152 strains respectively. Subsequently, we eliminated 50 Ns or ns sequences. After screening, the remaining sequences were 92214, 6341, 13655, 21611 (Table S4).

SARS-CoV-2 genome of NC_045512.2 (Wu et al. 2020) was utilized as the reference sequence. Multiple sequences alignments were performed using the progressive method (FFT-NS-2) implemented in MAFFT (version 7.4) (Katoh et al. 2002). The whole genome mutation analysis was carried out used the pipeline provided by CoVa (Ali. et al. 2021) (version 0.2) software. Finally, the ggplot2 was used for drawing in R.

### Coronavirus RdRp Conservation Analysis

Virus data collection is derived from International Committee on Taxonomy of Viruses (ICTV) and NCBI, multi-sequence comparisons using mega 6, and data visualization using texshade (Beitz 2000) (Table S4). The sequence of 523aa-734aa of RdRp is selected for display, where K545, R555, S682, N691 and D760 are the key sites for remdesivir binding to RdRp, and 759-761 (SDD) is the key site enzyme activity of RdRp.

### Statistical analysis

All values are mean ± SD or SEM of individual samples. Data analysis was performed with GraphPad Prism Software (GraphPad Software Inc., version 6.01). The statistical tests utilized are two-tailed and respective details have been indicated in figure legends. p-value of < 0.05 were considered statistically significant. (*, p-value of ≤ 0.05. **, p-value of ≤ 0.005. ***, p-value of ≤ 0.0005. ****, p-value of ≤ 0.0001).

## ACKNOWLEDGEMENTS

The project was supported by Shenzhen Science and Technology Program (JSGG20200225150431472, ZDSYS20190902093215877 & KQTD20180411143323605), Shenzhen Bay Laboratory (SZBL2019062801006), Guangdong Basic and Applied Basic Research Foundation (Grant #2020A1515110361) and National Natural Science Foundation of China (grant #32041002 & #81620108020). D.G. is also supported by Guangdong Zhujiang Talents Program and National Ten-thousand Talents Program. We thank Dr. Chuwen Lin from School of Medicine, Sun Yat-Sen University for the help in lung pathology analysis. We thank the Center for Disease Control and Prevention of Guangdong Province for providing the Delta variant of SARS-CoV-2.

---

**Figure S1. Antiviral activity of 21 compounds in SARS-CoV-2 Replicon system.**

(A) HEK 293T cells transfected with SARS-CoV-2-Rep-Luci were treated with DMSO and 24 compounds with 10 μM. 60 h post-transfection, the cells were subjected to the Dual-Luciferase® Reporter (DLR™) Assay.

(B) The values of EC_50_ of each compound were analyzed.

**Figure S2. Antiviral activity of compounds against SARS-CoV-2 (B.1) in Huh7 cells.**

Huh7 cells were infected with B.1 original strain SARS-CoV-2 at an MOI of 0.05 and treated with dilutions of each compound (0, 0.01, 0.1, 0.5, 1, 2, 5 and 10 μM) for 48 h. Viral yield in the cultured supernatant was then quantified by qRT-PCR. The values of EC_50_ of each compound was analyzed.

**Figure S3. Cytotoxicity assay of compounds.**

Vero-E6 cells were plated in 96-well plate and treated with increasing concentrations compound ranging from 0 to 200 μM for 48 h. Cell viability was tested using Cell Counting Kit-8 (CCK-8). (A) ATV006. (B) other compounds.

**Figure S4. Genomic RNA (gRNA) and Subgenomic RNA (sgRNA) of Coronavirus.**

Detect the target position of the primer/probe sets of genomic RNA (gRNA) and subgenomic RNA (sgRNA) of Coronavirus. In this article, only FP and RP were used to detect sgRNA of SARS-CoV-2, and no Prb was used. The sequences of primer/probe sets are listed in Table S3.

**Figure S5. Dose-response in vivo anti-MHV efficacy of ATV006, remdesivir and 69-0 via intranasal inoculation.**

(A) Schematic of the experiment. Mouse were divided them into the following groups: vehicle (control), ATV006 (50, 20, 10, 5, 2 mg/kg, IG, once daily), remdesivir (20 mg/kg, IV, once daily) or 69-0 (50 mg/kg, IG, once daily). After MHV-A59 infects the mice, the administration is continued for 4 days, and the body weight curve (C) and survival curve (B) of the mice were recorded for 14 days. (D) Viral titers from livers were harvested at 3 dpi and analyzed by qRT-PCR.

**Figure S6. Dose-response anti-HMV efficacy of ATV006 via intrahepatic inoculation.**

(A) Schematic of the experiment. 5 weeks Balb/c mouse were intranasally intrahepatic with 1 × 10^5^ PFU per mouse of MHV-A59, and treated with vehicle (control), ATV006 (50,10, 2 mg/kg, IG, twice daily). (B, C and D) Viral titers from liver tissue were harvested at 3 dpi and analyzed by plaque assay and qRT-PCR. (E and F) ALT and AST analysis of serum from vehicle group and ATV006 (50 mg/kg, IG, twice daily). (G) Histopathology analysis of liver from vehicle group and ATV006 (50 mg/kg, IG, twice daily). (H) Representative chemokine and cytokine assessment of the liver tissues of the indicated groups, as detected in lung tissue homogenate at 3 dpi. *p values ≤ 0.05; **p values ≤ 0.005; ***p values ≤ 0.0005; * ***p values ≤ 0.0001.

**Figure S7. Analysis of RdRp mutation of SARS-CoV-2 and its variants.**

The black triangles represent the key sites where remdesivir binds to RdRp. The red triangles represent the enzyme activity amino acid residues of RdRp. The red dots represent the amino acid sites whose mutation rate is greater than one percent and less than eighty percent compared with the original strain. The blue dots represent the amino acid sites whose mutation rate is greater than eighty percent compared with the original strain.

**Figure S8. Coronavirus RdRp Conservation Analysis.**

Coronavirus RdRp amino acid sequence alignment, (A) Alpha-coronavirus and (B) beta-coronavirus. The black dots represent the key sites where remdesivir binds to RdRp. The red triangles represent the enzyme activity amino acid residues of RdRp.

**Table S1.** Permeability and efflux ratio determination of 69-0, ATV006, ATV019 and ATV020 in Caco-2 cells

| Compound | Caco-2 AB/BA (Papp (10<sup>-6</sup> cm/s)) <sup>a</sup> | Efflux ratio |
| --- | --- | --- |
| 69-0 | 1.22/1.20 | 0.98 |
| ATV006 | 0.51/0.87 | 1.7 |
| ATV019 | 0.28/0.68 | 2.47 |
| ATV020 | 0.17/0.22 | 1.28 |

<sup>a</sup> Papp (A to B) < 2, low permeability; 2 < Papp (A to B) < 10, moderate permeability; Papp (A to B) > 10, high permeability.

**Table S2.**
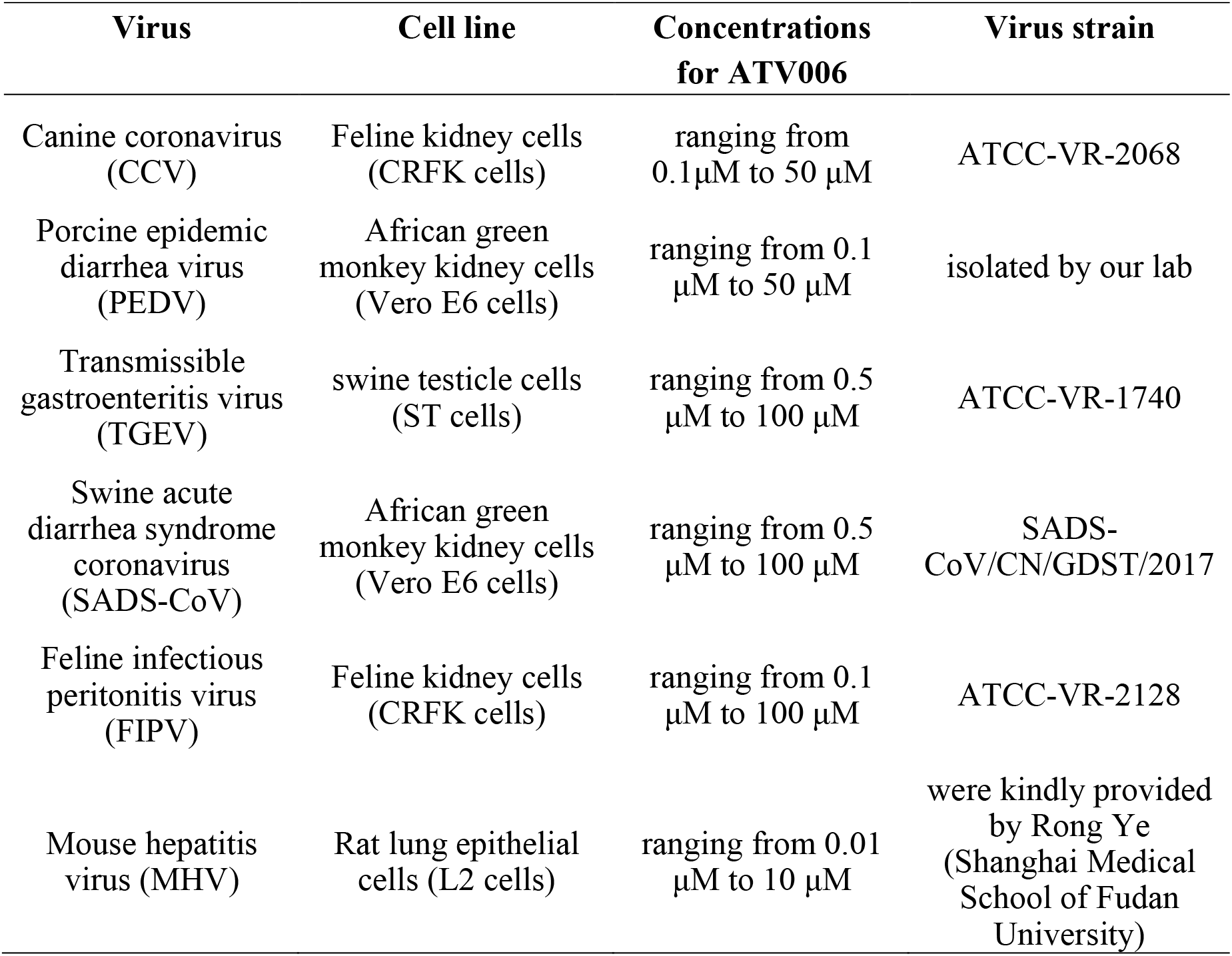
Animal coronaviruses and cells used in this study

**Table S3.**
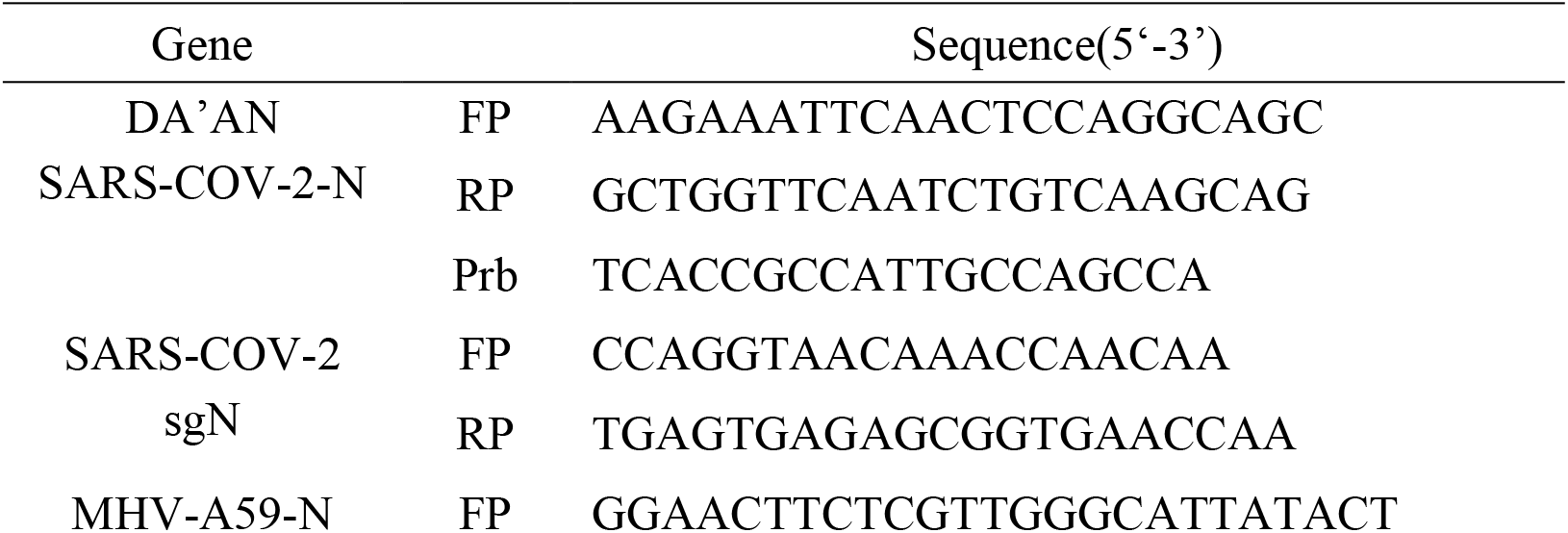
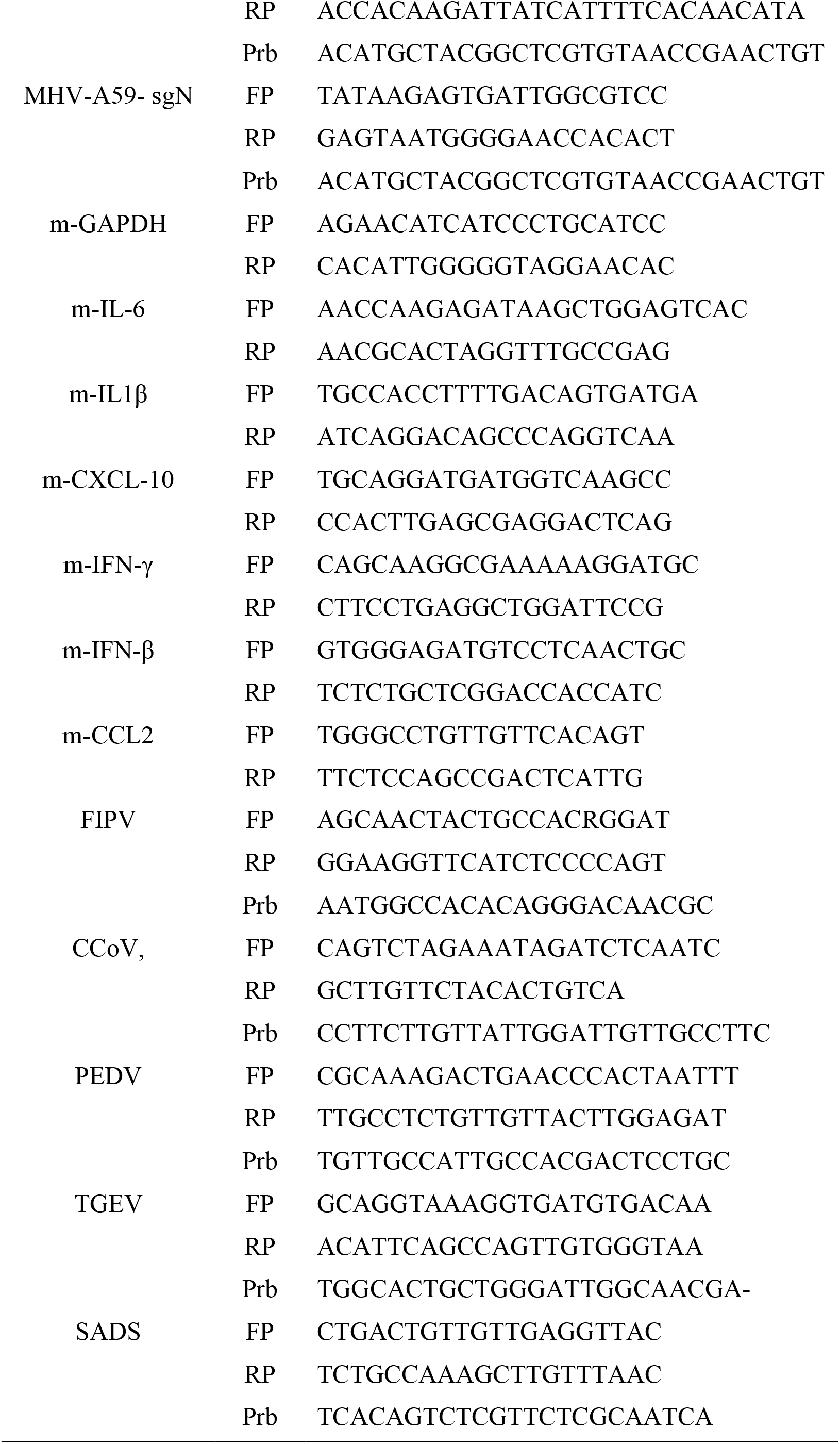
qPCR primers used for detection of viral genomes and various genes

**Table S4.**
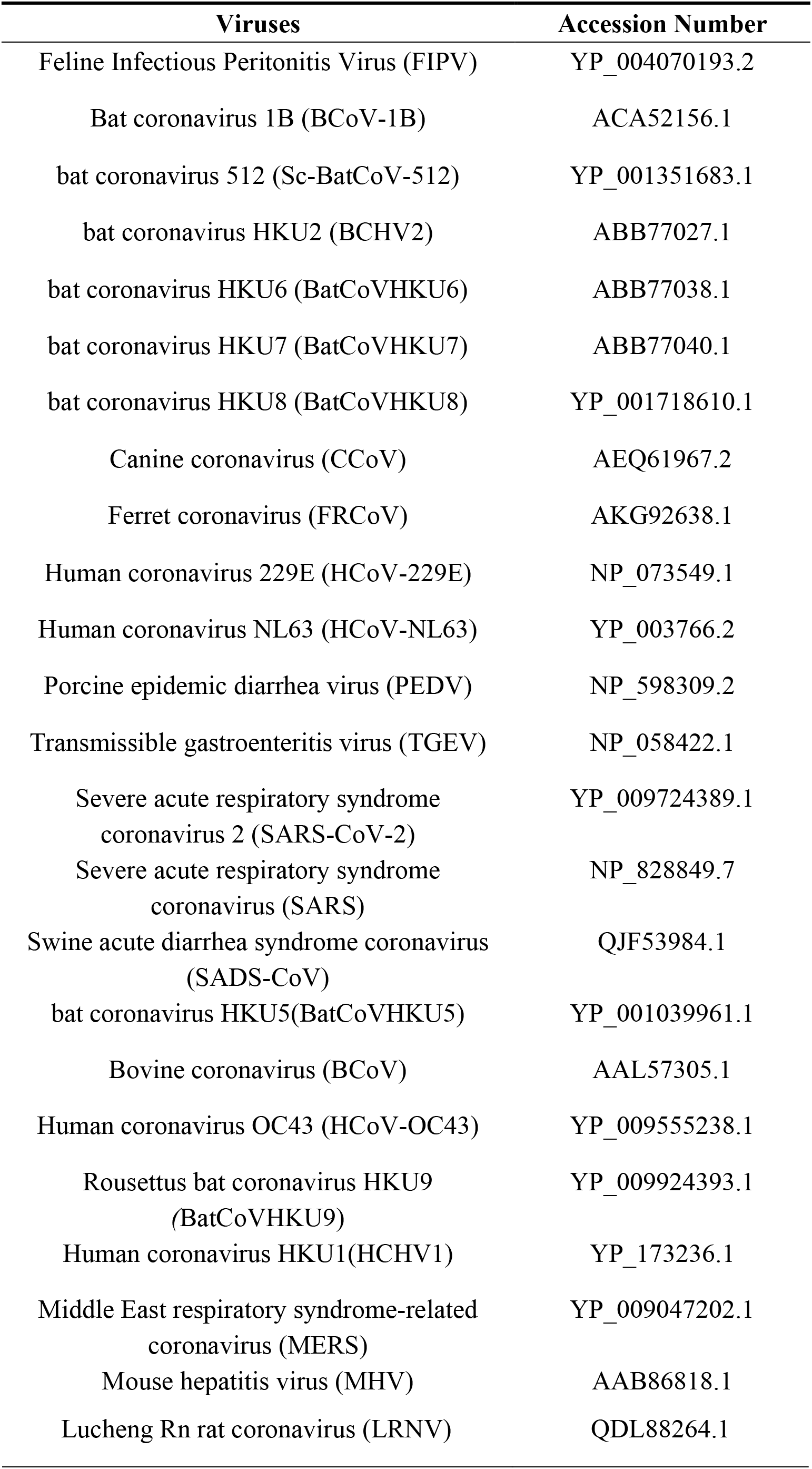
Viral genome sequences and accession numbers

## REFERENCES

Al-Tawfiq, J. A., and Z. A. Memish. 2014. ’Middle East respiratory syndrome coronavirus: transmission and phylogenetic evolution’, Trends Microbiol, 22: 573–9.

Alanazi, A. S., E. James, and Y. Mehellou. 2019. ’The ProTide Prodrug Technology: Where Next?’, ACS Med Chem Lett, 10: 2–5.

Ali., Farhan, Mohak Sharda., and Aswin Sai Narain Seshasayee. 2021. ’SARS-CoV-2 sequence typing, evolution and signatures of selection using CoVa, a Python-based command-line utility’, biovix.

Beigel, J. H., K. M. Tomashek, L. E. Dodd, A. K. Mehta, B. S. Zingman, A. C. Kalil, E. Hohmann, H. Y. Chu, A. Luetkemeyer, S. Kline, D. Lopez de Castilla, R. W. Finberg, K. Dierberg, V. Tapson, L. Hsieh, T. F. Patterson, R. Paredes, D. A. Sweeney, W. R. Short, G. Touloumi, D. C. Lye, N. Ohmagari, M. D. Oh, G. M. Ruiz-Palacios, T. Benfield, G. Fatkenheuer, M. G. Kortepeter, R. L. Atmar, C. B. Creech, J. Lundgren, A. G. Babiker, S. Pett, J. D. Neaton, T. H. Burgess, T. Bonnett, M. Green, M. Makowski, A. Osinusi, S. Nayak, H. C. Lane, and Actt-Study Group Members. 2020. ’Remdesivir for the Treatment of Covid-19 - Final Report’, N Engl J Med, 383: 1813–26.

Beitz, E. 2000. ’TEXshade: shading and labeling of multiple sequence alignments using LATEX2 epsilon’, Bioinformatics, 16: 135–9.

Chen, Y., Q. Liu, and D. Guo. 2020. ’Emerging coronaviruses: Genome structure, replication, and pathogenesis’, J Med Virol, 92: 2249.

Cho, A., O. L. Saunders, T. Butler, L. Zhang, J. Xu, J. E. Vela, J. Y. Feng, A. S. Ray, and C. U. Kim. 2012. ’Synthesis and antiviral activity of a series of 1’-substituted 4-aza-7,9-dideazaadenosine C-nucleosides’, Bioorg Med Chem Lett, 22: 2705–7.

Consortium, W. H. O. Solidarity Trial, H. Pan, R. Peto, A. M. Henao-Restrepo, M. P. Preziosi, V. Sathiyamoorthy, Q. Abdool Karim, M. M. Alejandria, C. Hernandez Garcia, M. P. Kieny, R. Malekzadeh, S. Murthy, K. S. Reddy, M. Roses Periago, P. Abi Hanna, F. Ader, A. M. Al-Bader, A. Alhasawi, E. Allum, A. Alotaibi, C. A. Alvarez-Moreno, S. Appadoo, A. Asiri, P. Aukrust, A. Barratt-Due, S. Bellani, M. Branca, H. B. C. Cappel-Porter, N. Cerrato, T. S. Chow, N. Como, J. Eustace, P. J. Garcia, S. Godbole, E. Gotuzzo, L. Griskevicius, R. Hamra, M. Hassan, M. Hassany, D. Hutton, I. Irmansyah, L. Jancoriene, J. Kirwan, S. Kumar, P. Lennon, G. Lopardo, P. Lydon, N. Magrini, T. Maguire, S. Manevska, O. Manuel, S. McGinty, M. T. Medina, M. L. Mesa Rubio, M. C. Miranda-Montoya, J. Nel, E. P. Nunes, M. Perola, A. Portoles, M. R. Rasmin, A. Raza, H. Rees, P. P. S. Reges, C. A. Rogers, K. Salami, M. I. Salvadori, N. Sinani, J. A. C. Sterne, M. Stevanovikj, E. Tacconelli, K. A. O. Tikkinen, S. Trelle, H. Zaid, J. A. Rottingen, and S. Swaminathan. 2021. ’Repurposed Antiviral Drugs for Covid-19 - Interim WHO Solidarity Trial Results’, N Engl J Med, 384: 497–511.

Faheem, B. K. Kumar, Kvgc Sekhar, S. Kunjiappan, J. Jamalis, R. Balaña-Fouce, B. L. Tekwani, and M. Sankaranarayanan. 2020. ’Druggable targets of SARS-CoV-2 and treatment opportunities for COVID-19’, Bioorg Chem, 104: 104269.

Fischer, W., J. J. Eron, W. Holman, M. S. Cohen, L. Fang, L. J. Szewczyk, T. P. Sheahan, R. Baric, K. R. Mollan, C. R. Wolfe, E. R. Duke, M. M. Azizad, K. Borroto-Esoda, D. A. Wohl, A. J. Loftis, P. Alabanza, F. Lipansky, and W. P. Painter. 2021. ’Molnupiravir, an Oral Antiviral Treatment for COVID-19’, medRxiv.

Fung, T. S., and D. X. Liu. 2019. ’Human Coronavirus: Host-Pathogen Interaction’, Annu Rev Microbiol, 73: 529–57.

Ghasemnejad-Berenji, M., and S. Pashapour. 2021. ’Favipiravir and COVID-19: A Simplified Summary’, Drug Res (Stuttg), 71: 166–70.

Goldman, J. D., D. C. B. Lye, D. S. Hui, K. M. Marks, R. Bruno, R. Montejano, C. D. Spinner, M. Galli, M. Y. Ahn, R. G. Nahass, Y. S. Chen, D. SenGupta, R. H. Hyland, A. O. Osinusi, H. Cao, C. Blair, X. Wei, A. Gaggar, D. M. Brainard, W. J. Towner, J. Munoz, K. M. Mullane, F. M. Marty, K. T. Tashima, G. Diaz, A. Subramanian, and Gs-Us-Investigators. 2020. ’Remdesivir for 5 or 10 Days in Patients with Severe Covid-19’, N Engl J Med, 383: 1827–37.

Good, S. S., J. Westover, K. H. Jung, X. J. Zhou, A. Moussa, P. La Colla, G. Collu, B. Canard, and J. P. Sommadossi. 2021. ’AT-527, a Double Prodrug of a Guanosine Nucleotide Analog, Is a Potent Inhibitor of SARS-CoV-2 In Vitro and a Promising Oral Antiviral for Treatment of COVID-19’, Antimicrob Agents Chemother, 65.

Haake, C., S. Cook, N. Pusterla, and B. Murphy. 2020. ’Coronavirus Infections in Companion Animals: Virology, Epidemiology, Clinical and Pathologic Features’, Viruses, 12.

Harvey, W. T., A. M. Carabelli, B. Jackson, R. K. Gupta, E. C. Thomson, E. M. Harrison, C. Ludden, R. Reeve, A. Rambaut, Covid-Genomics UK Consortium, S. J. Peacock, and D. L. Robertson. 2021. ’SARS-CoV-2 variants, spike mutations and immune escape’, Nat Rev Microbiol, 19: 409–24.

Hillen, H. S., G. Kokic, L. Farnung, C. Dienemann, D. Tegunov, and P. Cramer. 2020. ’Structure of replicating SARS-CoV-2 polymerase’, Nature, 584: 154–56.

Hsu, C. H., M. Jay, P. M. Bummer, and H. J. Lehmler. 2003. ’Chemical stability of esters of nicotinic acid intended for pulmonary administration by liquid ventilation’, Pharm Res, 20: 918–25.

Huang, Z., L. Gong, Z. Zheng, Q. Gao, X. Chen, Y. Chen, X. Chen, R. Xu, J. Zheng, Z. Xu, S. Zhang, H. Wang, and G. Zhang. 2021. ’GS-441524 inhibits African swine fever virus infection in vitro’, Antiviral Res, 191: 105081.

Hussain, S., J. Pan, Y. Chen, Y. Yang, J. Xu, Y. Peng, Y. Wu, Z. Li, Y. Zhu, P. Tien, and D. Guo. 2005. ’Identification of novel subgenomic RNAs and noncanonical transcription initiation signals of severe acute respiratory syndrome coronavirus’, J Virol, 79: 5288–95.

Izes, A. M., J. Yu, J. M. Norris, and M. Govendir. 2020. ’Current status on treatment options for feline infectious peritonitis and SARS-CoV-2 positive cats’, Vet Q, 40: 322–30.

Jin, Y., H. Lin, L. Cao, W. C. Wu, Y. Ji, L. Du, Y. Jiang, Y. Xie, K. Tong, F. Xing, F. Zheng, M. Shi, J. A. Pan, X. Peng, and D. Guo. 2021. ’A Convenient and Biosafe Replicon with Accessory Genes of SARS-CoV-2 and Its Potential Application in Antiviral Drug Discovery’, Virol Sin.

Kabinger, F., C. Stiller, J. Schmitzova, C. Dienemann, G. Kokic, H. S. Hillen, C. Hobartner, and P. Cramer. 2021. ’Mechanism of molnupiravir-induced SARS-CoV-2 mutagenesis’, Nat Struct Mol Biol.

Katoh, K., K. Misawa, K. Kuma, and T. Miyata. 2002. ’MAFFT: a novel method for rapid multiple sequence alignment based on fast Fourier transform’, Nucleic Acids Res, 30: 3059–66.

Kim, D., J. Y. Lee, J. S. Yang, J. W. Kim, V. N. Kim, and H. Chang. 2020. ’The Architecture of SARS-CoV-2 Transcriptome’, Cell, 181: 914–21 e10.

Kokic, G., H. S. Hillen, D. Tegunov, C. Dienemann, F. Seitz, J. Schmitzova, L. Farnung, A. Siewert, C. Hobartner, and P. Cramer. 2021. ’Mechanism of SARS-CoV-2 polymerase stalling by remdesivir’, Nat Commun, 12: 279.

Korner, R. W., M. Majjouti, M. A. A. Alcazar, and E. Mahabir. 2020. ’Of Mice and Men: The Coronavirus MHV and Mouse Models as a Translational Approach to Understand SARS-CoV-2’, Viruses, 12.

Laude, H., D. Rasschaert, B. Delmas, M. Godet, J. Gelfi, and B. Charley. 1990. ’Molecular biology of transmissible gastroenteritis virus’, Vet Microbiol, 23: 147–54.

Lavis, L. D. 2008. ’Ester bonds in prodrugs’, ACS Chem Biol, 3: 203–6.

Lee, C. 2015. ’Porcine epidemic diarrhea virus: An emerging and re-emerging epizootic swine virus’, Virol J, 12: 193.

Li, Y., L. Cao, G. Li, F. Cong, Y. Li, J. Sun, Y. Luo, G. Chen, G. Li, P. Wang, F. Xing, Y. Ji, J. Zhao, Y. Zhang, D. Guo, and X. Zhang. 2021. ’Remdesivir Metabolite GS-441524 Effectively Inhibits SARS-CoV-2 Infection in Mouse Models’, J Med Chem.

Liu, C., H. M. Ginn, W. Dejnirattisai, P. Supasa, B. Wang, A. Tuekprakhon, R. Nutalai, D. Zhou, A. J. Mentzer, Y. Zhao, H. M. E. Duyvesteyn, C. Lopez-Camacho, J. Slon-Campos, T. S. Walter, D. Skelly, S. A. Johnson, T. G. Ritter, C. Mason, S. A. Costa Clemens, F. Gomes Naveca, V. Nascimento, F. Nascimento, C. Fernandes da Costa, P. C. Resende, A. Pauvolid-Correa, M. M. Siqueira, C. Dold, N. Temperton, T. Dong, A. J. Pollard, J. C. Knight, D. Crook, T. Lambe, E. Clutterbuck, S. Bibi, A. Flaxman, M. Bittaye, S. Belij-Rammerstorfer, S. C. Gilbert, T. Malik, M. W. Carroll, P. Klenerman, E. Barnes, S. J. Dunachie, V. Baillie, N. Serafin, Z. Ditse, K. Da Silva, N. G. Paterson, M. A. Williams, D. R. Hall, S. Madhi, M. C. Nunes, P. Goulder, E. E. Fry, J. Mongkolsapaya, J. Ren, D. I. Stuart, and G. R. Screaton. 2021. ’Reduced neutralization of SARS-CoV-2 B.1.617 by vaccine and convalescent serum’, Cell, 184: 4220–36 e13.

Lo, M. K., R. Jordan, A. Arvey, J. Sudhamsu, P. Shrivastava-Ranjan, A. L. Hotard, M. Flint, L. K. McMullan, D. Siegel, M. O. Clarke, R. L. Mackman, H. C. Hui, M. Perron, A. S. Ray, T. Cihlar, S. T. Nichol, and C. F. Spiropoulou. 2017. ’GS-5734 and its parent nucleoside analog inhibit Filo-, Pneumo-, and Paramyxoviruses’, Sci Rep, 7: 43395.

Mackman, R. L., H. C. Hui, M. Perron, E. Murakami, C. Palmiotti, G. Lee, K. Stray, L. Zhang, B. Goyal, K. Chun, D. Byun, D. Siegel, S. Simonovich, V. Du Pont, J. Pitts, D. Babusis, A. Vijjapurapu, X. Lu, C. Kim, X. Zhao, J. Chan, B. Ma, D. Lye, A. Vandersteen, S. Wortman, K. T. Barrett, M. Toteva, R. Jordan, R. Subramanian, J. P. Bilello, and T. Cihlar. 2021. ’Prodrugs of a 1’-CN-4-Aza-7,9-dideazaadenosine C-Nucleoside Leading to the Discovery of Remdesivir (GS-5734) as a Potent Inhibitor of Respiratory Syncytial Virus with Efficacy in the African Green Monkey Model of RSV’, J Med Chem, 64: 5001–17.

Mari, A., T. Roloff, M. Stange, K. K. Sogaard, E. Asllanaj, G. Tauriello, L. T. Alexander, M. Schweitzer, K. Leuzinger, A. Gensch, A. E. Martinez, J. Bielicki, H. Pargger, M. Siegemund, C. H. Nickel, R. Bingisser, M. Osthoff, S. Bassetti, P. Sendi, M. Battegay, C. Marzolini, H. M. B. Seth-Smith, T. Schwede, H. H. Hirsch, and A. Egli. 2021. ’Global Genomic Analysis of SARS-CoV-2 RNA Dependent RNA Polymerase Evolution and Antiviral Drug Resistance’, Microorganisms, 9.

Martin, Y. C. 2005. ’A bioavailability score’, J Med Chem, 48: 3164–70.

McCallum, M., J. Bassi, A. Marco, A. Chen, A. C. Walls, J. D. Iulio, M. A. Tortorici, M. J. Navarro, C. Silacci-Fregni, C. Saliba, M. Agostini, D. Pinto, K. Culap, S. Bianchi, S. Jaconi, E. Cameroni, J. E. Bowen, S. W. Tilles, M. S. Pizzuto, S. B. Guastalla, G. Bona, A. F. Pellanda, C. Garzoni, W. C. Van Voorhis, L. E. Rosen, G. Snell, A. Telenti, H. W. Virgin, L. Piccoli, D. Corti, and D. Veesler. 2021. ’SARS-CoV-2 immune evasion by variant B.1.427/B.1.429’, bioRxiv.

Minghua, Li., Ferretti. Max, Ying. Baoling, Descamps. Hélène, Lee. Emily, Dittmar. Mark, Lee. Jae Seung, Whig. Kanupriya, Brinda Kamalia., Dohnalová. Lenka, Uhr. Giulia, Zarkoob. Hoda, Chen. Yu-Chi, Ramage. Holly, Ferrer. Marc, Lynch. Kristen, Schultz. David C., Christoph A. Thaiss., Diamond. Michael S., and Cherry. Sara. 2021. ’Pharmacological activation of STING blocks SARS-CoV-2 infection’, Sci Immunol, 6.

Motozono, C., M. Toyoda, J. Zahradnik, A. Saito, H. Nasser, T. S. Tan, I. Ngare, I. Kimura, K. Uriu, Y. Kosugi, Y. Yue, R. Shimizu, J. Ito, S. Torii, A. Yonekawa, N. Shimono, Y. Nagasaki, R. Minami, T. Toya, N. Sekiya, T. Fukuhara, Y. Matsuura, G. Schreiber, Consortium Genotype to Phenotype Japan, T. Ikeda, S. Nakagawa, T. Ueno, and K. Sato. 2021. ’SARS-CoV-2 spike L452R variant evades cellular immunity and increases infectivity’, Cell Host Microbe, 29: 1124–36 e11.

Murphy, B. G., M. Perron, E. Murakami, K. Bauer, Y. Park, C. Eckstrand, M. Liepnieks, and N. C. Pedersen. 2018. ’The nucleoside analog GS-441524 strongly inhibits feline infectious peritonitis (FIP) virus in tissue culture and experimental cat infection studies’, Vet Microbiol, 219: 226–33.

NCATS. 2021. ’GS-441524 Studies’, National Center for Advancing Translational Sciences. https://opendata.ncats.nih.gov/covid19/GS-441524.

Oladunni, F. S., J. G. Park, P. A. Pino, O. Gonzalez, A. Akhter, A. Allue-Guardia, A. Olmo-Fontanez, S. Gautam, A. Garcia-Vilanova, C. Ye, K. Chiem, C. Headley, V. Dwivedi, L. M. Parodi, K. J. Alfson, H. M. Staples, A. Schami, J. I. Garcia, A. Whigham, R. N. Platt, 2nd, M. Gazi, J. Martinez, C. Chuba, S. Earley, O. H. Rodriguez, S. D. Mdaki, K. N. Kavelish, R. Escalona, C. R. A. Hallam, C. Christie, J. L. Patterson, T. J. C. Anderson, R. Carrion, Jr., E. J. Dick, Jr., S. Hall-Ursone, L. S. Schlesinger, X. Alvarez, D. Kaushal, L. D. Giavedoni, J. Turner, L. Martinez-Sobrido, and J. B. Torrelles. 2020. ’Lethality of SARS-CoV-2 infection in K18 human angiotensin-converting enzyme 2 transgenic mice’, Nat Commun, 11: 6122.

Pedersen, N. C., M. Perron, M. Bannasch, E. Montgomery, E. Murakami, M. Liepnieks, and H. Liu. 2019. ’Efficacy and safety of the nucleoside analog GS-441524 for treatment of cats with naturally occurring feline infectious peritonitis’, J Feline Med Surg, 21: 271–81.

Perlman, S., and J. Netland. 2009. ’Coronaviruses post-SARS: update on replication and pathogenesis’, Nat Rev Microbiol, 7: 439–50.

Picarazzi, F., I. Vicenti, F. Saladini, M. Zazzi, and M. Mori. 2020. ’Targeting the RdRp of Emerging RNA Viruses: The Structure-Based Drug Design Challenge’, Molecules, 25.

Pruijssers, A. J., A. S. George, A. Schafer, S. R. Leist, L. E. Gralinksi, K. H. Dinnon, 3rd, B. L. Yount, M. L. Agostini, L. J. Stevens, J. D. Chappell, X. Lu, T. M. Hughes, K. Gully, D. R. Martinez, A. J. Brown, R. L. Graham, J. K. Perry, V. Du Pont, J. Pitts, B. Ma, D. Babusis, E. Murakami, J. Y. Feng, J. P. Bilello, D. P. Porter, T. Cihlar, R. S. Baric, M. R. Denison, and T. P. Sheahan. 2020. ’Remdesivir Inhibits SARS-CoV-2 in Human Lung Cells and Chimeric SARS-CoV Expressing the SARS-CoV-2 RNA Polymerase in Mice’, Cell Rep, 32: 107940.

Reardon, S. 2021. ’How the Delta variant achieves its ultrafast spread’, Nature.

Robson, F., K. S. Khan, T. K. Le, C. Paris, S. Demirbag, P. Barfuss, P. Rocchi, and W. L. Ng. 2020. ’Coronavirus RNA Proofreading: Molecular Basis and Therapeutic Targeting’, Mol Cell, 79: 710–27.

Sabbah, D. A., R. Hajjo, S. K. Bardaweel, and H. A. Zhong. 2021. ’An Updated Review on SARS-CoV-2 Main Proteinase (M(Pro)): Protein Structure and Small-Molecule Inhibitors’, Curr Top Med Chem, 21: 442–60.

Salleh, M. Z., J. P. Derrick, and Z. Z. Deris. 2021. ’Structural Evaluation of the Spike Glycoprotein Variants on SARS-CoV-2 Transmission and Immune Evasion’, Int J Mol Sci, 22.

Shu, Y., and J. McCauley. 2017. ’GISAID: Global initiative on sharing all influenza data - from vision to reality’, Euro Surveill, 22.

Sun, J., Z. Zhuang, J. Zheng, K. Li, R. L. Wong, D. Liu, J. Huang, J. He, A. Zhu, J. Zhao, X. Li, Y. Xi, R. Chen, A. N. Alshukairi, Z. Chen, Z. Zhang, C. Chen, X. Huang, F. Li, X. Lai, D. Chen, L. Wen, J. Zhuo, Y. Zhang, Y. Wang, S. Huang, J. Dai, Y. Shi, K. Zheng, M. R. Leidinger, J. Chen, Y. Li, N. Zhong, D. K. Meyerholz, P. B. McCray, Jr., S. Perlman, and J. Zhao. 2020a. ’Generation of a Broadly Useful Model for COVID-19 Pathogenesis, Vaccination, and Treatment’, Cell, 182: 734–43 e5.

Sun, S. H., Q. Chen, H. J. Gu, G. Yang, Y. X. Wang, X. Y. Huang, S. S. Liu, N. N. Zhang, X. F. Li, R. Xiong, Y. Guo, Y. Q. Deng, W. J. Huang, Q. Liu, Q. M. Liu, Y. L. Shen, Y. Zhou, X. Yang, T. Y. Zhao, C. F. Fan, Y. S. Zhou, C. F. Qin, and Y. C. Wang. 2020b. ’A Mouse Model of SARS-CoV-2 Infection and Pathogenesis’, Cell Host Microbe, 28: 124–33 e4.

Tegally, H., E. Wilkinson, M. Giovanetti, A. Iranzadeh, V. Fonseca, J. Giandhari, D. Doolabh, S. Pillay, E. J. San, N. Msomi, K. Mlisana, A. von Gottberg, S. Walaza, M. Allam, A. Ismail, T. Mohale, A. J. Glass, S. Engelbrecht, G. Van Zyl, W. Preiser, F. Petruccione, A. Sigal, D. Hardie, G. Marais, N. Y. Hsiao, S. Korsman, M. A. Davies, L. Tyers, I. Mudau, D. York, C. Maslo, D. Goedhals, S. Abrahams, O. Laguda-Akingba, A. Alisoltani-Dehkordi, A. Godzik, C. K. Wibmer, B. T. Sewell, J. Lourenco, L. C. J. Alcantara, S. L. Kosakovsky Pond, S. Weaver, D. Martin, R. J. Lessells, J. N. Bhiman, C. Williamson, and T. de Oliveira. 2021. ’Detection of a SARS-CoV-2 variant of concern in South Africa’, Nature, 592: 438–43.

Teyssou, E., H. Delagreverie, B. Visseaux, S. Lambert-Niclot, S. Brichler, V. Ferre, S. Marot, A. Jary, E. Todesco, A. Schnuriger, E. Ghidaoui, B. Abdi, S. Akhavan, N. Houhou-Fidouh, C. Charpentier, L. Morand-Joubert, D. Boutolleau, D. Descamps, V. Calvez, A. G. Marcelin, and C. Soulie. 2021. ’The Delta SARS-CoV-2 variant has a higher viral load than the Beta and the historical variants in nasopharyngeal samples from newly diagnosed COVID-19 patients’, J Infect.

Vuong, W., C. Fischer, M. B. Khan, M. J. van Belkum, T. Lamer, K. D. Willoughby, J. Lu, E. Arutyunova, M. A. Joyce, H. A. Saffran, J. A. Shields, H. S. Young, J. A. Nieman, D. L. Tyrrell, M. J. Lemieux, and J. C. Vederas. 2021. ’Improved SARS-CoV-2 M(pro) inhibitors based on feline antiviral drug GC376: Structural enhancements, increased solubility, and micellar studies’, Eur J Med Chem, 222: 113584.

Wahl, A., L. E. Gralinski, C. E. Johnson, W. Yao, M. Kovarova, K. H. Dinnon, 3rd, H. Liu, V. J. Madden, H. M. Krzystek, C. De, K. K. White, K. Gully, A. Schafer, T. Zaman, S. R. Leist, P. O. Grant, G. R. Bluemling, A. A. Kolykhalov, M. G. Natchus, F. B. Askin, G. Painter, E. P. Browne, C. D. Jones, R. J. Pickles, R. S. Baric, and J. V. Garcia. 2021. ’SARS-CoV-2 infection is effectively treated and prevented by EIDD-2801’, Nature, 591: 451–57.

Wang, M., R. Cao, L. Zhang, X. Yang, J. Liu, M. Xu, Z. Shi, Z. Hu, W. Zhong, and G. Xiao. 2020a. ’Remdesivir and chloroquine effectively inhibit the recently emerged novel coronavirus (2019-nCoV) in vitro’, Cell Res, 30: 269–71.

Wang, Q., J. Wu, H. Wang, Y. Gao, Q. Liu, A. Mu, W. Ji, L. Yan, Y. Zhu, C. Zhu, X. Fang, X. Yang, Y. Huang, H. Gao, F. Liu, J. Ge, Q. Sun, X. Yang, W. Xu, Z. Liu, H. Yang, Z. Lou, B. Jiang, L. W. Guddat, P. Gong, and Z. Rao. 2020b. ’Structural Basis for RNA Replication by the SARS-CoV-2 Polymerase’, Cell, 182: 417–28 e13.

Wang, Y., R. Chen, F. Hu, Y. Lan, Z. Yang, C. Zhan, J. Shi, X. Deng, M. Jiang, S. Zhong, B. Liao, K. Deng, J. Tang, L. Guo, M. Jiang, Q. Fan, M. Li, J. Liu, Y. Shi, X. Deng, X. Xiao, M. Kang, Y. Li, W. Guan, Y. Li, S. Li, F. Li, N. Zhong, and X. Tang. 2021. ’Transmission, viral kinetics and clinical characteristics of the emergent SARS-CoV-2 Delta VOC in Guangzhou, China’, EClinicalMedicine, 40: 101129.

Wei, D., T. Hu, Y. Zhang, W. Zheng, H. Xue, J. Shen, Y. Xie, and H. A. Aisa. 2021. ’Potency and pharmacokinetics of GS-441524 derivatives against SARS-CoV-2’, Bioorg Med Chem, 46: 116364.

Weiss, S. R., and J. L. Leibowitz. 2011. ’Coronavirus pathogenesis’, Adv Virus Res, 81: 85–164.

WHO. 2021a. ’SARS-CoV-2 Variants, Working Definitions and Actions Taken’. https://www.who.int/en/activities/tracking-SARS-CoV-2-variants

WHO. 2021b. ’WHO Coronavirus (COVID-19) Dashboard’. https://covid19.who.int.

Wu, F., S. Zhao, B. Yu, Y. M. Chen, W. Wang, Z. G. Song, Y. Hu, Z. W. Tao, J. H. Tian, Y. Y. Pei, M. L. Yuan, Y. L. Zhang, F. H. Dai, Y. Liu, Q. M. Wang, J. J. Zheng, L. Xu, E. C. Holmes, and Y. Z. Zhang. 2020. ’A new coronavirus associated with human respiratory disease in China’, Nature, 579: 265–69.

Yin, W., X. Luan, Z. Li, Y. Xie, Z. Zhou, J. Liu, M. Gao, X. Wang, F. Zhou, Q. Wang, Q. Wang, D. Shen, Y. Zhang, G. Tian, Haji A. Aisa, T. Hu, D. Wei, Y. Jiang, G. Xiao, H. Jiang, L. Zhang, X. Yu, J. Shen, S. Zhang, and H. E. Xu. 2020a. ’Structural basis for repurpose and design of nucleoside drugs for treating COVID-19’.

Yin, W., C. Mao, X. Luan, D. D. Shen, Q. Shen, H. Su, X. Wang, F. Zhou, W. Zhao, M. Gao, S. Chang, Y. C. Xie, G. Tian, H. W. Jiang, S. C. Tao, J. Shen, Y. Jiang, H. Jiang, Y. Xu, S. Zhang, Y. Zhang, and H. E. Xu. 2020b. ’Structural basis for inhibition of the RNA-dependent RNA polymerase from SARS-CoV-2 by remdesivir’, Science, 368: 1499–504.

Zhang, Q., R. Xiang, S. Huo, Y. Zhou, S. Jiang, Q. Wang, and F. Yu. 2021. ’Molecular mechanism of interaction between SARS-CoV-2 and host cells and interventional therapy’, Signal Transduct Target Ther, 6: 233.

Zhang, Y., J. Sun, Y. Sun, Y. Wang, and Z. He. 2013. ’Prodrug design targeting intestinal PepT1 for improved oral absorption: design and performance’, Curr Drug Metab, 14: 675–87.

Zhou, P., H. Fan, T. Lan, X. L. Yang, W. F. Shi, W. Zhang, Y. Zhu, Y. W. Zhang, Q. M. Xie, S. Mani, X. S. Zheng, B. Li, J. M. Li, H. Guo, G. Q. Pei, X. P. An, J. W. Chen, L. Zhou, K. J. Mai, Z. X. Wu, D. Li, D. E. Anderson, L. B. Zhang, S. Y. Li, Z. Q. Mi, T. T. He, F. Cong, P. J. Guo, R. Huang, Y. Luo, X. L. Liu, J. Chen, Y. Huang, Q. Sun, X. L. Zhang, Y. Y. Wang, S. Z. Xing, Y. S. Chen, Y. Sun, J. Li, P. Daszak, L. F. Wang, Z. L. Shi, Y. G. Tong, and J. Y. Ma. 2018. ’Fatal swine acute diarrhoea syndrome caused by an HKU2-related coronavirus of bat origin’, Nature, 556: 255–58.

