## Supplemental figures for "The adenosine analogue prodrug ATV006 is orally bioavailable and has potent preclinical efficacy against SARS-CoV-2 and its variants"

Figure S1. Antiviral activities of the 21 compounds in SARS-CoV-2 replicon system.

A

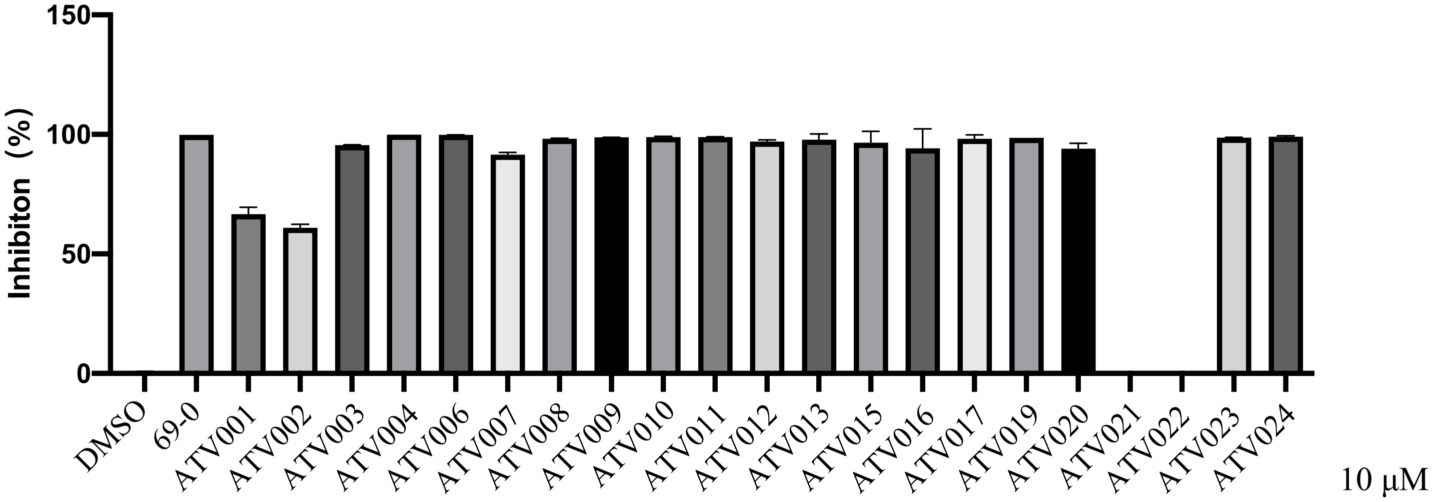


B

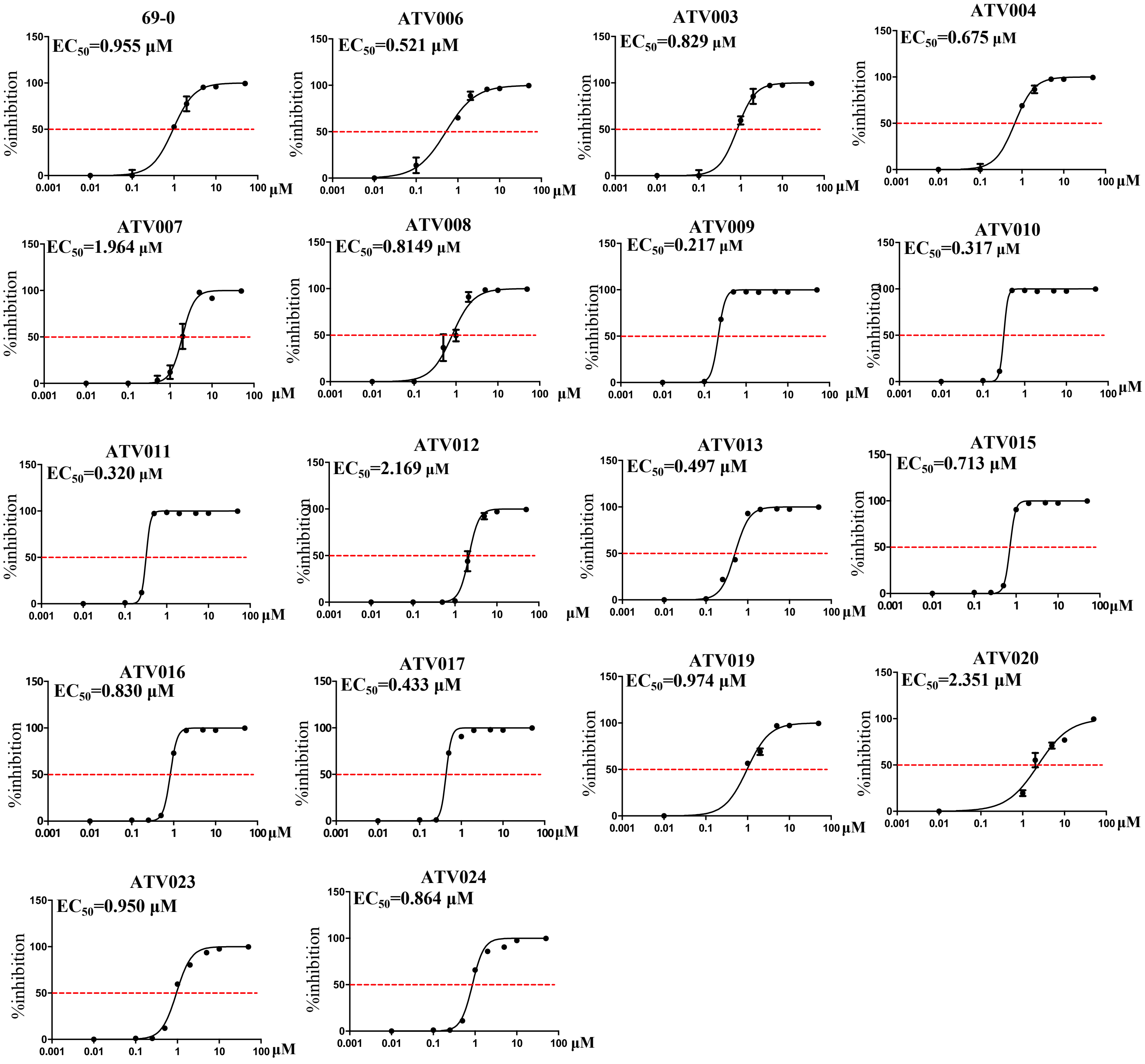

#### Figure S2. Antiviral activities of compounds against SARS-CoV-2 (B.1) in Huh7 cells.

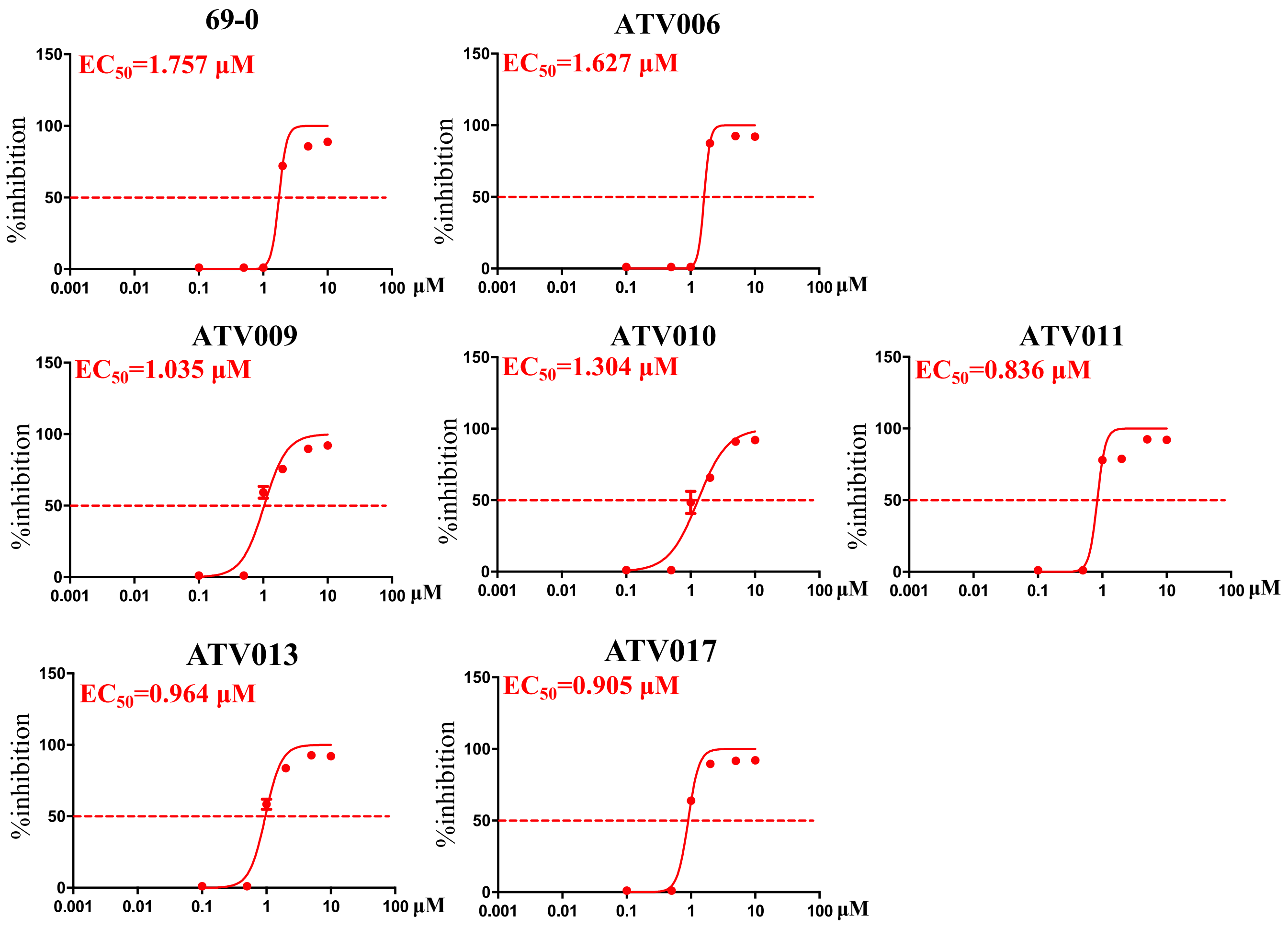

### Figure S3. Cytotoxicity assay of compounds in Vero E6 cells.

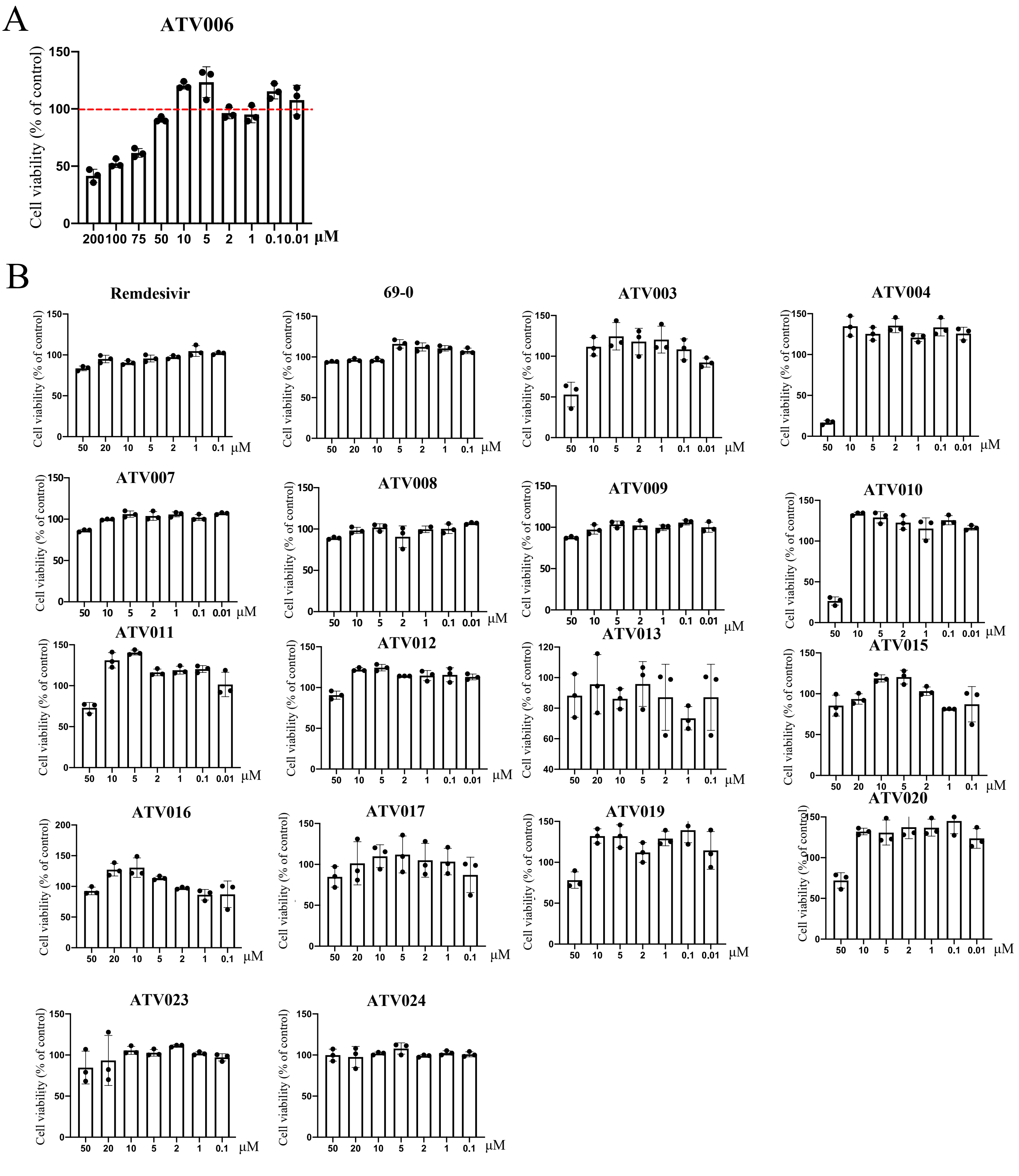


### Figure S4. Genomic RNA (gRNA) and subgenomic RNA (sgRNA) of Coronavirus.

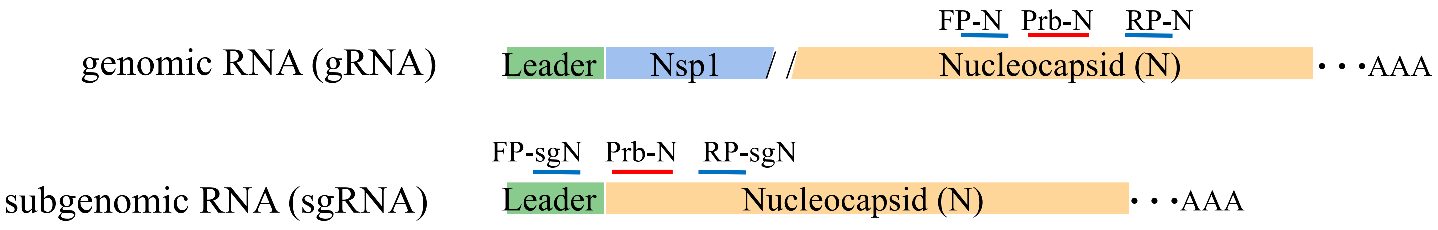

**Figure S5. Dose-response *in vivo* anti-MHV efficacy of ATV006, remdesivir and 69-0 via intranasal inoculation.**

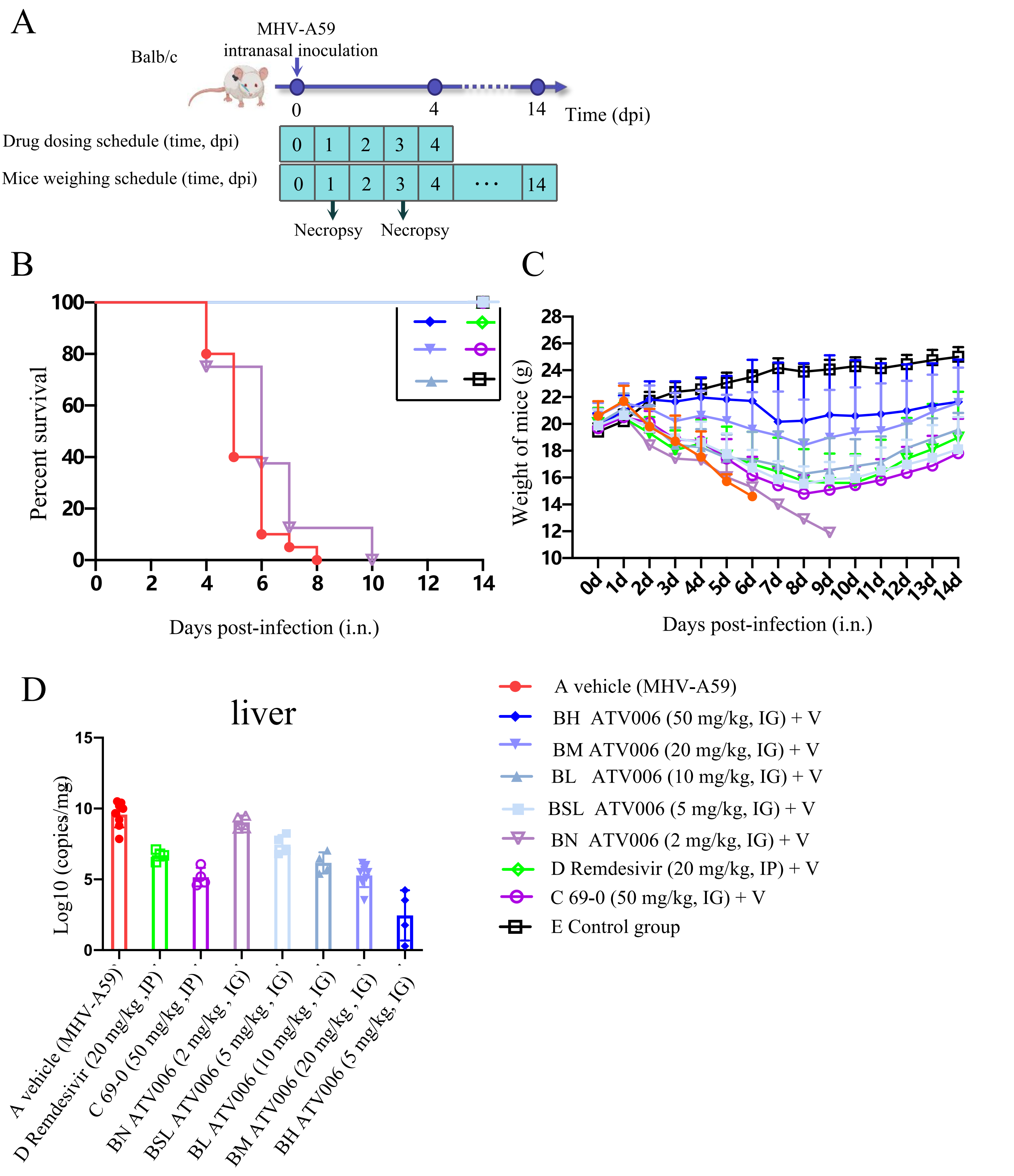

**Figure S6. Dose-response anti-MHV efficacy of ATV006 via intrahepatic inoculation.**

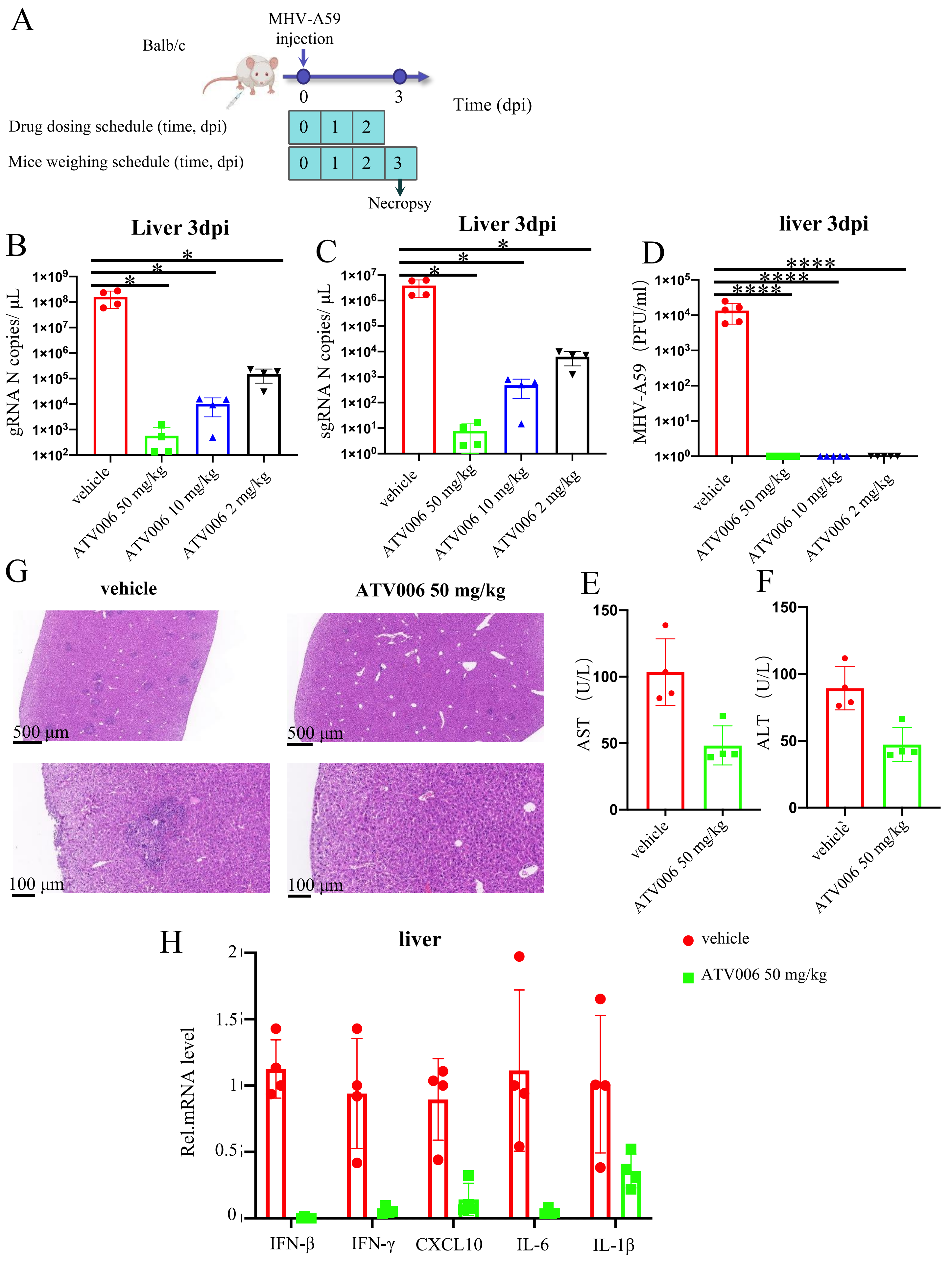

### Figure S7. Analysis of RdRp mutation of SARS-CoV-2 and its variants.

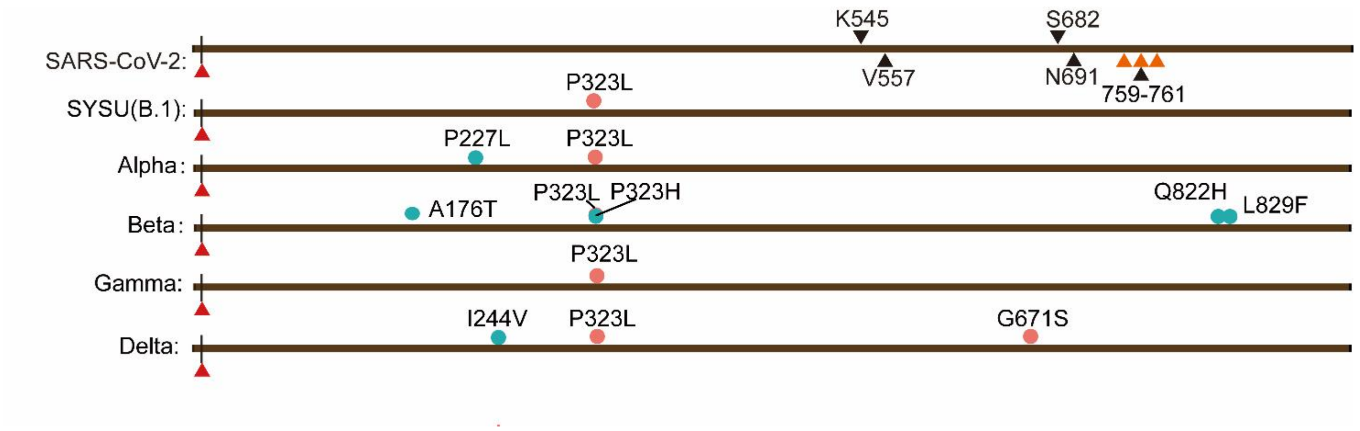

### Figure S8. Coronavirus RdRP Conservation Analysis.

A

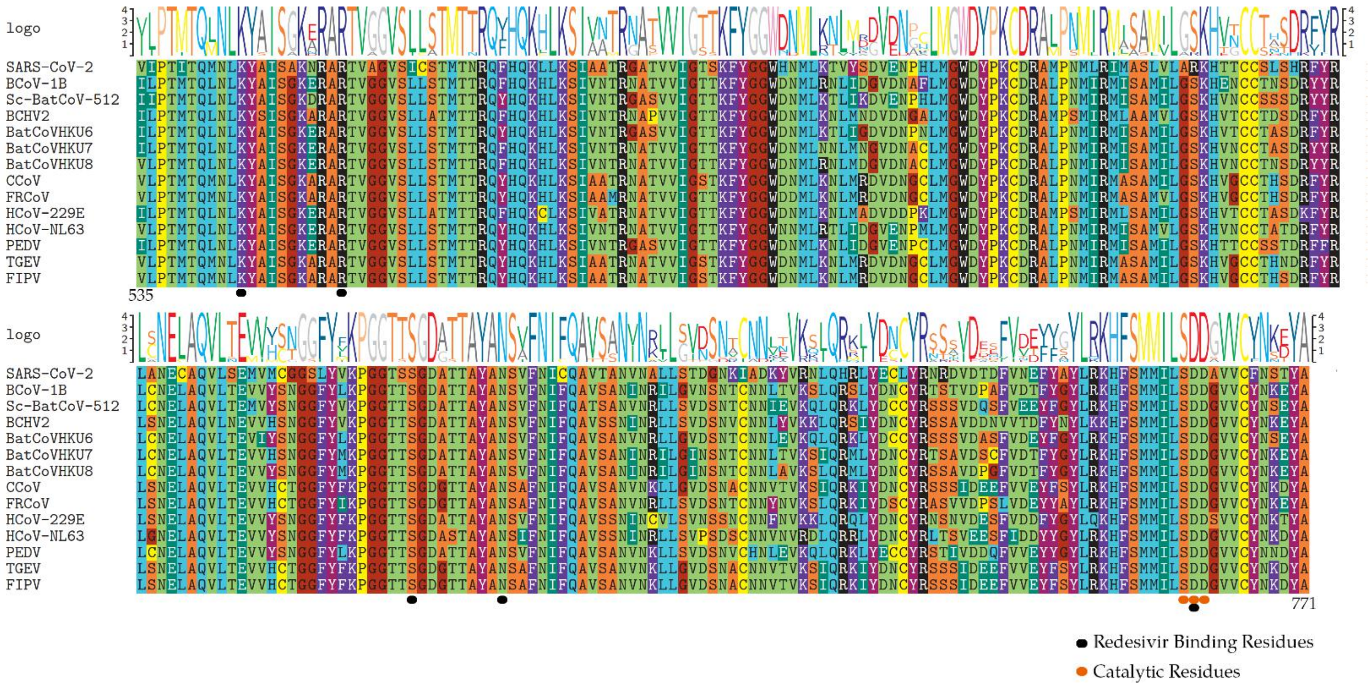


B

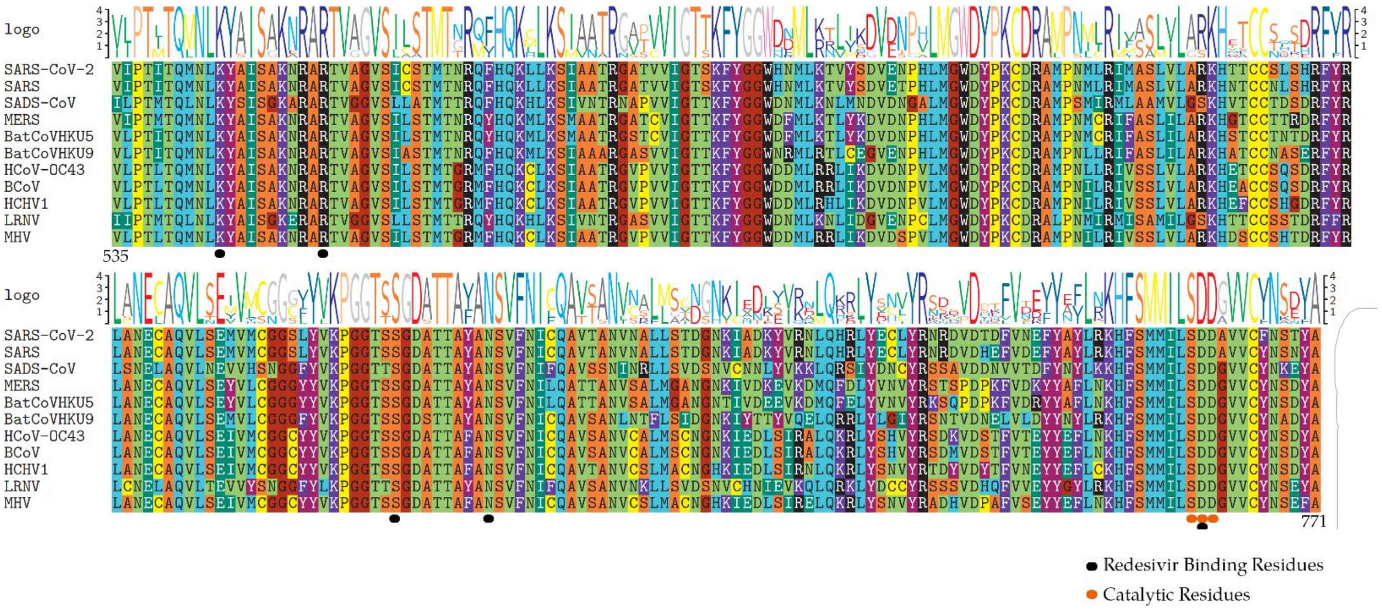
