## Supplemental for "The adenosine analogue prodrug ATV006 is orally bioavailable and has potent preclinical efficacy against SARS-CoV-2 and its variants"

#### Scheme 1. Synthesis of 69-0 prodrugs [a]

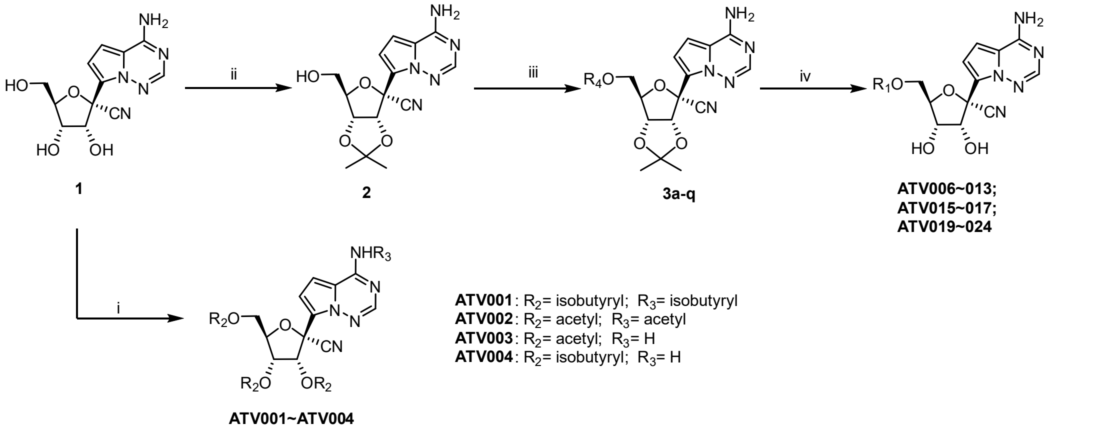

**3a, ATV006**: $R_1$=$R_4$= isobutyryl
**3b, ATV007**: $R_1$=$R_4$= acetyl
**3c, ATV008**: $R_1$=$R_4$= propionyl
**3d, ATV009**: $R_1$=$R_4$= *n*-butyryl
**3e, ATV010**: $R_1$=$R_4$= *n*-nonanoyl
**3f, ATV011**: $R_1$=$R_4$= 2-ethylbutanoyl
**3g, ATV012**: $R_1$=$R_4$= cyclopropanecarbonyl
**3h, ATV013**: $R_1$=$R_4$= benzoyl
**3i, ATV015**: $R_1$=$R_4$= cyclopentanecarbonyl
**3j, ATV016**: $R_1$=$R_4$= 3,3,3-trifluoropropanoyl
**3k, ATV017**: $R_1$=$R_4$= 3-methylbutanoyl
**3l, ATV019**: $R_1$= *D*-Valyl; $R_4$= *N*-Boc-*D*-Valyl
**3m, ATV020**: $R_1$= *L*-Valyl; $R_4$= *N*-Boc-*L*-Valyl
**3n, ATV021**: $R_1$= *L*-phenylalanyl; $R_4$= *N*-Boc-*L*-phenylalanyl
**3o, ATV022**: $R_1$= *D*-phenylalanyl; $R_4$= *N*-Boc-*D*-phenylalanyl
**3p, ATV023**: $R_1$= (2*S*)-2-amino-3-methylpentanoyl; $R_4$= *N*-Boc-(2*S*)-2-amino-3-methylpentanoyl
**3q, ATV024**: $R_1$= (2*R*)-2-amino-3-methylpentanoyl; $R_4$= *N*-Boc-(2*R*)-2-amino-3-methylpentanoyl

[a] Reaction conditions: i) $R_2OR_2$, DMAP, EDMA, ACN, 40 °C, 0.5 h; ii) 2,2-Dimethoxypropane, Conc. $H_2SO_4$, Acetone, rt~45 °C, 4 h; iii) ROH, DCC, DMAP, DCM, rt, 12 h; iv) 6 N HCl, THF, 0 °C, 7 h.

#### General procedure:

All reagents used were commercially available. Reactions were monitored by thin-layer chromatography (TLC) on glass plates coated with silica gel with a fluorescent indicator (GF254). Flash silica gel column chromatography was performed using Tsingdao silica gel (60, particle size 300-400 mesh). All the $^1$H NMR and $^{13}$C NMR spectra were recorded on a Bruker 400 MHz or 600 MHz spectrometer. Chemical shifts (d) were expressed in parts per million using tetramethylsilane as an internal reference. High-resolution mass spectra (HRMS) were measured with an Agilent Accurate-Mass Q-TOF 6530 in ESI mode (Agilent, Santa Clara, CA, USA). HPLC analyses were performed using a Hewlett Packard Model HP 1100 Series instruments, the compounds are at least ≥95% pure.

**(2*R*,3*R*,4*R*,5*R*)-2-cyano-2-(4-isobutyramidopyrrolo[2,1-*f*][1,2,4]triazin-7-yl)-5-((isobutyryloxy)methyl)tetrahydrofuran-3,4-diyl bis(2-methylpropanoate) (ATV001)**

To a solution of **1** (594 mg, 2 mmol), 4-dimethylaminopyridine (50 mg, 0.4 mmol), EDMA (1.2 mL, 11 mmol) in acetonitrile (10 mL) was added isobutyric anhydride (1.66 mL, 10 mmol). The mixture was stirred at 40 °C for 1 hour. The mixture was concentrated under vacuum to give crude product which was purified by silica gel column chromatography using MeOH/DCM (V/V=5/95) as eluent. **ATV001** was isolated as colorless liquid (624 mg, 61%). HPLC purity: 95.83% (eluent, water/ACN = 10/90; flow rate 1 mL/min; wavelength 254 nm). $^1$H NMR (400 MHz, $CDCl_3$) δ 9.33 (s, 1H), 8.21 (s, 1H), 7.34 (d, *J* = 4.9 Hz, 1H), 7.06 (d, *J* = 4.9 Hz, 1H), 6.23 (d, *J* = 5.8 Hz, 1H), 5.51 (dd, *J* = 5.8, 4.3 Hz, 1H), 4.67 (q, *J* = 4.0 Hz, 1H), 4.41 (qd, *J* = 12.3, 3.9 Hz, 2H), 3.19 (dt, *J* = 13.4, 6.7 Hz, 1H), 2.74-2.62 (m, 2H), 2.56 (dq, *J* = 14.0, 7.0 Hz, 1H), 1.35-1.10 (m, 24H). $^{13}$C NMR (101 MHz, $CDCl_3$) δ 177.46, 176.45, 175.76, 174.98, 151.38, 145.87, 123.21, 118.26, 114.91, 113.27, 106.29, 81.60, 76.86, 71.97, 70.54, 62.56, 36.01, 33.85, 33.84, 33.74, 19.13, 19.11, 18.91, 18.85, 18.81, 18.70, 18.67, 18.54. HRMS (m/z): calculated for $C_{28}H_{38}N_5O_8^+$ [M+H]$^+$ 572.2715; found, 572.2705.

**(2*R*,3*R*,4*R*,5*R*)-2-(4-acetamidopyrrolo[2,1-*f*][1,2,4]triazin-7-yl)-5-(acetoxymethyl)-2-cyanotetrahydrofuran-3,4-diyl diacetate (ATV002)**

**ATV002** was prepared using a procedure analogous to the synthesis of compound **ATV001** but acetic anhydride in place of isobutyric anhydride. **ATV002** was isolated as a white solid (518 mg, 56% yield). HPLC purity: 100% (eluent, water/ACN = 10/90; flow rate 1 mL/min; wavelength 254 nm). $^1$H NMR (400 MHz, $CDCl_3$) δ 9.16 (s, 1H), 8.23 (s, 1H), 7.21 (d, *J* = 4.8 Hz, 1H), 7.11 (d, *J* = 4.8 Hz, 1H), 6.25 (d, *J* = 5.9 Hz, 1H), 5.56-5.41 (m, 1H), 4.65 (dd, *J* = 8.5, 4.7 Hz, 1H), 4.47 (dd, *J* = 12.3, 3.6 Hz, 1H), 4.34 (dd, *J* = 12.3, 4.9 Hz, 1H), 2.63 (s, 3H), 2.19 (s, 3H), 2.17 (s, 3H), 2.09 (s, 3H). $^{13}$C NMR (101 MHz, $CDCl_3$) δ 172.03, 170.43, 169.84, 169.03, 151.01, 146.16, 122.96, 117.82, 114.85, 114.01, 103.74, 81.00, 77.21, 71.79, 70.60, 62.58, 26.12, 20.76, 20.53, 20.51. HRMS (m/z): calculated for $C_{20}H_{22}N_5O_8^+$ [M+H]$^+$ 460.1463; found, 460.1454.

**(2*R*,3*R*,4*R*,5*R*)-5-(acetoxymethyl)-2-(4-aminopyrrolo[2,1-*f*][1,2,4]triazin-7-yl)-2-cyanotetrahydrofuran-3,4-diyl diacetate (ATV003)**

**ATV003** was prepared using a procedure analogous to the synthesis of compound **ATV002**. **ATV003** was isolated as a white solid (384 mg, 46%). HPLC purity: 96.10% (eluent, water/ACN = 10/90; flow rate 1 mL/min; wavelength 254 nm). $^1$H NMR (400 MHz, $CDCl_3$) δ 7.94 (s, 1H), 6.92 (d, *J* = 4.6 Hz, 1H), 6.61 (d, *J* = 4.7 Hz, 1H), 6.30 (d, *J* = 5.9 Hz, 3H), 5.61-5.43 (m, 1H), 4.63 (dd, *J* = 8.7, 4.9 Hz, 1H), 4.49 (dd, *J* = 12.2, 3.7 Hz, 1H), 4.34 (dd, *J* = 12.2, 5.1 Hz, 1H), 2.18 (s, 3H), 2.16 (s, 3H), 2.08 (s, 3H). $^{13}$C NMR (101 MHz, $CDCl_3$) δ 170.55, 169.91, 169.16, 155.54, 147.39, 121.63, 117.23, 115.28, 112.61, 100.23, 80.85, 77.48, 71.90, 70.67, 62.67, 20.77,

20.55. HRMS (m/z): calculated for $C_{18}H_{20}N_5O_7^+$ $[M+H]^+$ 418.1357; found, 418.1348.

**(2*R*,3*R*,4*R*,5*R*)-2-(4-aminopyrrolo[2,1-*f*][1,2,4]triazin-7-yl)-2-cyano-5-((isobutyryl oxy)methyl)tetrahydrofuran-3,4-diyl bis(2-methylpropanoate) (ATV004)**

**ATV004** was prepared using a procedure analogous to the synthesis of compound **ATV001**. **ATV004** was isolated as colorless liquid (410 mg, 35%). HPLC purity: 97.49% (eluent, water/ACN = 10/90; flow rate 1 mL/min; wavelength 254 nm). $^1H$ NMR (400 MHz, $CDCl_3$) δ 7.89 (s, 1H), 6.86 (d, *J* = 4.7 Hz, 1H), 6.70 (d, *J* = 4.7 Hz, 1H), 6.28 (d, *J* = 5.9 Hz, 1H), 5.53 (dd, *J* = 5.7, 4.4 Hz, 1H), 4.65 (q, *J* = 4.1 Hz, 1H), 4.42 (qd, *J* = 12.3, 4.1 Hz, 2H), 2.75-2.51 (m, 3H), 1.32-1.10 (m, 18H). $^{13}C$ NMR (101 MHz, $CDCl_3$) δ 176.58, 175.85, 175.11, 155.65, 146.56, 122.08, 117.09, 115.34, 112.03, 101.09, 81.50, 77.04, 71.99, 70.63, 62.66, 33.85, 33.82, 33.74, 18.96, 18.82, 18.78, 18.69, 18.67, 18.54. HRMS (m/z): calculated for $C_{24}H_{32}N_5O_7^+$ $[M+H]^+$ 502.2296; found, 502.2287.

**(3a*R*,4*R*,6*R*,6a*R*)-4-(4-aminopyrrolo[2,1-*f*][1,2,4]triazin-7-yl)-6-(hydroxymethyl)-2,2-dimethyltetrahydrofuro[3,4-*d*][1,3]dioxole-4-carbonitrile (2)**

To a solution of **1** (5.62 g, 19.3 mmol) in acetone (30 mL) was added 2,2-dimethoxypropane (11.5 mL, 92.6 mmol). Then Conc. $H_2SO_4$ (1.34 mL, 25.1 mmol) was added dropwise at room temperature. The mixture was stirred at 45°C. After 0.5 h, the reaction was completed as monitored by TLC. The mixture was neutralized with saturated $NaHCO_3$ and then concentrated. The residue was extracted into ethyl acetate (100 mL×3). The combined organic extracts were washed with water and brine, dried with anhydrous $Na_2SO_4$, filtered and concentrated in *vacuo* to give crude product which was purified by silica gel column chromatography using PE/EA (V/V) = 1/2) as eluent. **5** was obtained as a white solid (6.20 g, 97%). $^1H$ NMR (400 MHz, Chloroform-*d*) δ 7.95 (s, 1H), 7.11 (d, *J* = 4.7 Hz, 1H), 6.69 (dd, *J* = 4.8, 2.4 Hz, 1H), 5.77 (s, 2H), 5.42 (d, *J* = 6.6 Hz, 1H), 5.24 (dd, *J* = 6.6, 2.4 Hz, 1H), 4.67 (q, *J* = 1.9 Hz, 1H), 3.99 (dd, *J* = 12.5, 1.9 Hz, 1H), 3.84 (dd, *J* = 12.5, 1.7 Hz, 1H), 1.81 (s, 3H), 1.40 (s, 3H).

**((3a*R*,4*R*,6*R*,6a*R*)-6-(4-aminopyrrolo[2,1-*f*][1,2,4]triazin-7-yl)-6-cyano-2,2-dimeth yltetrahydrofuro[3,4-*d*][1,3]dioxol-4-yl)methyl isobutyrate (3a)**

To a solution of compound **2** (1.50 g, 4.5 mmol), isobutyric acid (0.42 mL, 4.5 mmol), 4-dimethylaminopyridine (55.40 mg, 0.45 mmol) in DCM (15 mL) was added dicyclohexylcarbodiimide (1.02 g, 4.9 mmol). The mixture was stirred at room temperature for 24

h. The suspension was filtered and the solution was washed with 30 mL of saturated solution of $Na_2CO_3$ and then with 30 mL of an aqueous solution of citric acid (20 % w/v). The organic layer was dried with $Na_2SO_4$ and the solvent was removed under reduced pressure. The products were purified by column chromatography (PE/EA = 1:1). Compound **3a** was isolated as a white solid (1.71 g, 94% yield). $^1H$ NMR (400 MHz, $CDCl_3$) δ (ppm): 7.99 (s, 1H), 6.99 (d, $J$ = 4.6 Hz, 1H), 6.62 (d, $J$ = 4.6 Hz, 1H), 5.72 (br, 2H), 5.49 (d, $J$ = 6.8 Hz, 1H), 4.93-4.90 (dd, $J$ = 6.8 Hz, 4.3 Hz, 1H), 4.61-4.58 (q, $J$ = 4.4 Hz, 1H), 4.44-4.26 (m, 2H), 2.61-2.50 (m, 1H), 1.77 (s, 3H), 1.42 (s, 3H), 1.17-1.14 (q, $J$ = 3.8 Hz, 6H). $^{13}C$ NMR (100 MHz, $CDCl_3$) δ (ppm): 176.7, 155.2, 147.3, 123.5, 117.2, 116.7, 115.6, 112.6, 100.0, 83.8, 83.0, 82.0, 81.4, 63.1, 33.8, 26.4, 25.6, 18.9.

**((2*R*,3*S*,4*R*,5*R*)-5-(4-aminopyrrolo[2,1-*f*][1,2,4]triazin-7-yl)-5-cyano-3,4-dihydroxytetrahydrofuran-2-yl)methyl isobutyrate (ATV006)**

Compound **3a** (1.50 g, 3.7 mmol) was dissolved in 37% hydrochloric acid aqueous solution (3 mL) and tetrahydrofuran (15 mL). After stirring for 6 hours, sodium carbonate was added to adjust pH to 8, the solvent was removed *in vacuo*, and the residue was purified by silica gel column chromatography using PE/EA (V/V=1/3) as eluent. **ATV006** was isolated as a white solid (0.66 g, 49% yield). HPLC purity: 100% (OD-3; eluent, n-hexane/isopropanol = 80/20; flow rate 0.8 mL/min; temperature 30 °C; wavelength 254 nm; HPLC analysis data are reported in relative area % and were not adjusted to weight %). $^1H$ NMR (400 MHz, CD3OD) δ 7.76 (s, 1H), 6.78 (s, 2H), 4.78 (d, $J$ = 5.3 Hz, 1H), 4.40-4.24 (m, 2H), 4.24-4.11 (m, 1H), 4.10-4.01 (m, 1H), 2.42 (p, $J$ = 7.0 Hz, 1H), 0.99 (dd, $J$ = 7.0, 4.1 Hz, 6H). $^{13}C$ NMR (101 MHz, CD3OD) δ 176.96, 155.82, 146.92, 124.25, 116.54, 116.29, 110.75, 101.20, 82.04, 80.00, 74.27, 70.68, 62.93, 33.58, 25.00, 17.95, 17.87. HRMS (m/z): calculated for $C_{16}H_{20}N_5O_5^+$ $[M+H]^+$ 362.1459; found, 362.1451.

**((2*R*,3*S*,4*R*,5*R*)-5-(4-aminopyrrolo[2,1-*f*][1,2,4]triazin-7-yl)-5-cyano-3,4-dihydroxytetrahydrofuran-2-yl)methyl acetate (ATV007)**

Compound **3b** was prepared using a procedure analogous to the synthesis of compound **3a** but acetic acid in place of isobutyric acid. **3b** was obtained as a white solid (1.78 g, 98%).

**ATV007** was prepared using a procedure analogous to the synthesis of compound **ATV006** but compound **3b** in place of compound **3a**. **ATV007** was obtained as a white solid (0.68 g, 51% yield). HPLC purity: 96.46% (OD-3; eluent, *n*-hexane/isopropanol = 80/20; flow rate 0.8 mL/min;

temperature 30 °C; wavelength 254 nm; HPLC analysis data are reported in relative area % and were not adjusted to weight %). $^1$H NMR (600 MHz, $CD_3OD$) δ (ppm): 7.86 (s, 1H), 6.89 (t, *J* = 5.0 Hz, 2H), 4.87 (s, 1H), 4.43-4.41 (dd, *J* = 12 Hz, 2.8 Hz, 1H), 4.37-4.34 (m, 1H), 4.30-4.27 (m, 1H), 4.13 (t, *J* = 5.7 Hz, 1H), 2.03 (s, 3H). $^{13}$C NMR (150 MHz, $CD_3OD$) δ (ppm): 171.0, 155.8, 146.9, 124.2, 116.6, 116.2, 110.7, 101.1, 81.9, 80.2, 74.1, 70.7, 63.1, 19.3. HRMS (m/z): calculated for $C_{14}H_{16}N_5O_5^+$ $[M+H]^+$ 334.1146; found, 334.1140.

**((2*R*,3*S*,4*R*,5*R*)-5-(4-aminopyrrolo[2,1-*f*][1,2,4]triazin-7-yl)-5-cyano-3,4-dihydroxytetrahydrofuran-2-yl)methyl propionate (ATV008)**

Compound **3c** was prepared using a procedure analogous to the synthesis of compound **3a** but propionic acid in place of isobutyric acid. **3c** was obtained as a white solid (1.74 g, 99% yield).

**ATV008** was prepared using a procedure analogous to the synthesis of compound **ATV006** but **3c** in place of compound **3a**. **ATV008** was obtained as a white solid (0.68 g, 48% yield). HPLC purity 99.56% (OD-3; eluent, *n*-hexane/isopropanol = 80/20; flow rate 0.8 mL/min; temperature 30 °C; wavelength 254 nm; HPLC analysis data are reported in relative area % and were not adjusted to weight %). $^1$H NMR (600 MHz, $CD_3OD$) δ (ppm): 7.86 (s, 1H), 6.90-6.88 (q, *J* = 4.5 Hz, 2H), 4.87-4.86 (m, 1H), 4.46-4.43 (dd, *J* = 12, 2.8 Hz, 1H), 4.37-4.36 (m, 1H), 4.31-4.28 (m, 1H), 4.15 (t, *J* = 5.8 Hz, 1H), 2.38-2.28 (m, 2H), 1.08 (t, *J* = 7.5 Hz, 3H). $^{13}$C NMR (150 MHz, $CD_3OD$) δ (ppm): 174.3, 155.8, 146.9, 124.2, 116.5, 116.2, 110.7, 101.1, 82.0, 80.1, 74.2, 70.7, 62.9, 26.7, 7.9. HRMS (m/z): calculated for $C_{15}H_{18}N_5O_5^+$ $[M+H]^+$ 348.1302; found, 348.1296.

**((2*R*,3*S*,4*R*,5*R*)-5-(4-aminopyrrolo[2,1-*f*][1,2,4]triazin-7-yl)-5-cyano-3,4-dihydroxytetrahydrofuran-2-yl)methyl butyrate (ATV009)**

Compound **3d** was prepared using a procedure analogous to the synthesis of compound **3a** but *n*-butyric acid in place of isobutyric acid. **3d** was obtained as a white solid (1.78 g, 98%).

**ATV009** was prepared using a procedure analogous to the synthesis of compound **ATV006** but compound **3d** in place of compound **3a**. **ATV009** was obtained as a white solid (0.76 g, 56%). HPLC purity 99.52% (OD-3; eluent, *n*-hexane/isopropanol = 80/20; flow rate 0.8 mL/min; temperature 30 °C; wavelength 254 nm; HPLC analysis data are reported in relative area % and were not adjusted to weight %). $^1$H NMR (600 MHz, $CD_3OD$) δ (ppm): 7.86 (s, 1H), 6.90-6.88 (q,

*J* = 4.5 Hz, 2H), 4.87-4.86 (m, 1H), 4.44-4.42 (dd, *J* = 12 Hz, 2.8 Hz, 1H), 4.37-4.35 (m, 1H), 4.31-4.28 (m, 1H), 4.14 (t, *J* = 5.8 Hz, 1H), 2.32-2.23 (m, 2H), 1.62-1.56 (m,2H), 0.91 (t, *J* = 7.4 Hz, 3H). $^{13}$C NMR (150 MHz, $CD_3OD$) δ (ppm): 174.3, 155.9, 146.9, 124.3, 116.5, 116.2, 110.7, 101.1, 82.0, 80.1, 74.2, 70.7, 62.8, 35.4, 17.9, 12.5. HRMS (m/z): calculated for $C_{16}H_{20}N_5O_5^+$ $[M+H]^+$ 362.1459; found, 362.1452.

#### ((2*R*,3*S*,4*R*,5*R*)-5-(4-aminopyrrolo[2,1-*f*][1,2,4]triazin-7-yl)-5-cyano-3,4-dihydroxytetrahydrofuran-2-yl)methyl nonanoate (ATV010)

Compound **3e** was prepared using a procedure analogous to the synthesis of compound **3a** but pelargonic acid in place of isobutyric acid. **3e** was obtained as a white solid (2.07 g, 97%).

**ATV010** was prepared using a procedure analogous to the synthesis of compound **ATV006** but compound **3e** in place of compound **3a**. **ATV010** was obtained as a white solid (0.55 g, 40.3%). HPLC purity was 97.84% (OD-3; eluent, *n*-hexane/isopropanol = 80/20; flow rate 0.8 mL/min; temperature 30 °C; wavelength 254 nm; HPLC analysis data are reported in relative area % and were not adjusted to weight %). $^1$H NMR (600 MHz, $CD_3OD$) δ (ppm): 7.86 (s, 1H), 6.90-6.88 (q, *J* = 4.5 Hz, 2H), 4.87-4.86 (m, 1H), 4.43-4.41 (dd, *J* = 12 Hz, 2.8 Hz, 1H), 4.37-4.35 (m, 1H), 4.32-4.29 (m, 1H), 4.14 (t, *J* = 5.8 Hz, 1H), 2.38-2.23 (m, 2H), 1.56-1.53 (m, 2H), 1.29-1.27 (m, 10H), 0.87 (t, *J* = 7.0 Hz, 3H). $^{13}$C NMR (150 MHz, $CD_3OD$) δ (ppm): 173.7, 155.9, 146.9, 124.3, 116.5, 116.2, 110.7, 101.1, 82.0, 74.2, 70.7, 62.8, 33.5, 31.5, 28.8, 28.7, 24.6, 22.3. HRMS (m/z): calculated for $C_{21}H_{30}N_5O_5^+$ $[M+H]^+$ 432.2241; found, 432.2234.

#### ((2*R*,3*S*,4*R*,5*R*)-5-(4-aminopyrrolo[2,1-*f*][1,2,4]triazin-7-yl)-5-cyano-3,4-dihydroxytetrahydrofuran-2-yl)methyl 2-ethylbutanoate (ATV011)

Compound **3f** was prepared using a procedure analogous to the synthesis of compound **3a** but 2-ethyl butyric acid in place of isobutyric acid. **3f** was obtained as a white solid (1.94 g, 99%).

**ATV011** was prepared using a procedure analogous to the synthesis of compound **ATV006** but compound **3f** in place of compound **3a**. **ATV011** was obtained as a white solid (0.70 g, 51.3%). HPLC purity 99.61% (OD-3; eluent, *n*-hexane/isopropanol = 80/20; flow rate 0.8 mL/min; temperature 30 °C; wavelength 254 nm; HPLC analysis data are reported in relative area % and were not adjusted to weight %). $^1$H NMR (600 MHz, $CD_3OD$) δ (ppm): 7.86 (s, 1H), 6.89 (s,2H),

4.87-4.86 (m, 1H), 4.39-4.43 (dd, *J* = 12 Hz, 2.8 Hz, 1H), 4.37-4.35 (m, 1H), 4.14 (t, *J* = 5.8 Hz, 1H), 2.38-2.22 (m, 1H), 1.60-1.45 (m, 4H), 0.86-0.82 (m, 6H). $^{13}$C NMR (150 MHz, $CD_3OD$) δ (ppm): 176.1, 155.9, 146.9, 124.3, 116.6, 116.2, 110.7, 101.1, 81.9, 79.9, 74.2, 70.7, 62.8, 48.9, 24.7, 24.6. 10.7, 10.6. HRMS (m/z): calculated for $C_{18}H_{24}N_5O_5^+$ $[M+H]^+$ 390.1772; found, 390.1764.

#### ((2*R*,3*S*,4*R*,5*R*)-5-(4-aminopyrrolo[2,1-*f*][1,2,4]triazin-7-yl)-5-cyano-3,4-dihydroxytetrahydrofuran-2-yl)methyl cyclopropanecarboxylate (ATV012)

Compound **3g** was prepared using a procedure analogous to the synthesis of compound **3a** but cyclopropanoic acid in place of isobutyric acid. **3g** was obtained as a white solid (1.52 g, 99%). **ATV012** was prepared using a procedure analogous to the synthesis of compound **ATV006** but compound **3g** in place of compound **3a**. **ATV012** was obtained as a white solid (0.98 g, 62%). HPLC purity 98.56% (OD-3; eluent, *n*-hexane/isopropanol = 80/20; flow rate 0.8 mL/min; temperature 30 °C; wavelength 254 nm; HPLC analysis data are reported in relative area % and were not adjusted to weight %). $^1$H NMR (600 MHz, $CD_3OD$) δ (ppm): 7.86 (s, 1H), 6.89 (t, *J* = 4.5Hz, 2H), 4.87-4.86 (m, 1H), 4.46-4.44 (dd, *J* = 12 Hz, 2.8 Hz, 1H), 4.36-4.34 (m, 1H), 4.29-4.26 (m, 1H), 4.15 (t, *J* = 5.8 Hz, 1H), 1.64-1.60 (m, 1H), 0.92-0.87 (m, 4H). $^{13}$C NMR (150 MHz, $CD_3OD$) δ (ppm): 174.9, 155.9, 146.9, 124.2, 116.6, 116.2, 110.7, 101.1, 80.2, 80.1, 74.2, 70.6, 63.0, 12.1, 7.5, 7.4. HRMS (m/z): calculated for $C_{16}H_{18}N_5O_5^+$ $[M+H]^+$ 360.1302; found, 360.1295.

#### ((2*R*,3*S*,4*R*,5*R*)-5-(4-aminopyrrolo[2,1-*f*][1,2,4]triazin-7-yl)-5-cyano-3,4-dihydroxytetrahydrofuran-2-yl)methyl benzoate (ATV013)

**ATV013** was prepared using a procedure analogous to the synthesis of compound **ATV006**. **ATV013** was obtained as a white solid (0.21 g, 34.9% total yield). HPLC purity 99.29% (OD-3; eluent, *n*-hexane/isopropanol = 80/20; flow rate 0.8 mL/min; temperature 30 °C; wavelength 254 nm; HPLC analysis data are reported in relative area % and were not adjusted to weight %). $^1$H NMR (600 MHz, DMSO-$d_6$) δ (ppm): 7.92 (br, 2H), 7.90 (d, *J* = 7.4 Hz, 2H), 7.86 (s, 1H), 7.68 (t, *J* = 7.4 Hz, 1H), 7.52 (t, *J* = 7.7 Hz, 2H), 6.87 (d, *J* = 4.5 Hz, 1H), 6.81 (d, *J* = 4.5 Hz, 1H), 6.36 (d, *J* = 5.9 Hz, 1H), 5.46 (d, *J* = 5.9 Hz, 1H), 4.79 (t, *J* = 5.3 Hz, 1H), 4.61-4.58 (dd, *J* = 12.2 Hz, 2.6

Hz, 1H), 4.45-4.42 (dd, $J$ = 12.3 Hz, 4.8 Hz, 1H), 4.39-4.37 (m, 1H), 4.14-4.10 (m, 1H). $^{13}C$ NMR (150 MHz, DMSO-$d_6$) δ (ppm): 166.0, 156.1,148.4, 134.0, 129.8, 129.7, 129.2, 123.9, 117.4, 117.1, 110.8, 101.3, 81.7, 79.7, 74.5, 70.6, 63.9. HRMS (m/z): calculated for $C_{19}H_{18}N_5O_5^+$ $[M+H]^+$ 396.1302; found, 396.1296.

#### ((2*R*,3*S*,4*R*,5*R*)-5-(4-aminopyrrolo[2,1-*f*][1,2,4]triazin-7-yl)-5-cyano-3,4-dihydroxytetrahydrofuran-2-yl)methyl cyclopentanecarboxylate (ATV015)

**ATV015** was prepared using a procedure analogous to the synthesis of compound **ATV006**. **ATV015** was obtained as a white solid (0.33 g, 56.1% total yield). HPLC purity 99.22% (OD-3; eluent, *n*-hexane/isopropanol = 80/20; flow rate 0.8 mL/min; temperature 30 °C; wavelength 254 nm; HPLC analysis data are reported in relative area % and were not adjusted to weight %). $^1H$ NMR (600 MHz, $CD_3OD$) δ (ppm): 7.86 (s, 1H), 6.90-6.87 (q, $J$ = 4.6 Hz, 2H), 4.85-4.83 (m, 1H), 4.39-4.43 (dd, $J$ = 12.1 Hz, 3.1 Hz, 1H), 4.37-4.35 (m, 1H), 4.14 (t, $J$ = 5.7 Hz, 1H), 2.75-2.70 (m, 1H), 1.87-1.80 (m, 2H), 1.75-1.53 (m, 6H). $^{13}C$ NMR (150 MHz, $CD_3OD$) δ (ppm): 176.5, 155.9, 146.9, 124.3, 116.5, 116.2, 110.7, 101.1, 82.0, 80.0, 74.3, 70.7, 62.8, 43.5, 29.5, 29.4, 25.3. HRMS (m/z): calculated for $C_{18}H_{22}N_5O_5^+$ $[M+H]^+$ 388.1615; found, 388.1607.

#### ((2*R*,3*S*,4*R*,5*R*)-5-(4-aminopyrrolo[2,1-*f*][1,2,4]triazin-7-yl)-5-cyano-3,4-dihydroxytetrahydrofuran-2-yl)methyl 3,3,3-trifluoropropanoate (ATV016)

**ATV016** was prepared using a procedure analogous to the synthesis of compound **ATV006**. **ATV016** was obtained as a white solid (0.31 g, 50.8% total yield). HPLC purity 98.84% (OD-3; eluent, *n*-hexane/isopropanol = 80/20; flow rate 0.8 mL/min; temperature 30 °C; wavelength 254 nm; HPLC analysis data are reported in relative area % and were not adjusted to weight %). $^1H$ NMR (600 MHz, $CD_3OD$) δ (ppm): 7.86 (s, 1H), 6.90-6.88 (q, $J$ = 4.6 Hz, 2H), 4.89 (d, $J$ = 5.3 Hz, 1H), 4.54-4.50 (m, 1H), 4.42-4.38 (m, 2H), 4.15 (t, $J$ = 5.7 Hz, 1H), 3.45-3.35 (m, 2H). $^{13}C$ NMR (150 MHz, $CD_3OD$) δ (ppm): 164.3 ($J$ = 4.0 Hz), 155.5, 146.9, 123.8 (q, $J$ = 273.6 Hz), 124.1, 116.6, 116.2, 110.8, 101.2, 81.7, 80.2, 74.0, 70.6, 64.1. HRMS (m/z): calculated for $C_{15}H_{15}F_3N_5O_5^+$ $[M+H]^+$ 402.1019; found, 402.1013.

**((2*R*,3*S*,4*R*,5*R*)-5-(4-aminopyrrolo[2,1-*f*][1,2,4]triazin-7-yl)-5-cyano-3,4-dihydroxytetrahydrofuran-2-yl)methyl 3-methylbutanoate (ATV017)**

**ATV017** was prepared using a procedure analogous to the synthesis of compound **ATV006**. **ATV017** was obtained as a white solid (0.27 g, 47.2% total yield). HPLC purity 97.35% (OD-3; eluent, n-hexane/isopropanol = 80/20; flow rate 0.8 mL/min; temperature 30 °C; wavelength 254 nm; HPLC analysis data are reported in relative area % and were not adjusted to weight %). $^1$H NMR (600 MHz, $CD_3OD$) δ (ppm): 7.86 (s, 1H), 6.90-6.88 (q, *J* = 4.6 Hz, 2H), 4.87 (d, *J* = 5.3 Hz, 1H), 4.43-4.40 (m, 1H), 4.39-4.35 (m, 2H), 4.31-4.29 (m,1 H), 4.14 (t, *J* = 5.7 Hz, 1H), 2.18-2.16 (m, 2H), 2.04-1.97 (m, 1H), 0.91-0.90 (q, *J* = 3.2 Hz, 6H). $^{13}$C NMR (150 MHz, $CD_3OD$) δ (ppm): 155.9, 146.9, 124.3, 116.5, 116.2, 110.7, 101.1, 82.0, 80.0, 74.2, 70.7, 70.6, 62.8, 62.7, 42.6, 25.4, 21.3, 21.2. HRMS (m/z): calculated for $C_{17}H_{22}N_5O_5^+$ $[M+H]^+$ 376.1615; found, 376.1608.

**((3a*R*,4*R*,6*R*,6a*R*)-6-(4-aminopyrrolo[2,1-*f*][1,2,4]triazin-7-yl)-6-cyano-2,2-dimethyltetrahydrofuro[3,4-*d*][1,3]dioxol-4-yl)methyl (tert-butoxycarbonyl)-*D*-valinate (3l)**

To a solution of compound **2** (1.80 g, 5.4 mmol), (*tert*-butoxycarbonyl)-*D*-valine (1.18 g, 5.4 mmol), 4-dimethylaminopyridine (66.48 mg, 0.54 mmol) in DCM (15 mL) was dropwise added a solution of dicyclohexylcarbodiimide (1.22 g, 6 mmol) in DCM (5 mL). The mixture was stirred at room temperature for 24 h. The suspension was filtered and the solution was washed with 30 mL of saturated solution of $Na_2CO_3$ and then with 30 mL of an aqueous solution of citric acid (20 % w/v). The organic layer was dried with $Na_2SO_4$ and the solvent was removed under reduced pressure. The products were purified by column chromatography (PE/EA = 1:1). Compound **3l** was isolated as a white solid (2.81 g, 97%). $^1$H NMR (400 MHz, CD3OD). $^1$H NMR (600 MHz, CD3OD) δ 7.79 (s, 1H), 6.79 (s, 2H), 5.39 (s, 1H), 4.90 (dd, *J* = 6.5, 3.4 Hz, 1H), 4.51 (q, *J* = 4.1 Hz, 1H), 4.29 (dd, *J* = 12.0, 3.8 Hz, 1H), 4.24 (dd, *J* = 12.1, 5.2 Hz, 1H), 3.77 (d, *J* = 6.0 Hz, 1H), 3.27-3.11 (m, 1H), 1.61 (s, 4H), 1.32 (d, *J* = 2.5 Hz, 9H), 1.24 (s, 3H), 0.73 (dd, *J* = 19.0, 6.8 Hz, 6H). $^{13}$C NMR (151 MHz, MeOD) δ 172.00, 156.84, 155.83, 147.06, 123.47, 116.84, 116.25, 115.65, 110.76, 101.11, 84.49, 82.89, 82.02, 81.17, 79.18, 63.54, 59.24, 53.42, 48.04, 47.91, 47.90, 47.84, 47.76, 47.62, 47.56, 47.48, 47.33, 47.19, 33.37, 30.06, 27.32, 25.35, 25.14, 24.66, 24.14, 18.14, 16.90.

**((2*R*,3*S*,4*R*,5*R*)-5-(4-aminopyrrolo[2,1-*f*][1,2,4]triazin-7-yl)-5-cyano-3,4-dihydroxytetrahydrofuran-2-yl)methyl *D*-valinate (ATV019)**

Compound **3l** (2.50 g, 4.7 mmol) was dissolved in 37% hydrochloric acid aqueous solution (3 mL) and tetrahydrofuran (15 mL). After stirring for 6 hours, sodium carbonate was added to adjust pH to 8, the solvent was removed *in vacuo*, and the residue was purified by silica gel column chromatography using MeOH/EA (V/V=1/20) as eluent. **ATV019** was isolated as a white solid (0.99 g, 54%). HPLC purity 100% (OD-3; eluent, *n*-hexane/isopropanol = 80/20; flow rate 0.8 mL/min; temperature 30 °C; wavelength 254 nm; HPLC analysis data are reported in relative area % and were not adjusted to weight %). $^{1}$H NMR (400 MHz, CD3OD) δ 7.76 (s, 1H), 6.80 (s, 2H), 4.79 (s, 1H), 4.42-4.24 (m, 3H), 4.08 (d, *J* = 5.5 Hz, 1H), 3.23 (d, *J* = 11.1 Hz, 1H), 1.90-1.76 (m, 1H), 0.82 (d, *J* = 6.9 Hz, 3H), 0.74 (d, *J* = 6.9 Hz, 3H). HRMS (m/z): calculated for $C_{17}H_{23}N_6O_5^+$ $[M+H]^+$ 391.1724; found, 391.1719.

**((2*R*,3*S*,4*R*,5*R*)-5-(4-aminopyrrolo[2,1-*f*][1,2,4]triazin-7-yl)-5-cyano-3,4-dihydroxytetrahydrofuran-2-yl)methyl *L*-valinate (ATV020)**

Compound **3m** was prepared using a procedure analogous to the synthesis of compound **3l** but (*tert*-butoxycarbonyl)-*L*-valine in place of (*tert*-butoxycarbonyl)-*D*-valine. **3m** was obtained as a white solid (2.28 g, 95%).

**ATV020** was prepared using a procedure analogous to the synthesis of **ATV019** but compound **3m** in place of compound **3l**. **ATV020** was obtained as a white solid (0.85 g, 50%). HPLC purity 99.94% (OD-3; eluent, *n*-hexane/isopropanol = 80/20, 0.1% HCOOH; flow rate 0.8 mL/min; temperature 30 °C; wavelength 254 nm; HPLC analysis data are reported in relative area % and were not adjusted to weight %). $^{1}$H NMR (600 MHz, CD3OD) δ 7.76 (s, 1H), 6.80 (d, *J* = 1.6 Hz, 2H), 4.81 (d, *J* = 5.3 Hz, 1H), 4.42-4.26 (m, 3H), 4.04 (t, *J* = 5.8 Hz, 1H), 3.25 (d, *J* = 4.9 Hz, 1H), 1.97-1.84 (m, 1H), 0.83 (d, *J* = 6.9 Hz, 3H), 0.79 (d, *J* = 6.9 Hz, 3H). $^{13}$C NMR (151 MHz, CD3OD) δ 173.76, 155.85, 146.93, 124.12, 116.62, 116.21, 110.86, 101.11, 81.75, 80.16, 74.04, 70.76, 63.66, 59.27,31.62, 17.75, 16.46. HRMS (ESI) of $C_{17}H_{22}N_6O_5$: *m/z*391.1724 $[M+H^+]$. HRMS (m/z): calculated for $C_{17}H_{23}N_6O_5^+$ $[M+H]^+$ 391.1724; found, 391.1718.

**((2*R*,3*S*,4*R*,5*R*)-5-(4-aminopyrrolo[2,1-*f*][1,2,4]triazin-7-yl)-5-cyano-3,4-dihydroxytetrahydrofuran-2-yl)methyl *L*-phenylalaninate (ATV021)**

**ATV021** was prepared using a procedure analogous to the synthesis of **ATV016** but (tert-butoxycarbonyl)-*L*-phenylalanine in place of (*tert*-butoxycarbonyl)-*D*-valine. **ATV021** was obtained as a white solid (0.1 g, 16.9% two-step yield). HPLC purity 99.92% (OD-3; eluent, *n*-hexane/isopropanol = 80/20, 0.1% HCOOH; flow rate 0.8 mL/min; temperature 30 °C; wavelength 254 nm; HPLC analysis data are reported in relative area % and were not adjusted to weight %). $^{1}$H NMR (600 MHz, DMSO-$d_6$) δ (ppm): 7.96 (br, 1H), 7.95 (s, 1H), 7.87 (br, 1H), 7.21-7.13 (m, 5H), 6.93 (d, *J* = 4.5 Hz, 1H), 6.81 (d, *J* = 4.5 Hz, 1H), 6.33 (d, *J* = 6.2 Hz, 1H), 5.36 (br, 1H), 4.70 (t, *J* = 5.0 Hz, 1H), 4.28-4.24 (m, 2H), 4.19-4.16 (m, 1H), 3.88 (t, *J* = 5.5 Hz, 1H), 3.57 (t, *J* = 6.7 Hz, 1H), 2.84-2.73 (m, 2H), 1.85 (br, 2H). $^{13}$C NMR (150 MHz, DMSO-$d_6$) δ (ppm): 174.5, 155.4, 147.8, 137.5, 129.0, 127.9, 126.1, 123.4, 116.8, 116.4, 110.1, 100.7, 81.1, 78.9, 73.8, 70.0, 63.1, 55.6, 40.4. HRMS (m/z): calculated for $C_{21}H_{23}N_6O_5^+$ $[M+H]^+$ 439.1724; found, 439.1719.

**((2R,3S,4R,5R)-5-(4-aminopyrrolo[2,1-f][1,2,4]triazin-7-yl)-5-cyano-3,4-dihydroxytetrahydrofuran-2-yl)methyl D-phenylalaninate (ATV022)**

**ATV022** was prepared using a procedure analogous to the synthesis of **ATV016** but (tert-butoxycarbonyl)-*D*-phenylalanine in place of (*tert*-butoxycarbonyl)-*D*-valine. **ATV022** was obtained as a white solid (0.1 g, 15.3% two-step yield). HPLC purity 98.75% (OD-3; eluent, *n*-hexane/isopropanol = 80/20, 0.1% HCOOH; flow rate 0.8 mL/min; temperature 30 °C; wavelength 254 nm; HPLC analysis data are reported in relative area % and were not adjusted to weight %). $^{1}$H NMR (600 MHz, DMSO-$d_6$) δ (ppm): 7.92 (s, 1H), 7.85 (br, 1H), 7.25-7.14 (m, 5H), 6.90 (d, *J* = 4.5 Hz, 1H), 6.80 (d, *J* = 4.5 Hz, 1H), 6.33 (d, *J* = 5.9 Hz, 1H), 5.39 (d, *J* = 5.6 Hz,1H), 4.71 (t, *J* = 5.3 Hz, 1H), 4.25-4.17 (m, 3H), 3.95-3.94 (m, 1H), 3.56 (t, *J* = 6.7 Hz,1H), 2.86-2.71 (m, 2H), 1.75 (br, 2H). $^{13}$C NMR (150 MHz, DMSO-$d_6$) δ (ppm): 175.2, 156.1, 148.4, 138.2, 129.7, 128.6, 126.8, 124.0, 117.4, 117.1, 110.8, 101.3, 81.7, 79.5, 74.5, 70.7, 63.9, 56.1. HRMS (m/z): calculated for $C_{21}H_{23}N_6O_5^+$ $[M+H]^+$ 439.1724; found, 439.1719.

**((2*R*,3*S*,4*R*,5*R*)-5-(4-aminopyrrolo[2,1-*f*][1,2,4]triazin-7-yl)-5-cyano-3,4-dihydroxytetrahydrofuran-2-yl)methyl (2*S*)-2-amino-3-methylpentanoate (ATV023)**

**ATV023** was prepared using a procedure analogous to the synthesis of **ATV016** but (2*S*)-2-((tert-butoxycarbonyl)amino)-3-methylpentanoic acid in place of (*tert*-butoxycarbonyl)-*D*-valine. **ATV023** was obtained as a white solid (0.06 g, 10.2% two-step yield). HPLC purity 95.73% (OD-3; eluent, *n*-hexane/isopropanol = 80/20; flow rate 0.8 mL/min; temperature 30 °C; wavelength 254 nm; HPLC analysis data are reported in relative area % and were not adjusted to weight %). $^1$H NMR (600 MHz, DMSO-$d_6$) δ (ppm): 7.95 (br, 1H), 7.92 (s, 1H), 7.87 (br, 1H), 6.92 (d, *J* = 5.8 Hz, 1H), 6.83 (d, *J* = 5.8 Hz, 1H), 6.35 (br, 1H), 5.40 (br, 1H), 4.73 (d, *J* = 4.6 Hz, 1H), 4.29-4.24 (m, 3H), 3.96 (t, *J* = 5.0 Hz, 1H), 3.18 (d, *J* = 4.2 Hz, 1H), 1.53-1.51 (m, 1H), 1.39-1.32 (m, 1H), 1.11-1.04 (m, 1H), 0.80-0.74 (m, 6H). $^{13}$C NMR (150 MHz, DMSO-$d_6$) δ (ppm): 175.6, 156.1, 148.4, 124.0, 117.4, 117.0, 110.8, 101.3, 81.6, 79.5, 74.5, 70.7, 63.5, 59.1, 39.1, 24.6, 16.0, 11.8. HRMS (m/z): calculated for $C_{18}H_{25}N_6O_5^+$ [M+H]$^+$ 405.1880; found, 405.1874.

**((2*R*,3*S*,4*R*,5*R*)-5-(4-aminopyrrolo[2,1-*f*][1,2,4]triazin-7-yl)-5-cyano-3,4-dihydroxytetrahydrofuran-2-yl)methyl (2*R*)-2-amino-3-methylpentanoate (ATV024)**

**ATV024** was prepared using a procedure analogous to the synthesis of **ATV016** but (2*R*)-2-((tert-butoxycarbonyl)amino)-3-methylpentanoic acid in place of (*tert*-butoxycarbonyl)-*D*-valine. **ATV024** was obtained as a white solid (0.06 g, 9.1% two-step yield). HPLC purity 93.39% (OD-3; eluent, *n*-hexane/isopropanol = 80/20; flow rate 0.8 mL/min; temperature 30 °C; wavelength 254 nm; HPLC analysis data are reported in relative area % and were not adjusted to weight %). $^1$H NMR (600 MHz, DMSO-$d_6$) δ (ppm): 7.92 (s, 1H), 7.86 (br, 2H), 6.92 (d, *J* = 5.8 Hz, 1H), 6.83 (d, *J* = 5.8 Hz, 1H), 6.33 (d, *J* = 4.7 Hz, 1H), 5.39 (br, 1H), 4.71 (br, 1H), 4.30-4.19 (m, 3H), 3.97 (t, *J* = 5.1 Hz, 1H), 3.15 (d, *J* = 5.3 Hz, 1H), 1.53-1.50 (m, 1H), 1.39-1.34 (m, 1H), 1.11-1.04 (m, 1H), 0.80-0.75 (m, 6H). $^{13}$C NMR (150 MHz, DMSO-$d_6$) δ (ppm): 175.6, 156.1, 148.4, 124.0, 117.4, 117.1, 110.8, 101.3, 81.7, 79.5, 74.5, 70.8, 63.8, 59.1, 39.0, 24.6, 16.1, 11.8. HRMS (m/z): calculated for $C_{18}H_{25}N_6O_5^+$ [M+H]$^+$ 405.1880; found, 405.1875.

### [1]H, [13]C NMR, HRMS and HPLC spectra

#### [1]H, [13]C NMR, HRMS and HPLC spectrum of ATV001

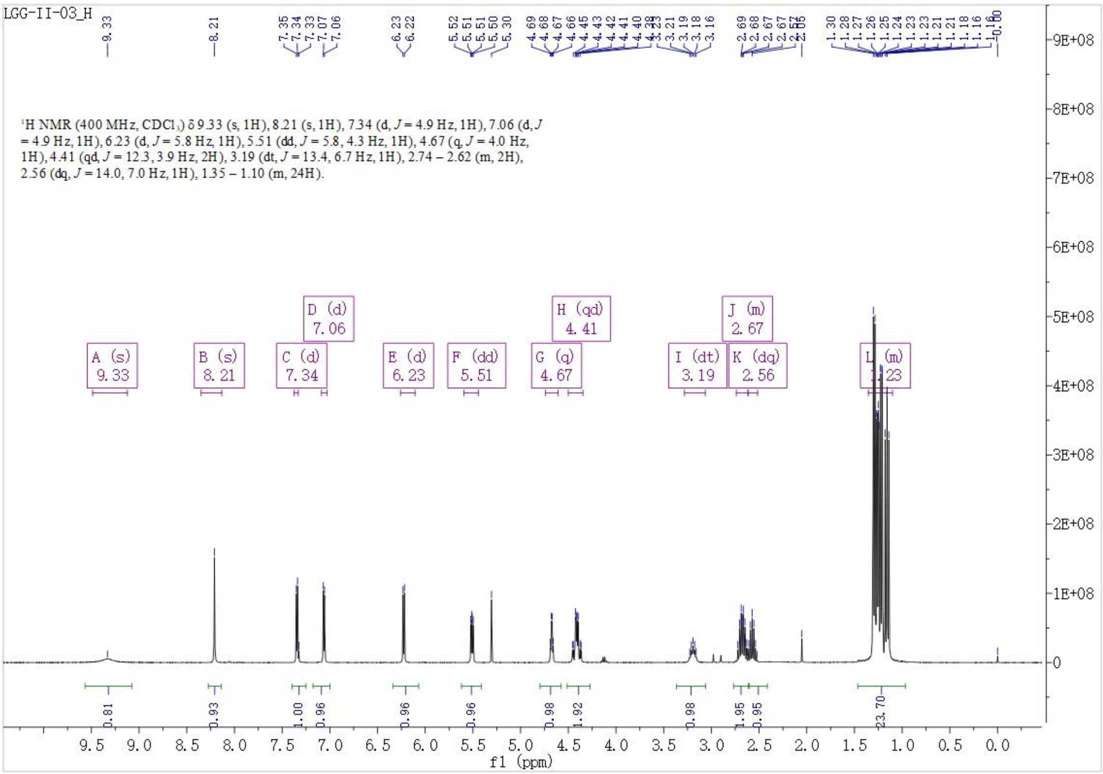

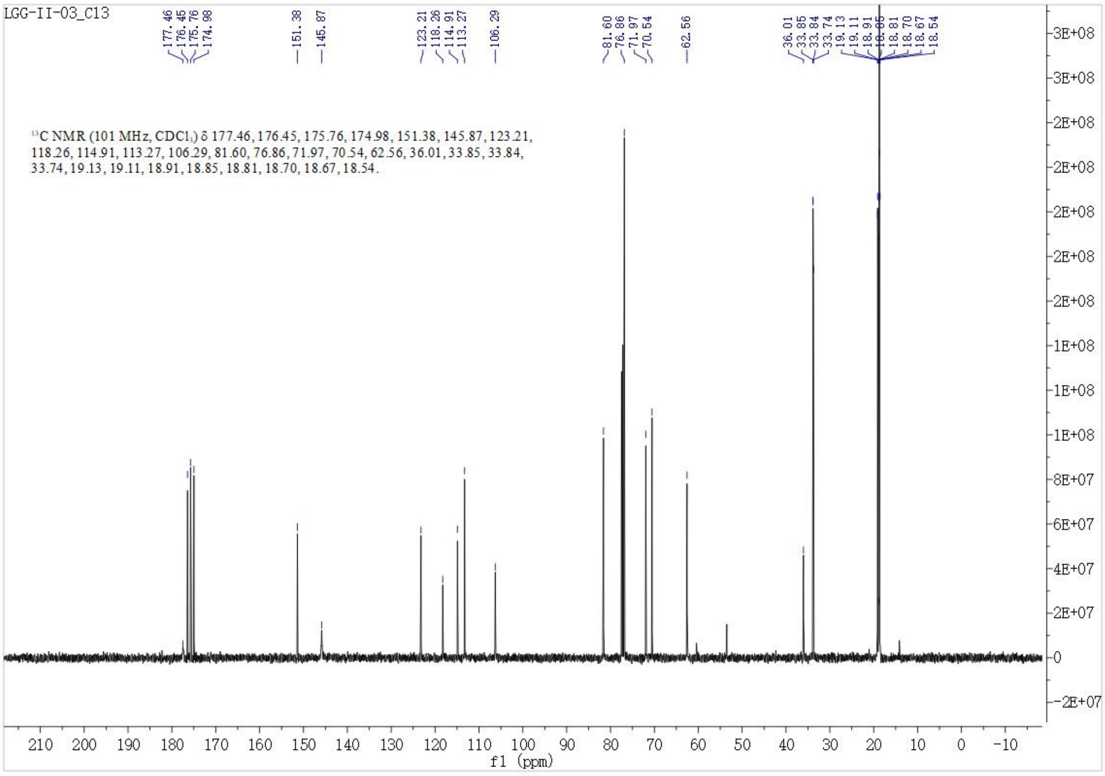

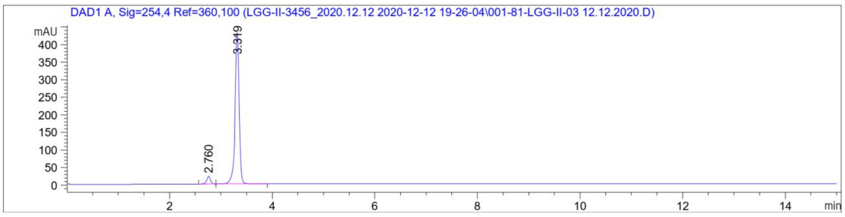

Signal 1: DAD1 A, Sig=254,4 Ref=360,100

| Peak # | RetTime [min] | Type | Width [min] | Area [mAU*s] | Height [mAU] | Area % |
|---|---|---|---|---|---|---|
| 1 | 2.760 | BB | 0.0715 | 103.42430 | 21.72923 | 4.1738 |
| 2 | 3.319 | BB | 0.0843 | 2374.49878 | 430.64349 | 95.8262 |
| Totals : | | | | 2477.92308 | 452.37272 | |

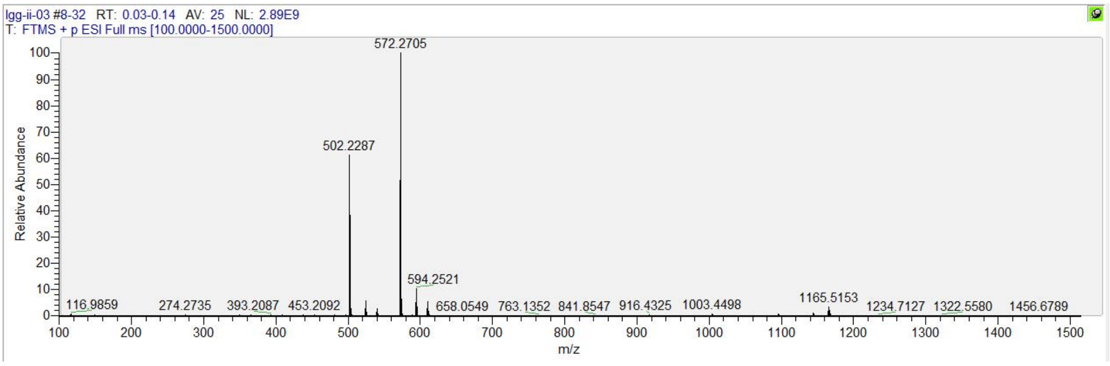

#### $^1$H, $^{13}$C NMR, HRMS and HPLC spectrum of ATV002

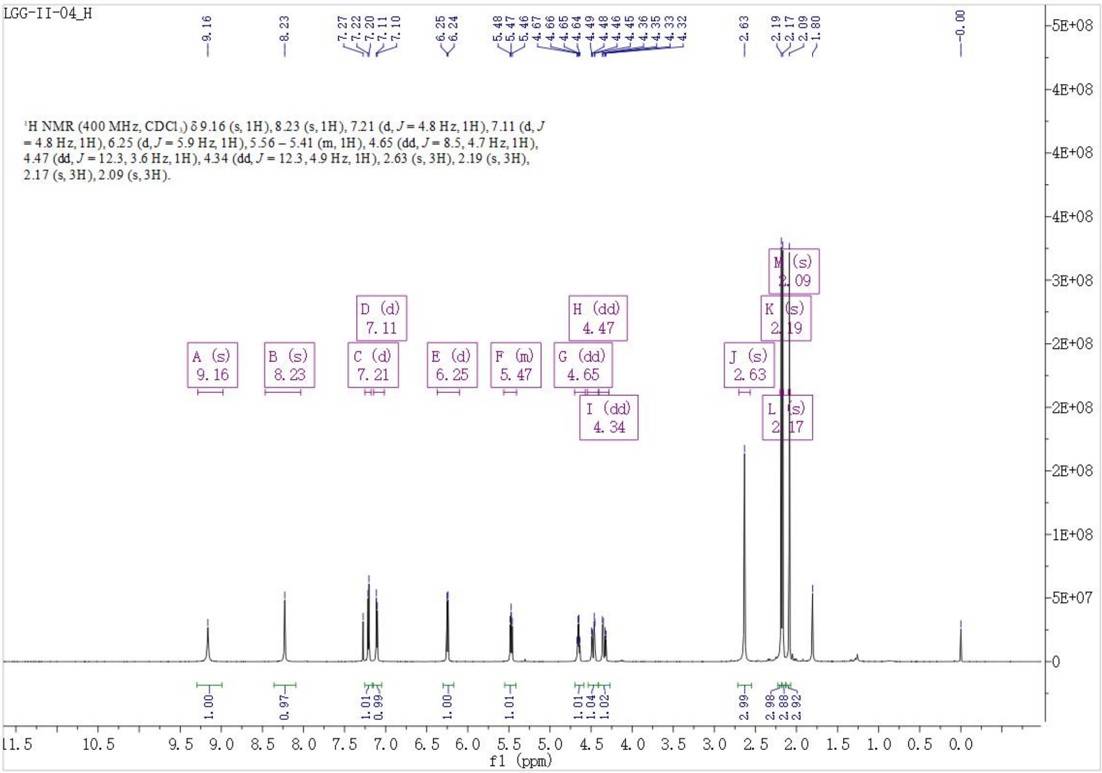

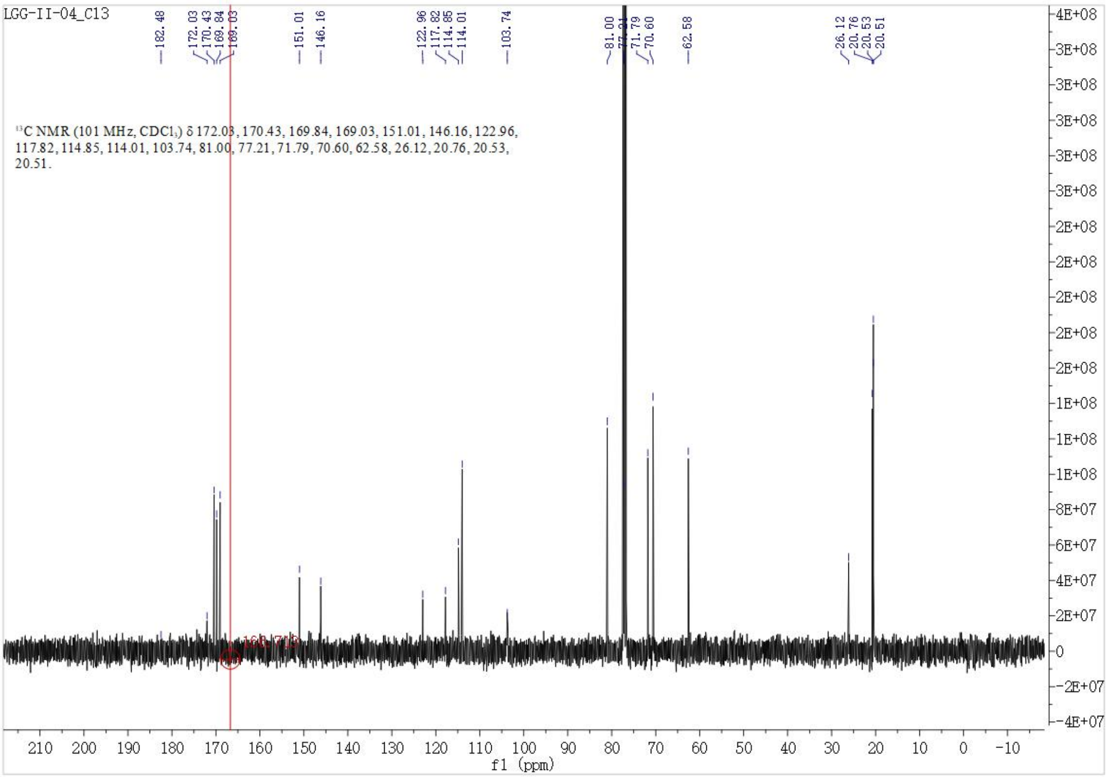

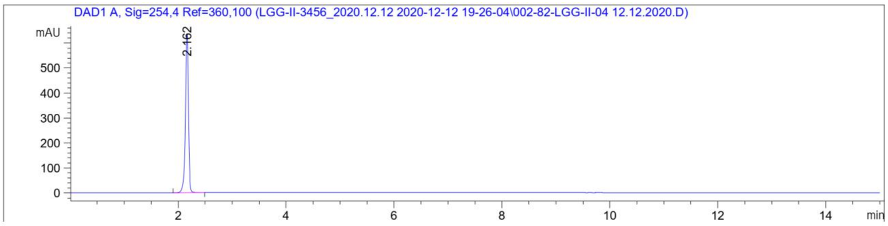

Signal 1: DAD1 A, Sig=254,4 Ref=360,100

| Peak # | RetTime [min] | Type | Width [min] | Area [mAU*s] | Height [mAU] | Area % |
|---|---|---|---|---|---|---|
| 1 | 2.162 | BB | 0.0585 | 2442.95532 | 639.94434 | 100.0000 |

Totals : 2442.95532 639.94434

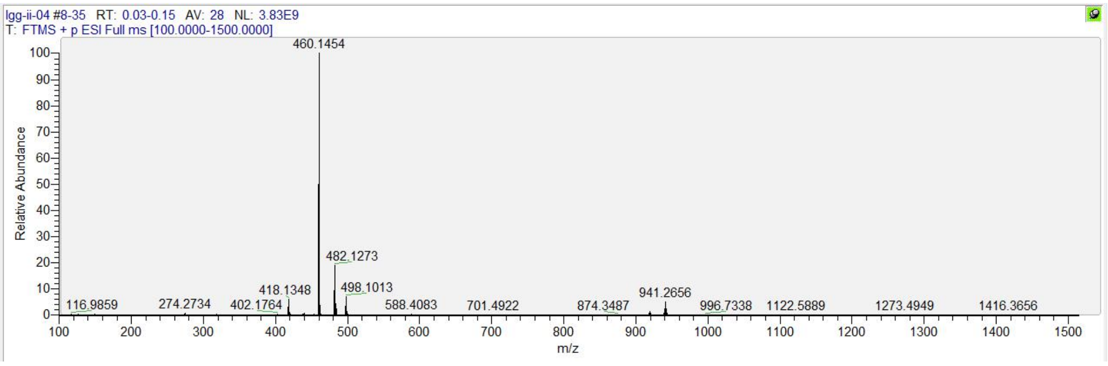

**[1]H, [13]C NMR, HRMS and HPLC spectrum of ATV003**

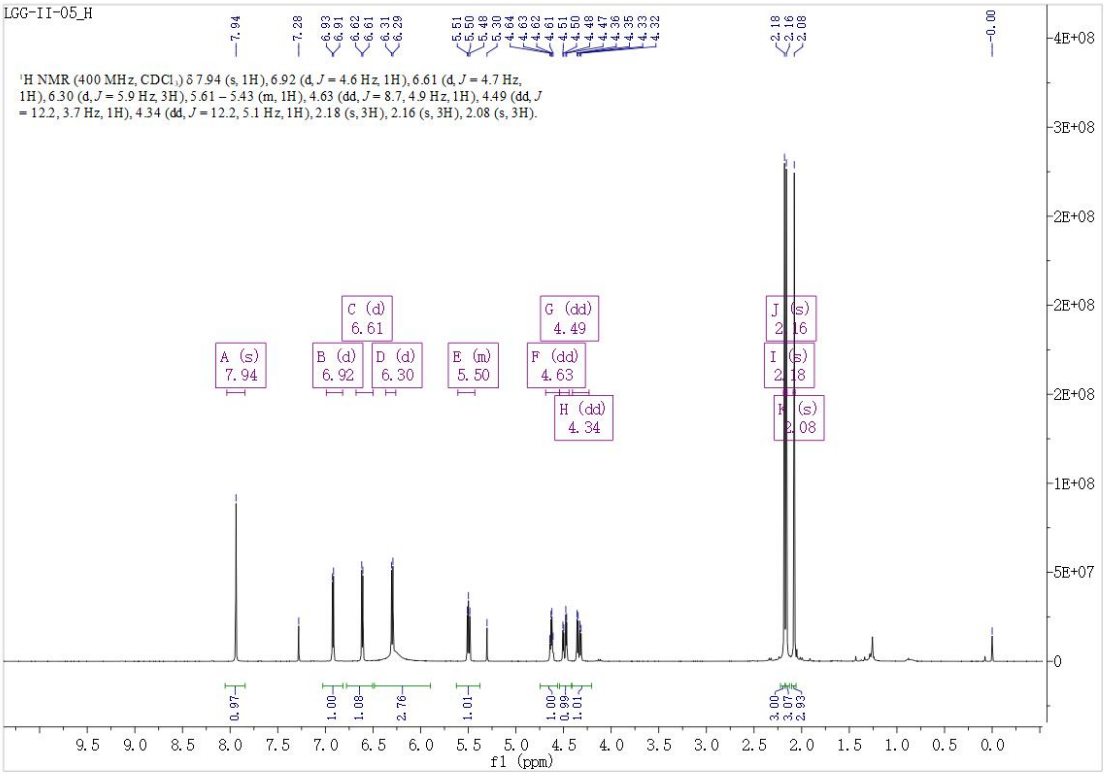

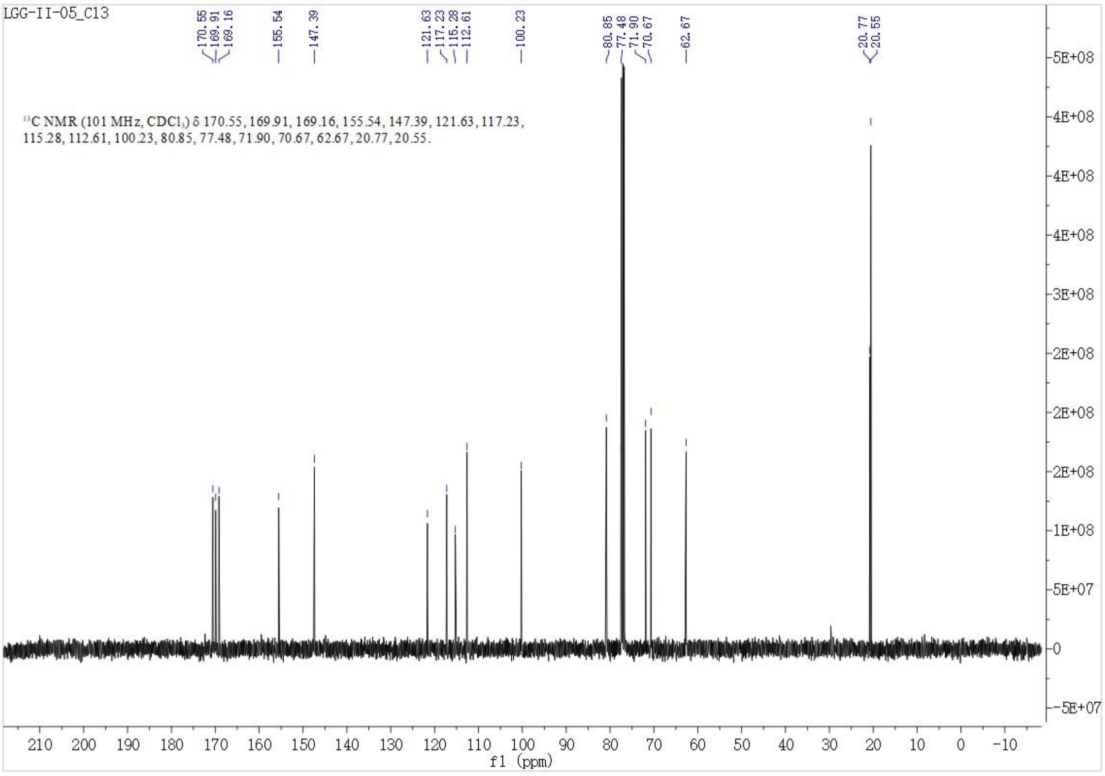

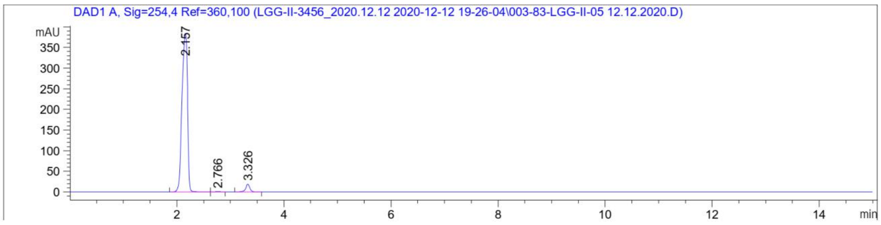

Signal 1: DAD1 A, Sig=254,4 Ref=360,100

| Peak # | RetTime [min] | Type | Width [min] | Area [mAU*s] | Height [mAU] | Area % |
|---|---|---|---|---|---|---|
| 1 | 2.157 | BB | 0.1218 | 2756.41626 | 383.72623 | 96.1023 |
| 2 | 2.766 | BB | 0.0745 | 6.15516 | 1.22679 | 0.2146 |
| 3 | 3.326 | BB | 0.0851 | 105.64048 | 18.94467 | 3.6831 |

Totals : 2868.21190 403.89768

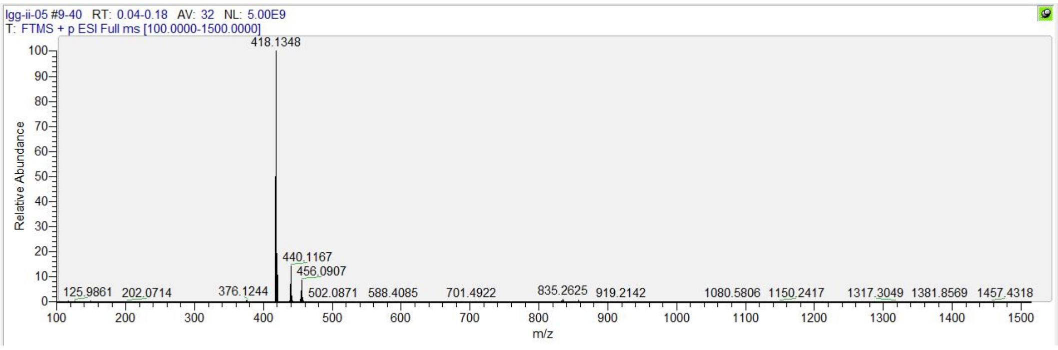

#### [1]H, [13]C NMR, HRMS and HPLC spectrum of ATV004

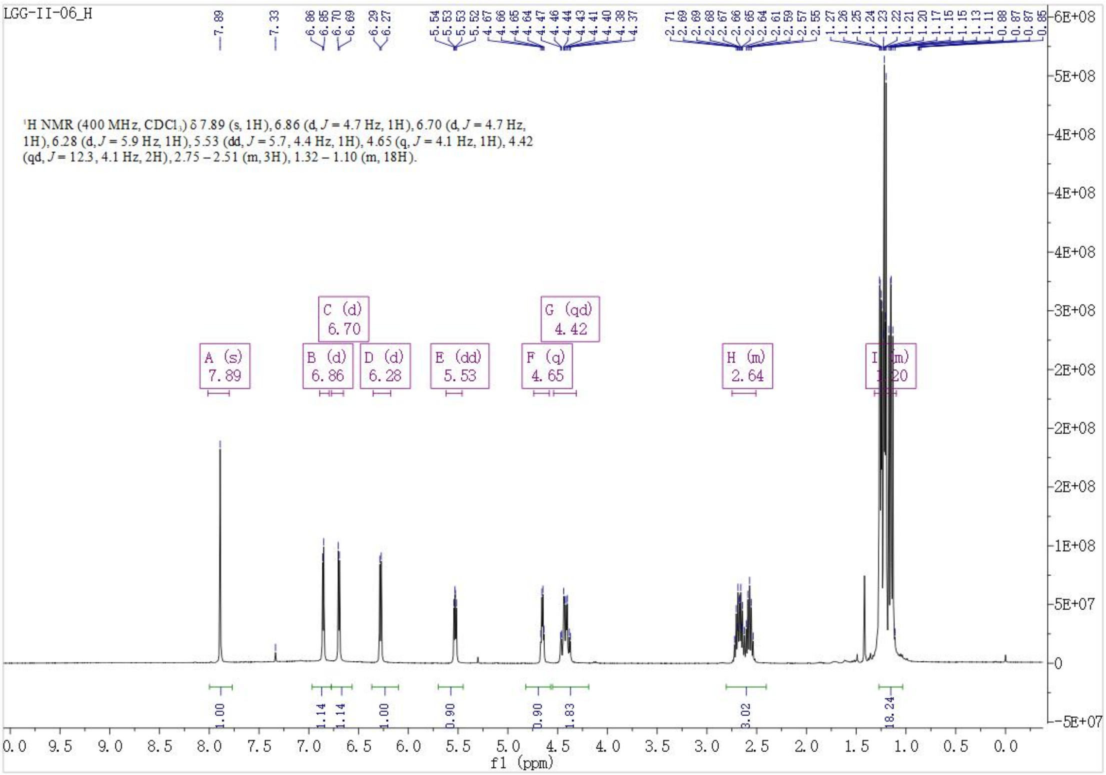

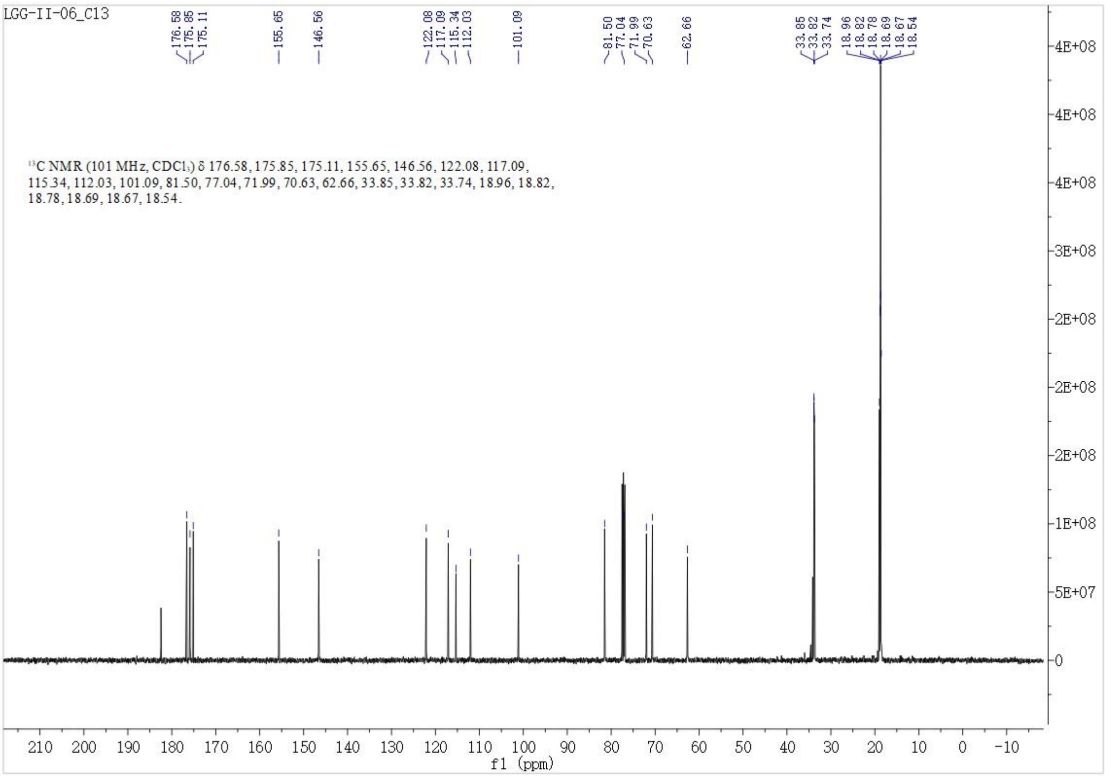

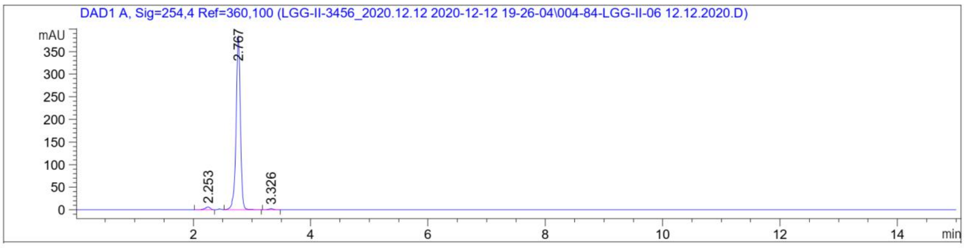

Signal 1: DAD1 A, Sig=254,4 Ref=360,100

| Peak # | RetTime [min] | Type | Width [min] | Area [mAU*s] | Height [mAU] | Area % |
|---|---|---|---|---|---|---|
| 1 | 2.253 | BB | 0.0916 | 36.86055 | 6.01004 | 1.8890 |
| 2 | 2.767 | BB | 0.0760 | 1902.37317 | 382.57452 | 97.4925 |
| 3 | 3.326 | BB | 0.0828 | 12.06856 | 2.24097 | 0.6185 |
| Totals : | | | | 1951.30228 | 390.82553 | |

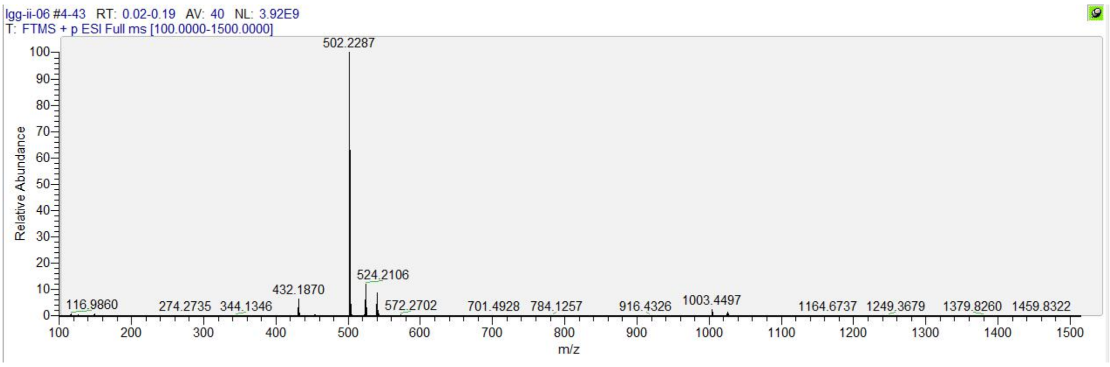

**[1]H, [13]C NMR, HRMS and HPLC spectrum of ATV007**

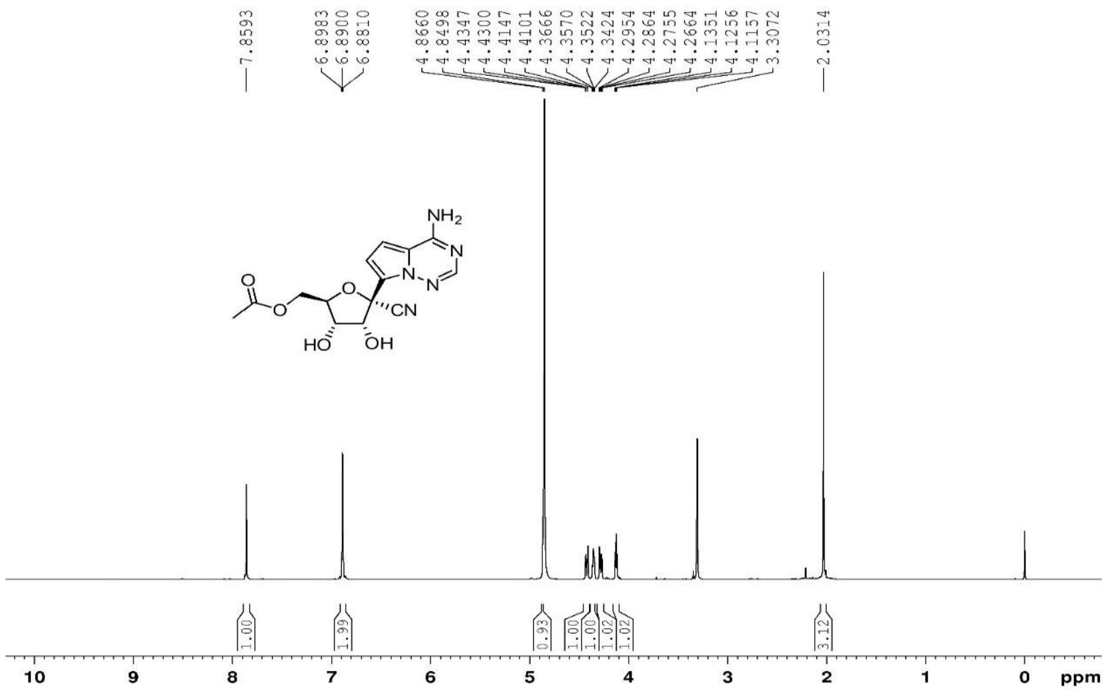

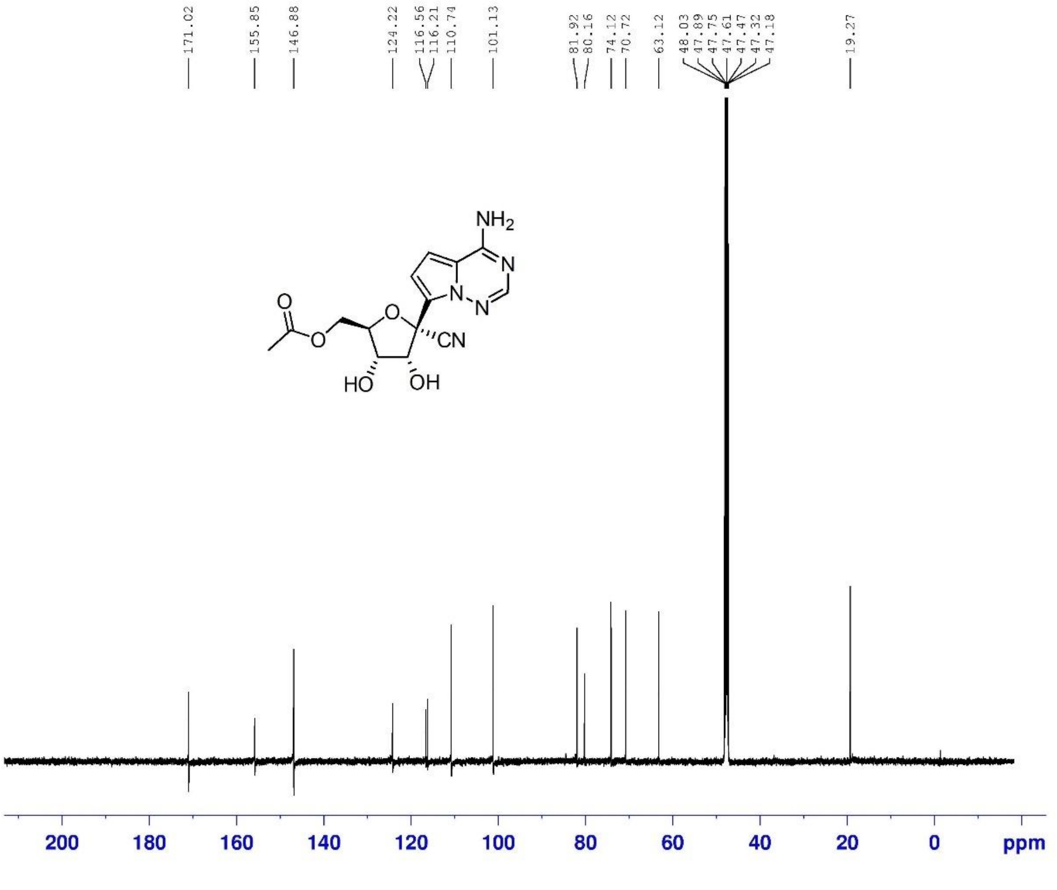

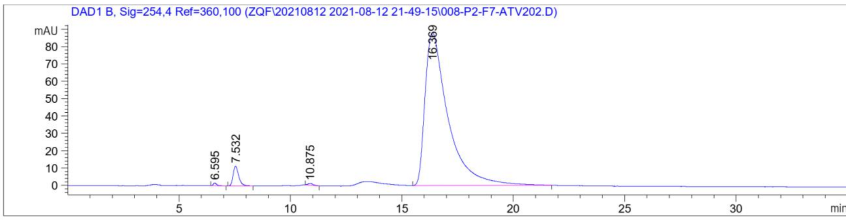

Signal 3: DAD1 B, Sig=254,4 Ref=360,100

| Peak # | RetTime [min] | Type | Width [min] | Area [mAU*s] | Height [mAU] | Area % |
|---|---|---|---|---|---|---|
| 1 | 6.595 | BB | 0.1525 | 16.13007 | 1.58786 | 0.2463 |
| 2 | 7.532 | BB | 0.2712 | 200.50430 | 11.39699 | 3.0611 |
| 3 | 10.875 | BB | 0.2062 | 15.10019 | 1.07296 | 0.2305 |
| 4 | 16.369 | BB | 1.0297 | 6318.23682 | 88.21001 | 96.4621 |

Totals : 6549.97138 102.26782

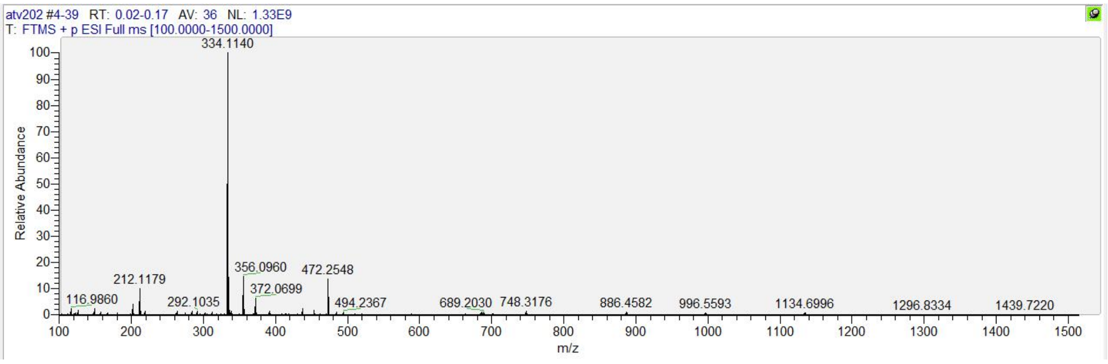

#### [1]H, [13]C NMR, HRMS and HPLC spectrum of ATV006

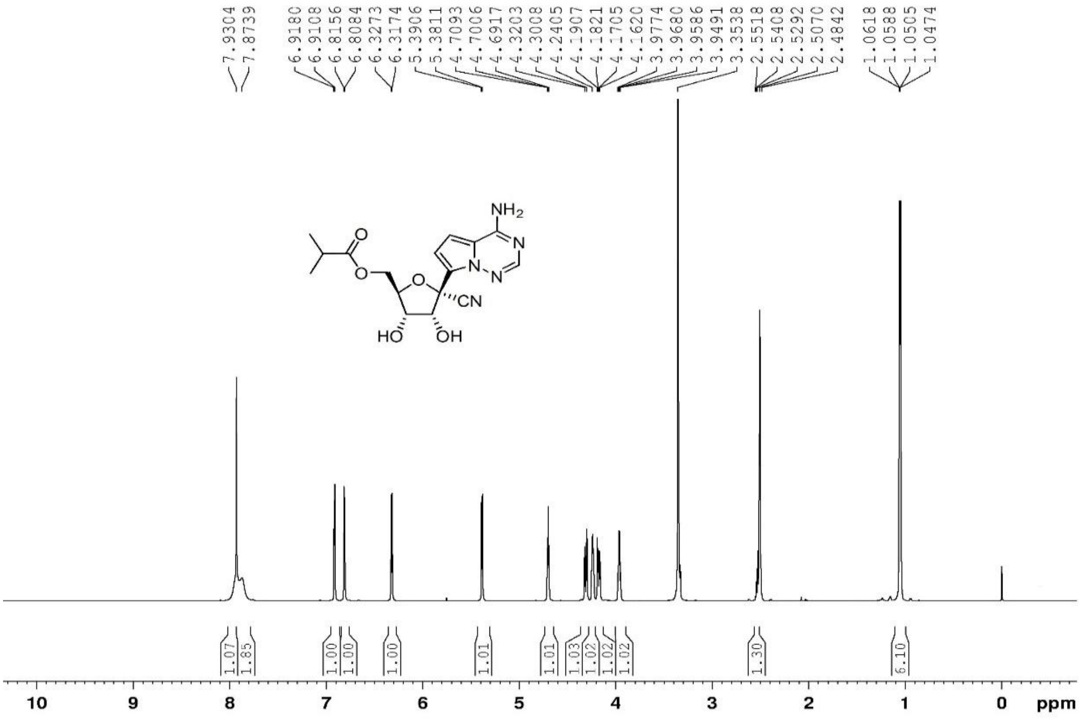

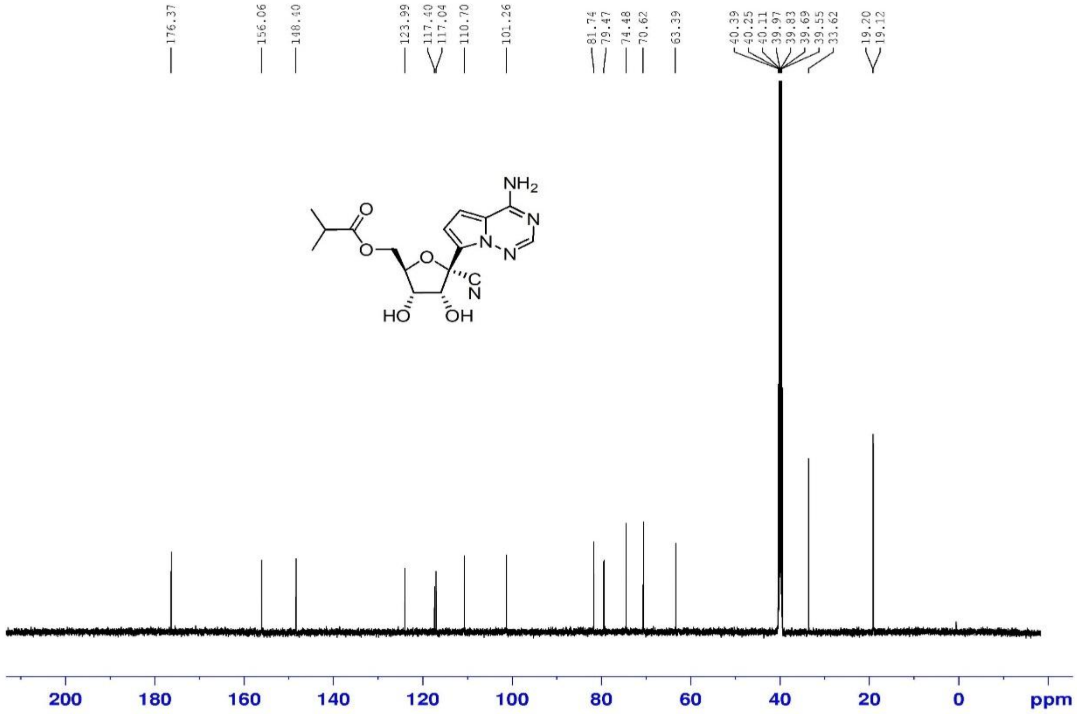

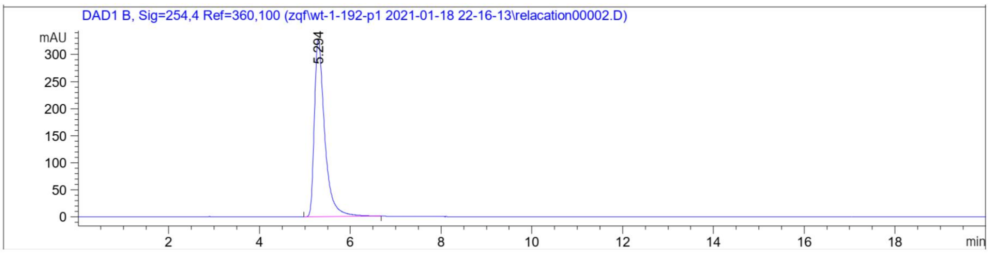

Signal 2: DAD1 B, Sig=254,4 Ref=360,100

| Peak # | RetTime [min] | Type | Width [min] | Area [mAU*s] | Height [mAU] | Area % |
|---|---|---|---|---|---|---|
| 1 | 5.294 | BB | 0.2370 | 5198.32275 | 327.79962 | 100.0000 |

| | | | | | |
|---|---|---|---|---|---|
| Totals : | | | | 5198.32275 | 327.79962 |

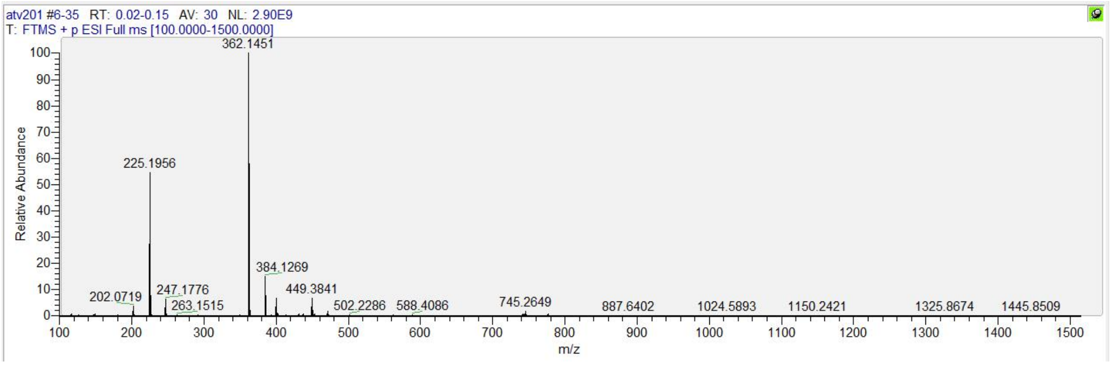

**[1]H, [13]C NMR, HRMS and HPLC spectrum of ATV008**

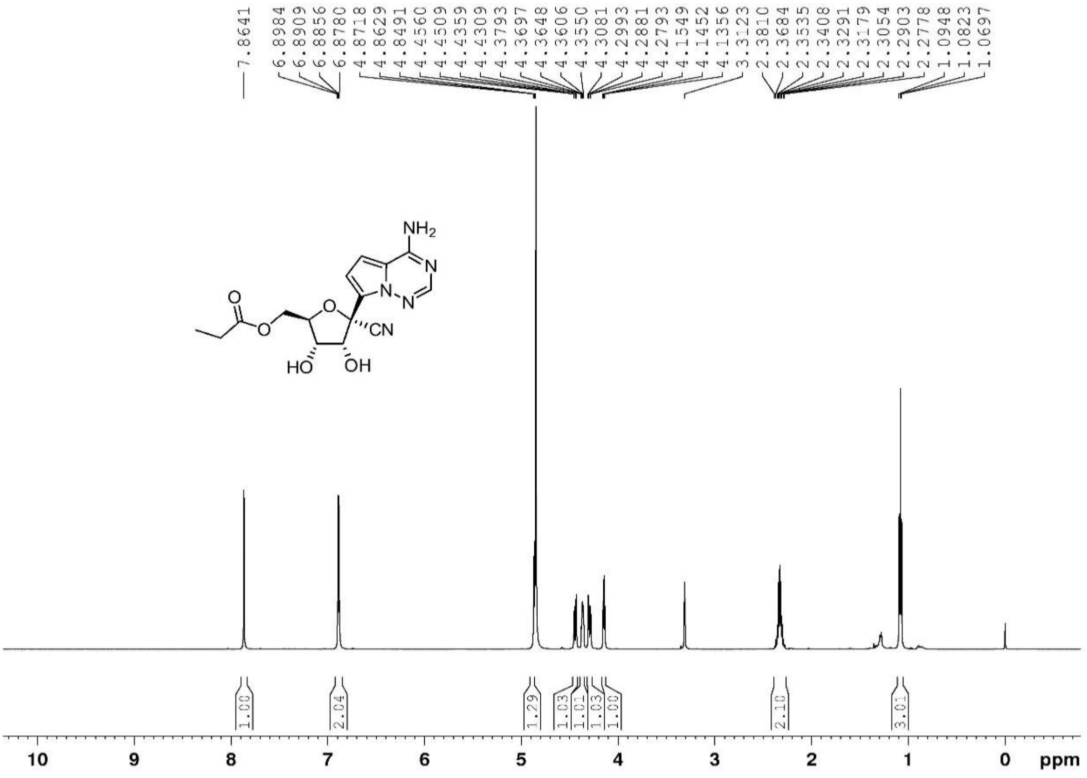

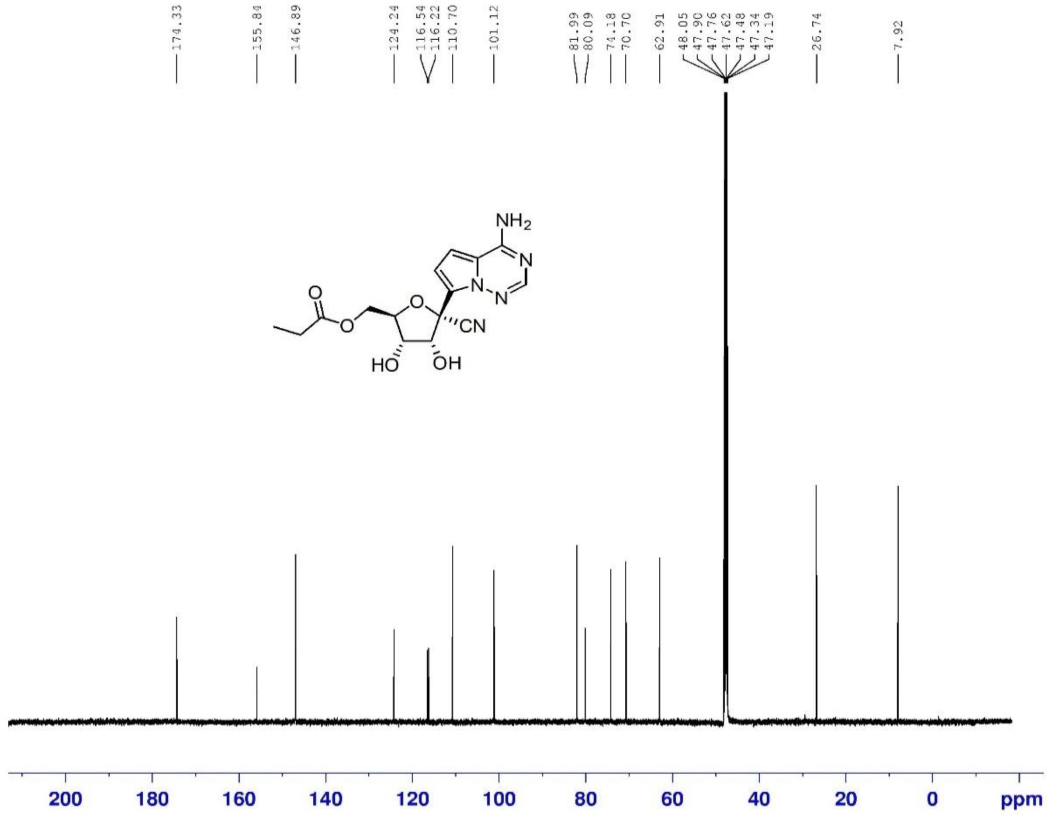

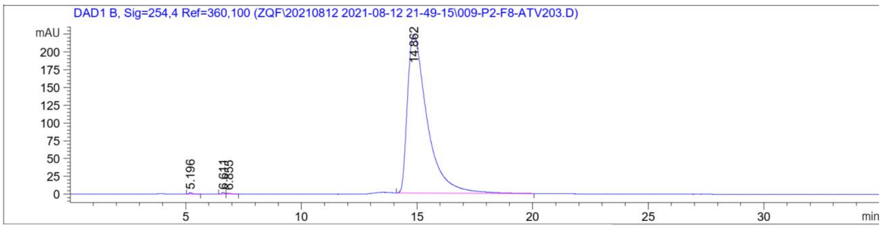

Signal 3: DAD1 B, Sig=254,4 Ref=360,100

| Peak # | RetTime [min] | Type | Width [min] | Area [mAU*s] | Height [mAU] | Area % |
|---|---|---|---|---|---|---|
| 1 | 5.196 | BB | 0.1433 | 19.22932 | 2.05232 | 0.1451 |
| 2 | 6.611 | BV | 0.1386 | 20.92410 | 2.24794 | 0.1579 |
| 3 | 6.855 | VB | 0.2011 | 17.74485 | 1.25372 | 0.1339 |
| 4 | 14.862 | BB | 0.8724 | 1.31976e4 | 223.02089 | 99.5632 |

Totals : 1.32555e4 228.57487

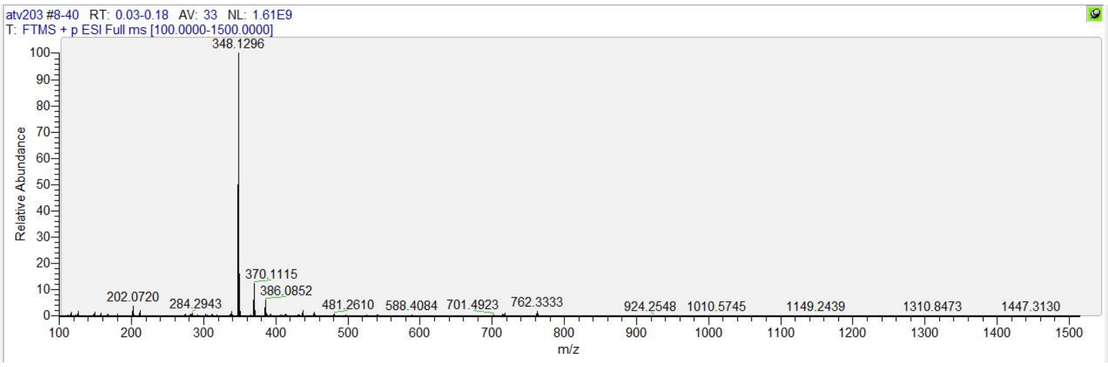

#### [1]H, [13]C NMR, HRMS and HPLC spectrum of ATV009

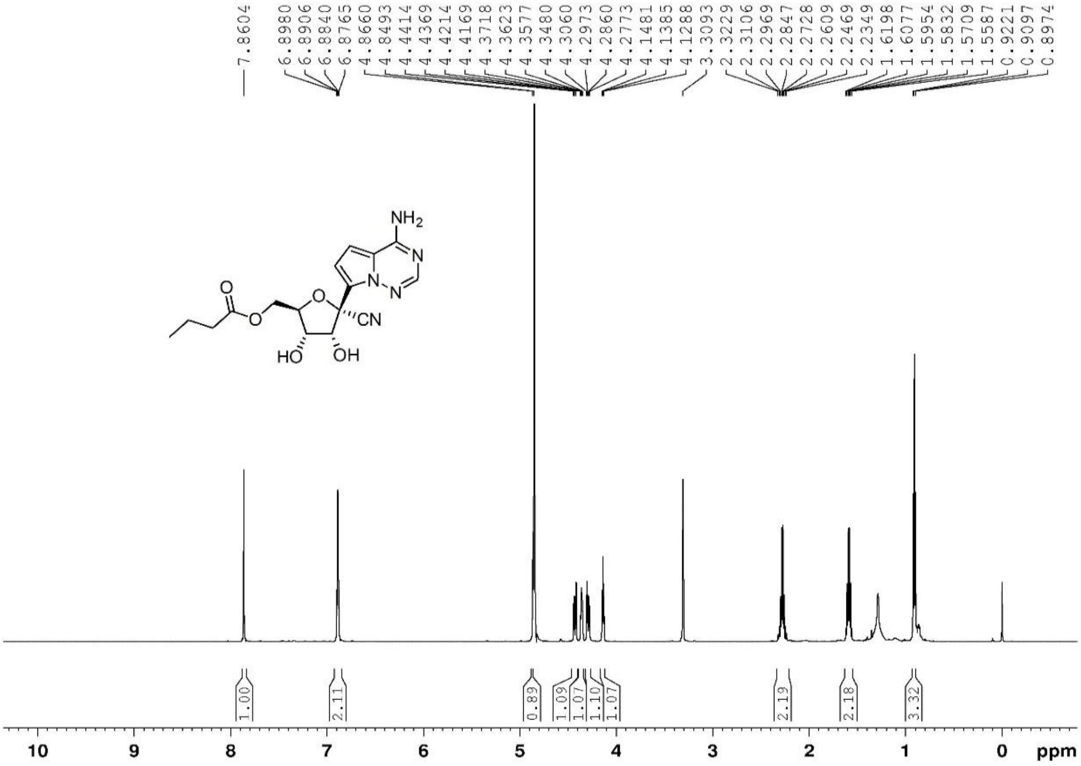

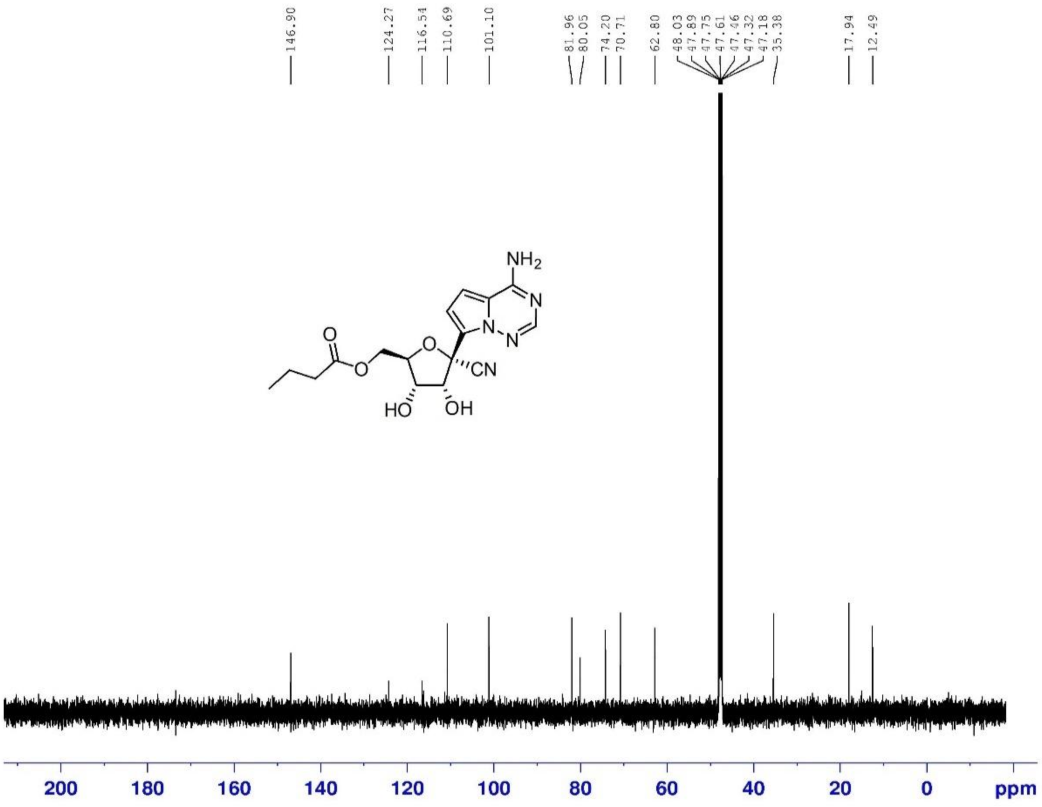

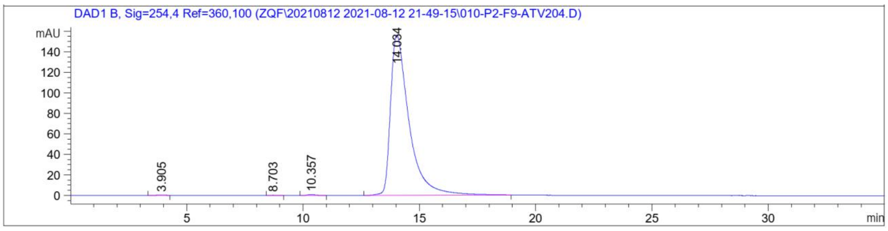

Signal 3: DAD1 B, Sig=254,4 Ref=360,100

| Peak # | RetTime [min] | Type | Width [min] | Area [mAU*s] | Height [mAU] | Area % |
|---|---|---|---|---|---|---|
| 1 | 3.905 | BB | 0.3579 | 14.00595 | 5.17198e-1 | 0.1612 |
| 2 | 8.703 | BB | 0.2424 | 5.38031 | 2.79687e-1 | 0.0619 |
| 3 | 10.357 | BB | 0.3429 | 22.46304 | 8.94659e-1 | 0.2585 |
| 4 | 14.034 | BB | 0.8080 | 8647.78320 | 156.18123 | 99.5184 |

Totals : 8689.63250 157.87277

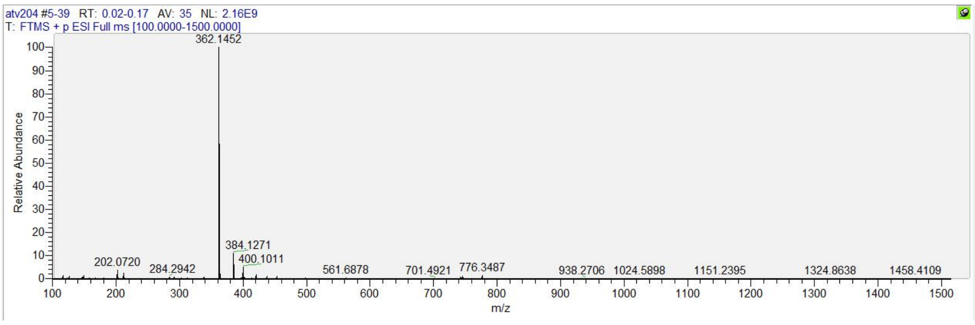

**[1]H, [13]C NMR, HRMS and HPLC spectrum of ATV010**

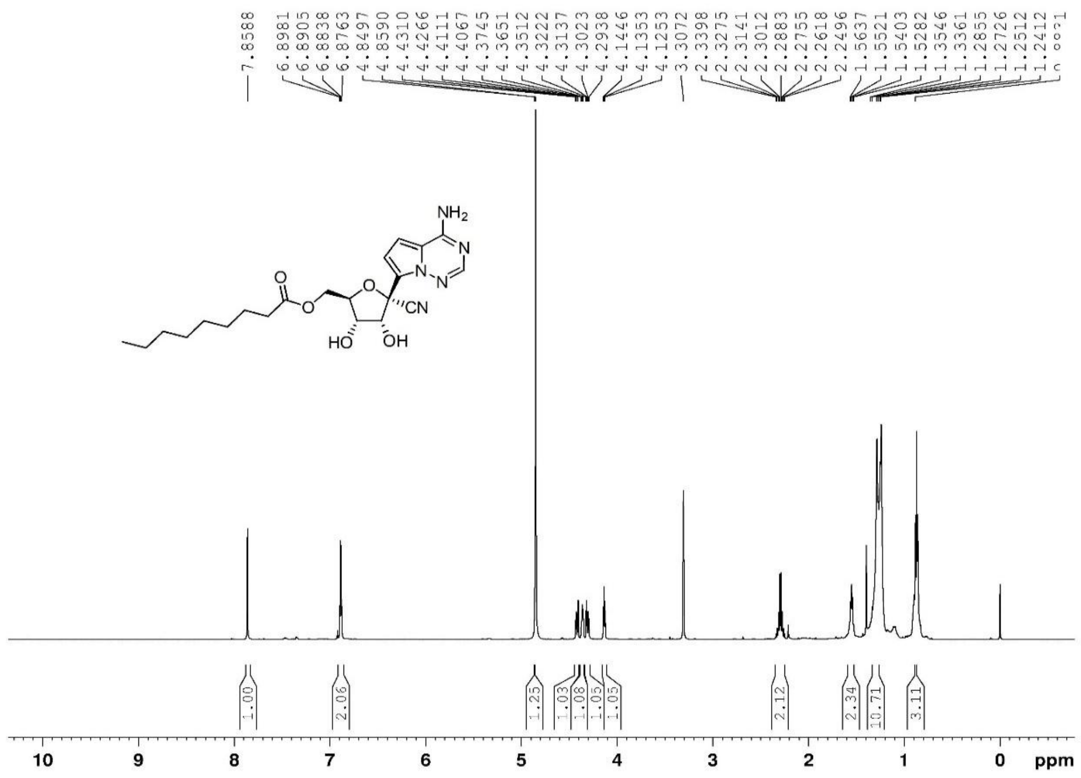

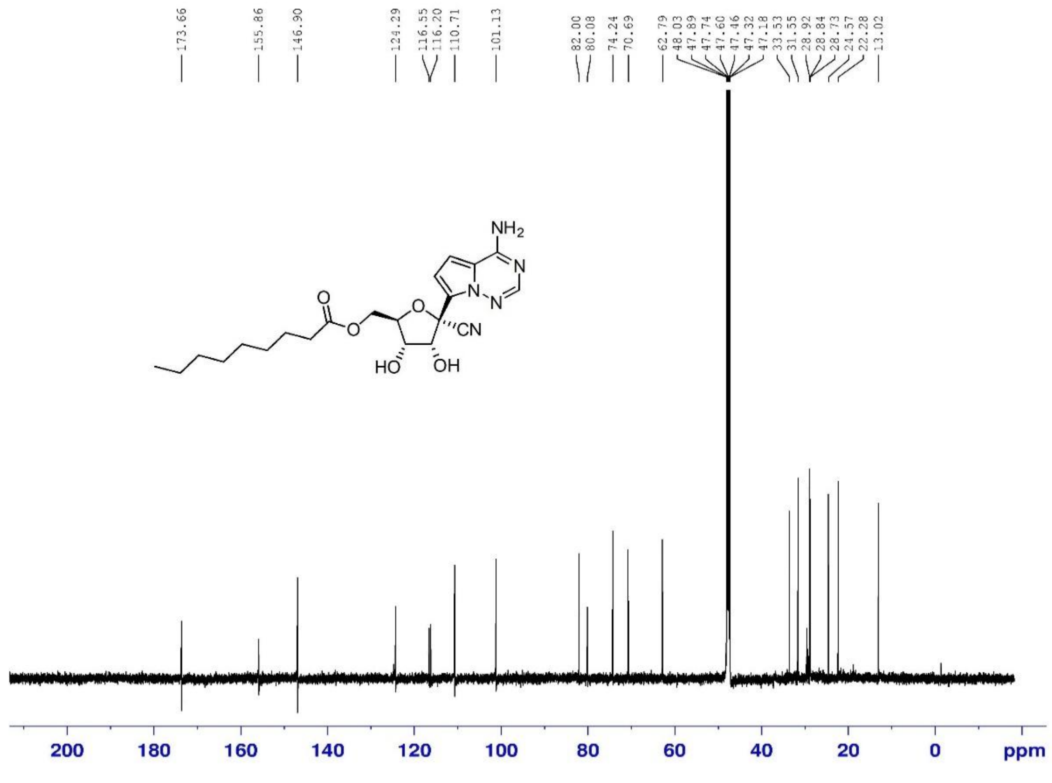

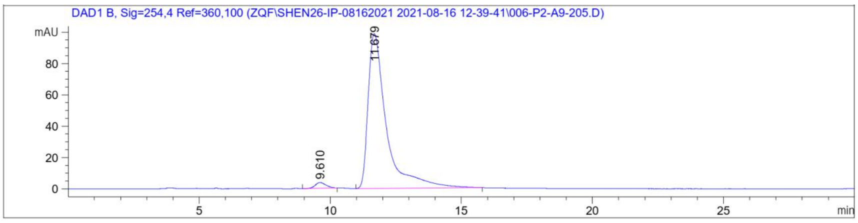

Signal 3: DAD1 B, Sig=254,4 Ref=360,100

| Peak # | RetTime [min] | Type | Width [min] | Area [mAU*s] | Height [mAU] | Area % |
|---|---|---|---|---|---|---|
| 1 | 9.610 | BB | 0.4016 | 105.78338 | 3.67771 | 2.1557 |
| 2 | 11.679 | BB | 0.7137 | 4801.44385 | 98.67751 | 97.8443 |

| | | |
|---|---|---|
| Totals : | 4907.22723 | 102.35522 |

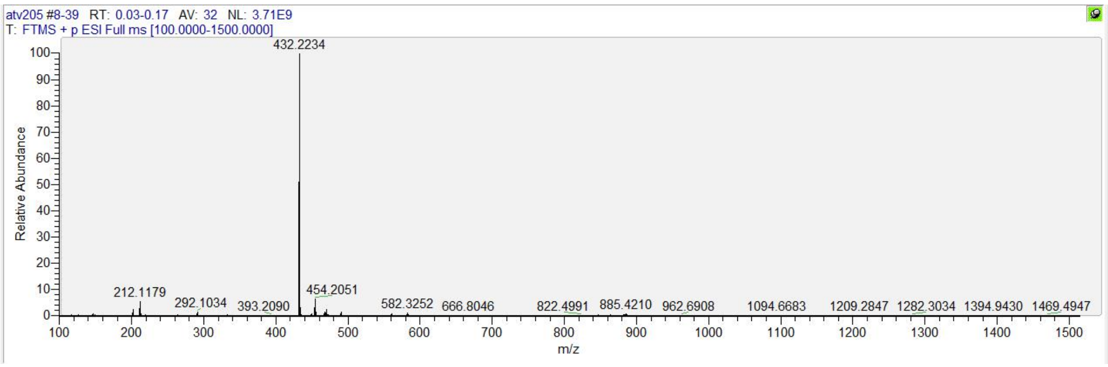

**$^{1}H$, $^{13}C$ NMR, HRMS and HPLC spectrum of ATV011**

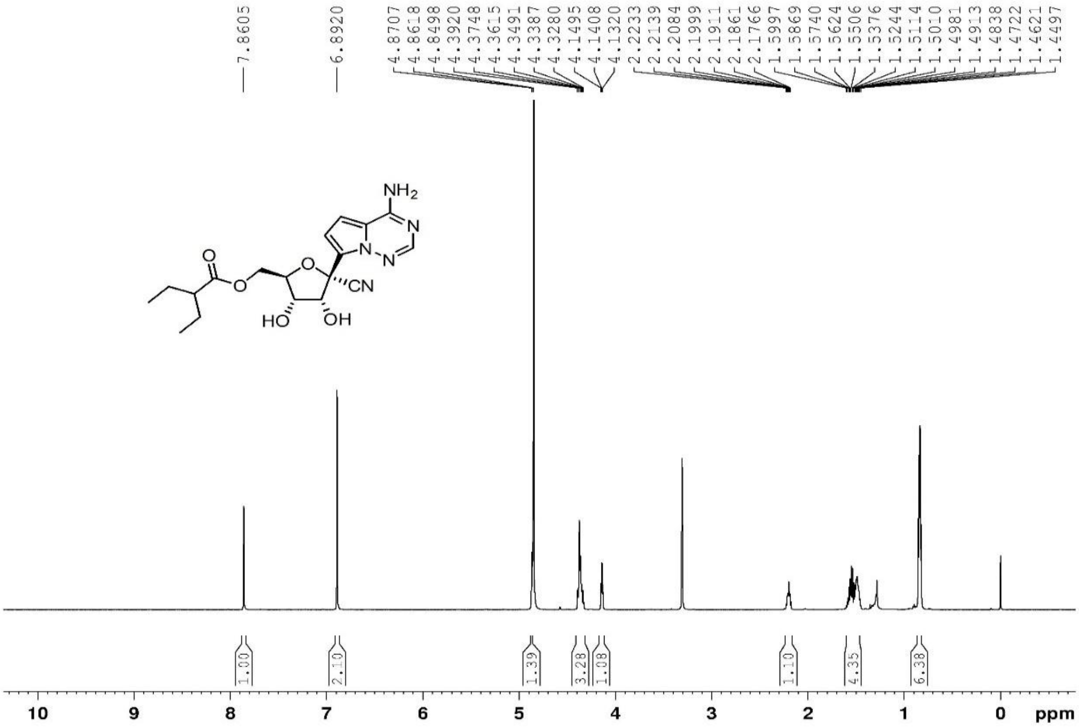

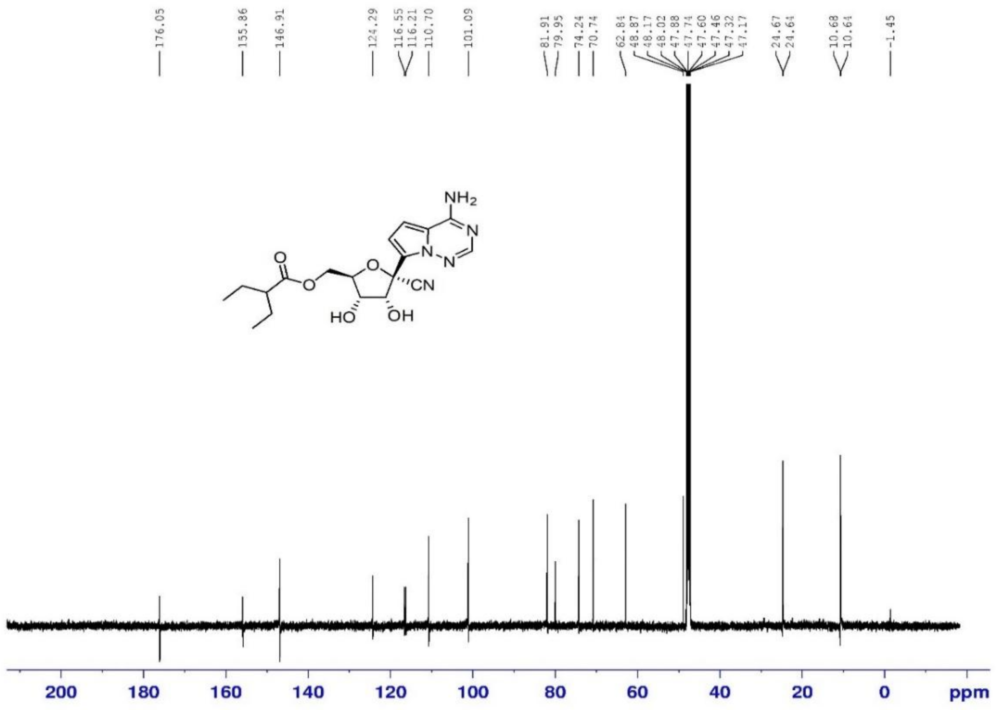

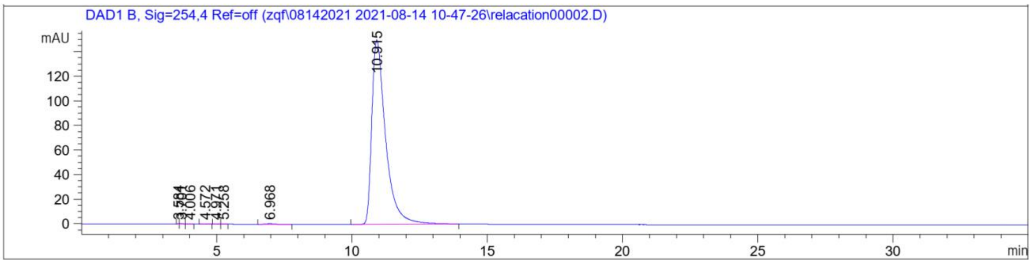

Signal 2: DAD1 B, Sig=254,4 Ref=off

| Peak # | RetTime [min] | Type | Width [min] | Area [mAU*s] | Height [mAU] | Area % |
|---|---|---|---|---|---|---|
| 1 | 3.584 | BV | 0.0649 | 1.48126 | 3.53152e-1 | 0.0278 |
| 2 | 3.701 | VB | 0.1210 | 3.29319 | 3.52413e-1 | 0.0618 |
| 3 | 4.006 | BB | 0.1087 | 8.42285e-1 | 1.16037e-1 | 0.0158 |
| 4 | 4.572 | BB | 0.1491 | 6.01578e-1 | 6.09889e-2 | 0.0113 |
| 5 | 4.971 | BV | 0.1159 | 1.18185 | 1.53305e-1 | 0.0222 |
| 6 | 5.258 | VB | 0.1227 | 5.67738e-1 | 6.20536e-2 | 0.0107 |
| 7 | 6.968 | BB | 0.2874 | 12.94937 | 6.05729e-1 | 0.2432 |
| 8 | 10.915 | BB | 0.5284 | 5304.64160 | 150.05687 | 99.6072 |

Totals : 5325.55888 151.76055

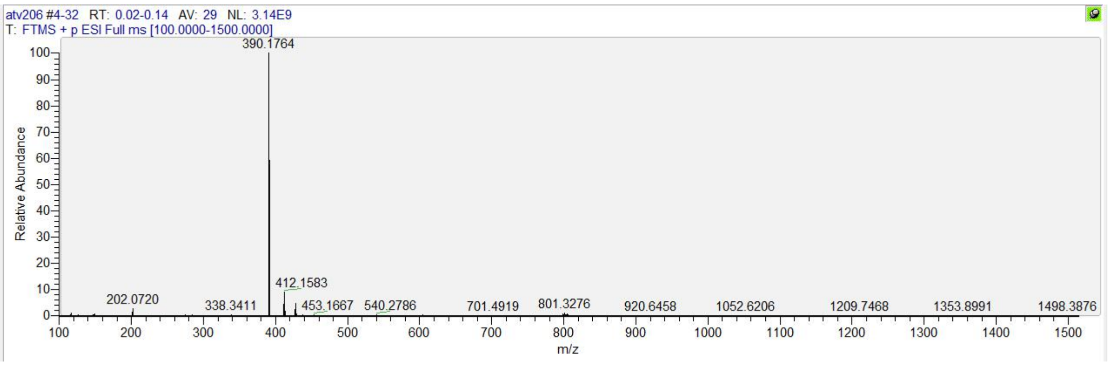

**[1]H, [13]C NMR, HRMS and HPLC spectrum of ATV012**

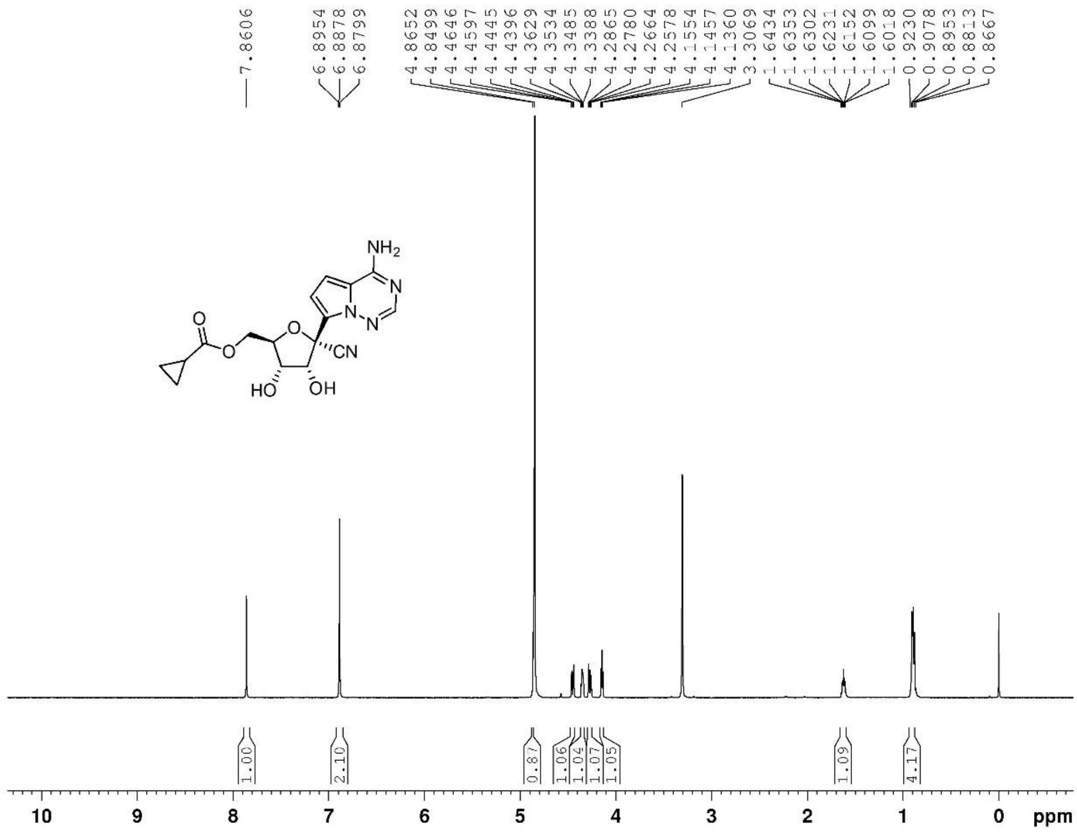

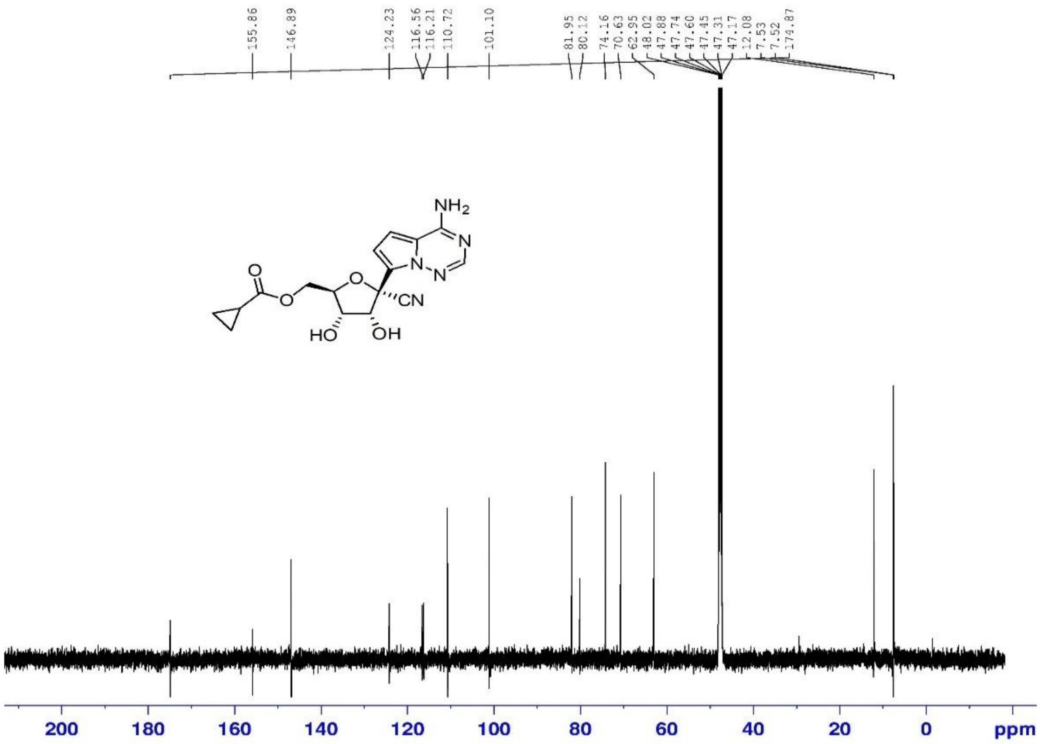

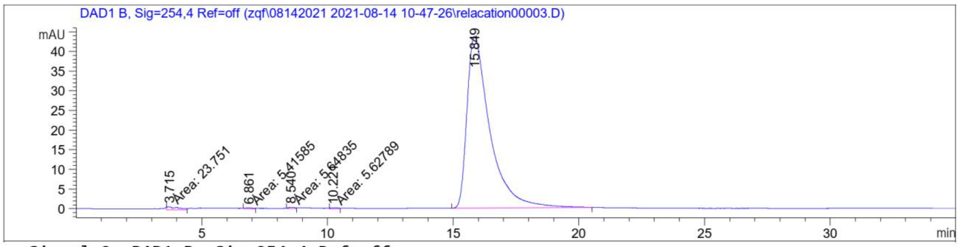

Signal 2: DAD1 B, Sig=254,4 Ref=off

| Peak # | RetTime [min] | Type | Width [min] | Area [mAU*s] | Height [mAU] | Area % |
|---|---|---|---|---|---|---|
| 1 | 3.715 | MM | 0.5380 | 23.75096 | 7.35762e-1 | 0.8450 |
| 2 | 6.861 | MM | 0.3460 | 5.41585 | 2.60856e-1 | 0.1927 |
| 3 | 8.540 | MM | 0.2972 | 5.64835 | 3.16764e-1 | 0.2010 |
| 4 | 10.221 | MM | 0.2488 | 5.62789 | 2.80089e-1 | 0.2002 |
| 5 | 15.849 | BB | 0.9217 | 2770.25024 | 43.69519 | 98.5611 |
| Totals : | | | | 2810.69330 | 45.28866 | |

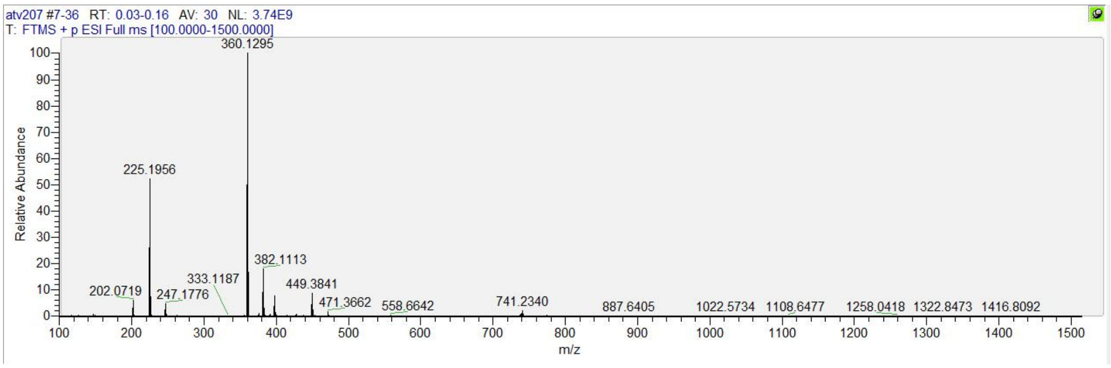

#### $^{1}$H, $^{13}$C NMR, HRMS and HPLC spectrum of ATV013

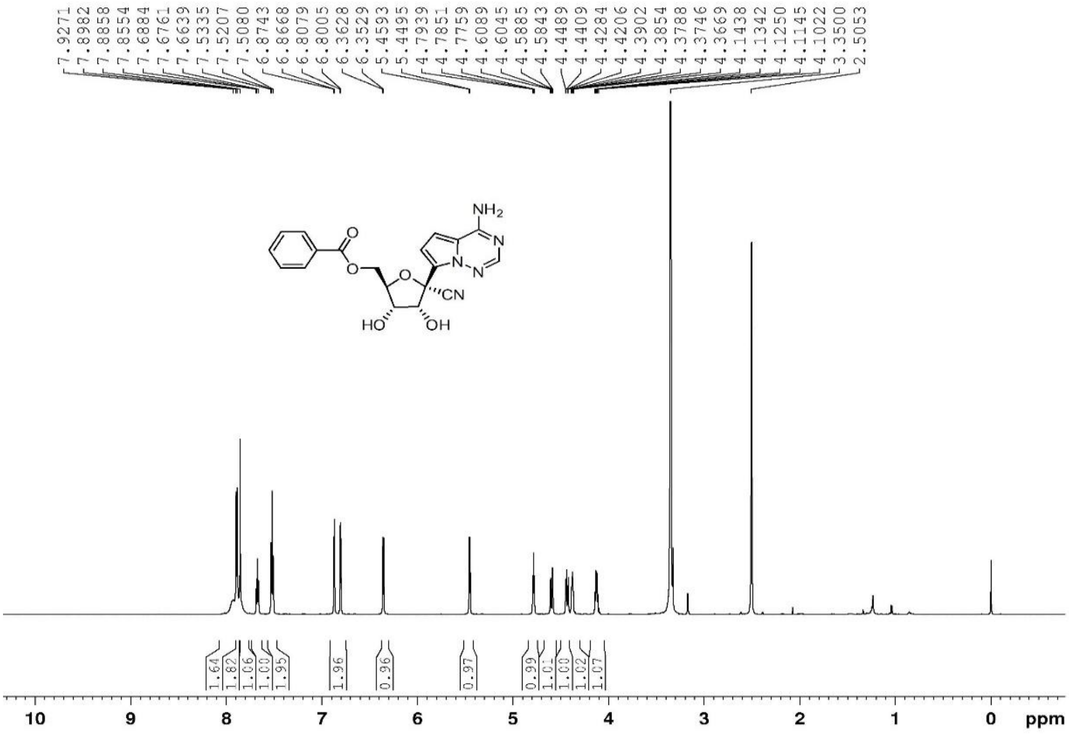

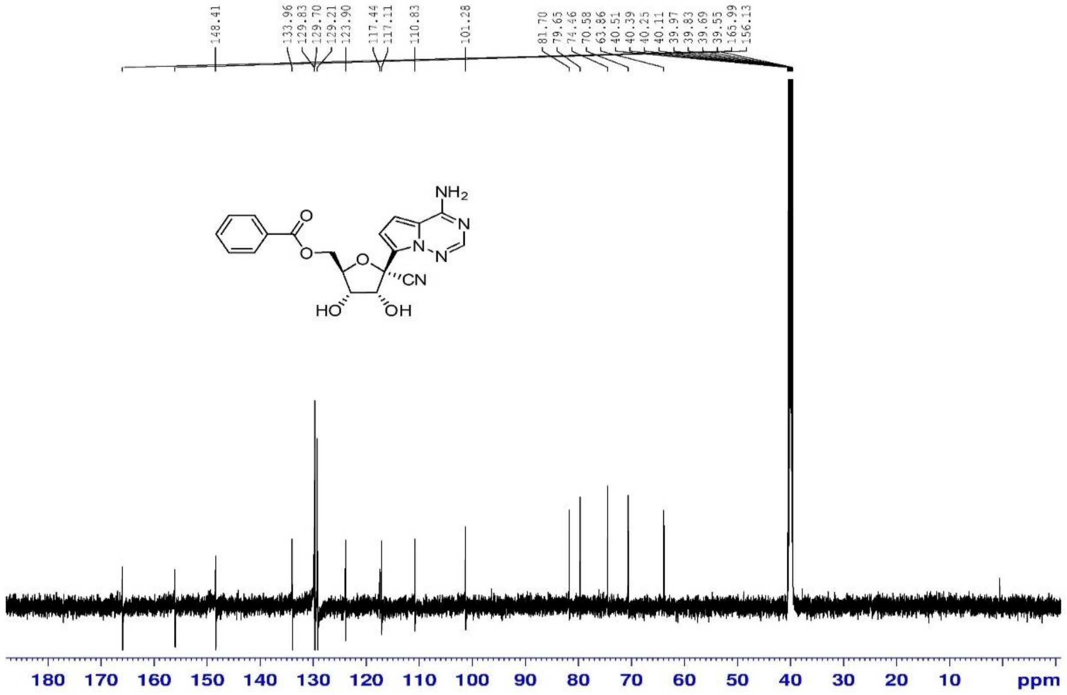

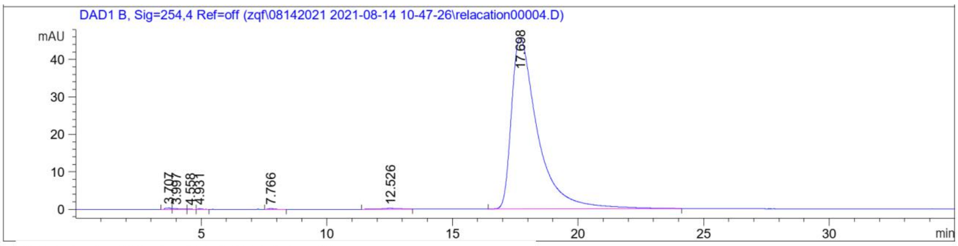

Signal 2: DAD1 B, Sig=254,4 Ref=off

| Peak # | RetTime [min] | Type | Width [min] | Area [mAU*s] | Height [mAU] | Area % |
|---|---|---|---|---|---|---|
| 1 | 3.707 | BV | 0.1964 | 5.01577 | 3.55890e-1 | 0.1507 |
| 2 | 3.997 | VV | 0.2059 | 2.94416 | 1.93067e-1 | 0.0885 |
| 3 | 4.558 | VB | 0.1134 | 6.29522e-1 | 8.59584e-2 | 0.0189 |
| 4 | 4.931 | BV R | 0.1190 | 1.23158 | 1.57861e-1 | 0.0370 |
| 5 | 7.766 | BB | 0.2539 | 3.33529 | 1.92718e-1 | 0.1002 |
| 6 | 12.526 | BB | 0.6508 | 10.59647 | 1.94145e-1 | 0.3184 |
| 7 | 17.698 | BB | 1.0610 | 3304.55225 | 45.62620 | 99.2863 |

Totals : 3328.30504 46.80584

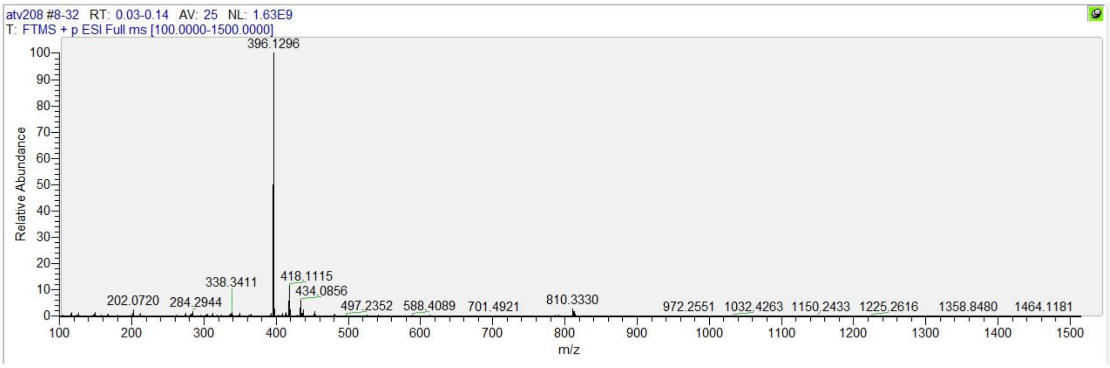

**[1]H, [13]C NMR, HRMS and HPLC spectrum of ATV015**

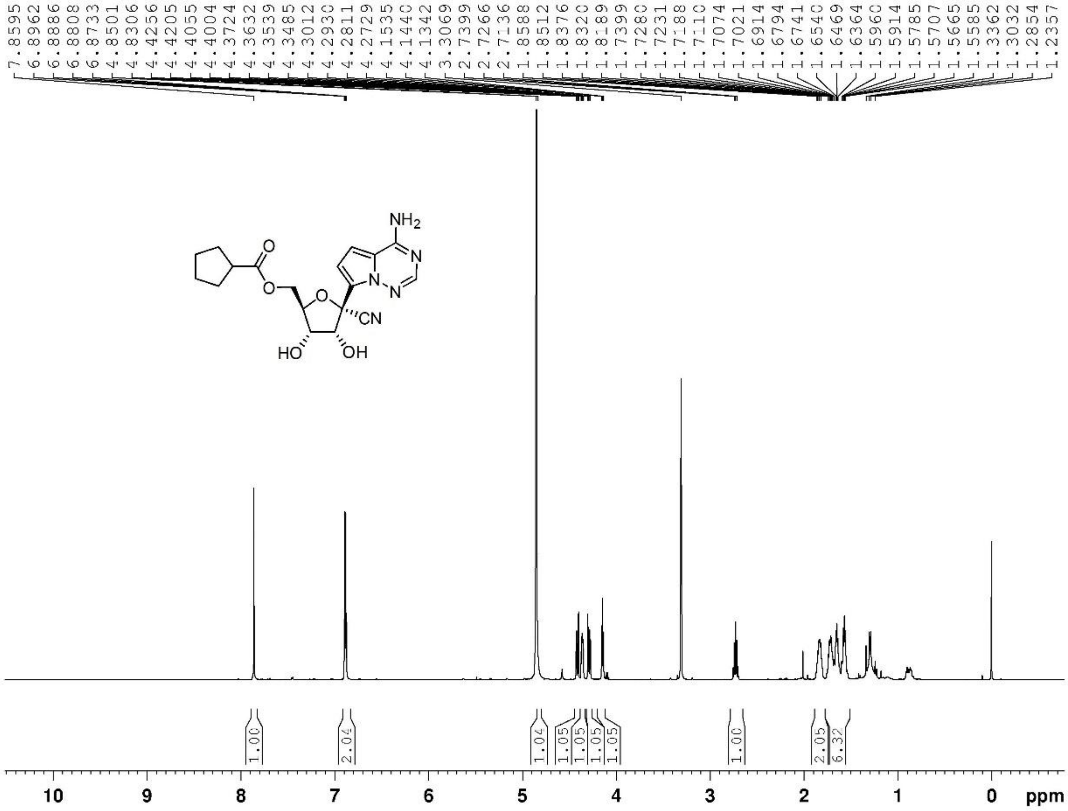

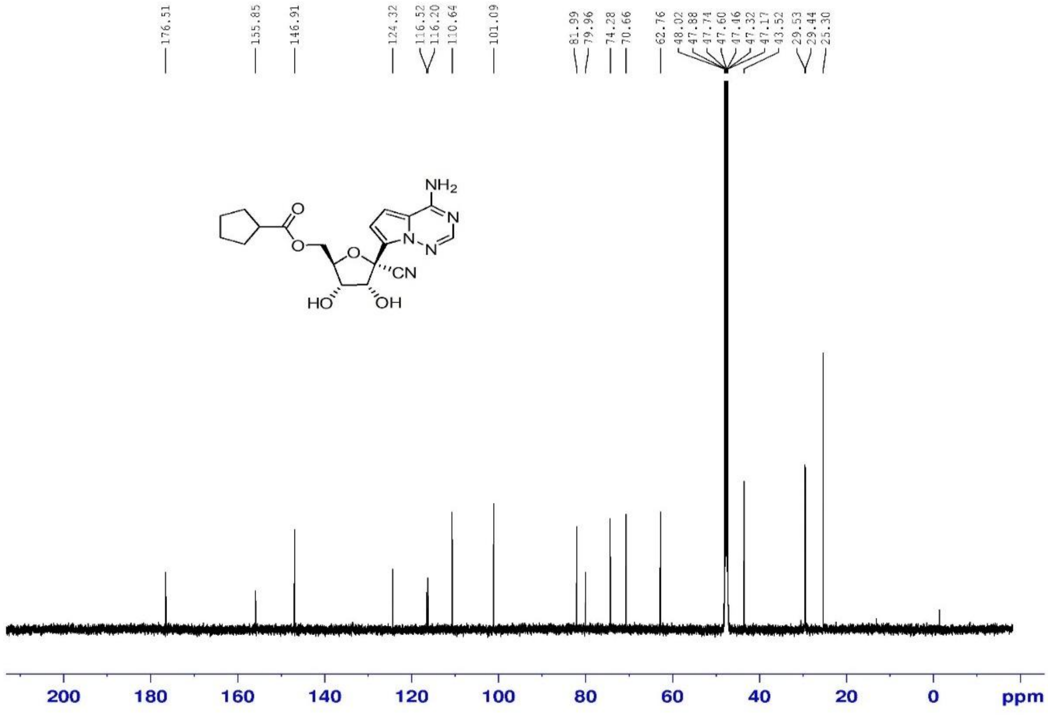

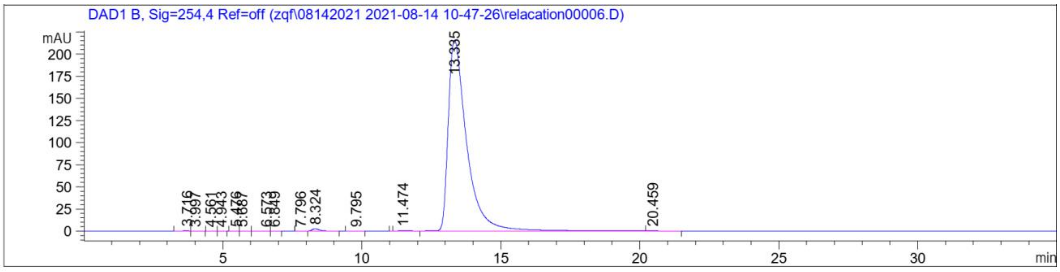

Signal 2: DAD1 B, Sig=254,4 Ref=off

| Peak # | RetTime [min] | Type | Width [min] | Area [mAU*s] | Height [mAU] | Area % |
|---|---|---|---|---|---|---|
| 1 | 3.716 | BV | 0.1956 | 6.21968 | 4.38130e-1 | 0.0619 |
| 2 | 3.997 | VB | 0.1860 | 2.84919 | 2.00195e-1 | 0.0284 |
| 3 | 4.561 | BB | 0.1091 | 7.11727e-1 | 9.31546e-2 | 7.085e-3 |
| 4 | 4.943 | BB | 0.1152 | 1.10662 | 1.48079e-1 | 0.0110 |
| 5 | 5.476 | BV | 0.1303 | 6.42078e-1 | 7.18377e-2 | 6.392e-3 |
| 6 | 5.687 | VB | 0.1969 | 8.21240e-1 | 5.53989e-2 | 8.175e-3 |
| 7 | 6.573 | BV | 0.1659 | 1.02436 | 8.66425e-2 | 0.0102 |
| 8 | 6.849 | VB | 0.1406 | 2.03975 | 2.11261e-1 | 0.0203 |
| 9 | 7.796 | BV E | 0.2022 | 1.69268 | 1.20277e-1 | 0.0169 |
| 10 | 8.324 | VB R | 0.2765 | 48.62589 | 2.61974 | 0.4841 |
| 11 | 9.795 | BB | 0.2468 | 1.43522 | 7.19271e-2 | 0.0143 |
| 12 | 11.474 | BV E | 0.3822 | 7.47619 | 2.64123e-1 | 0.0744 |
| 13 | 13.335 | VV R | 0.6894 | 9967.29785 | 215.58659 | 99.2234 |
| 14 | 20.459 | VB E | 0.4195 | 3.36857 | 9.88593e-2 | 0.0335 |

Totals : 1.00453e4 220.06622

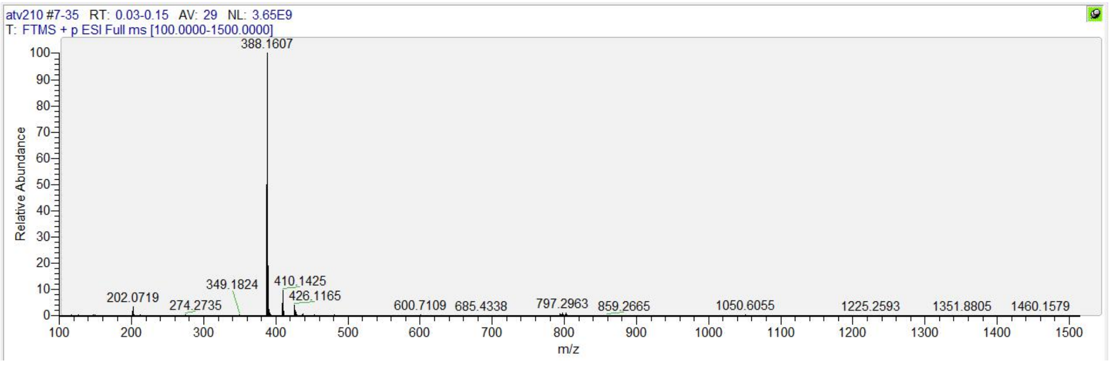

**[1]H, [13]C NMR, HRMS and HPLC spectrum of ATV016**

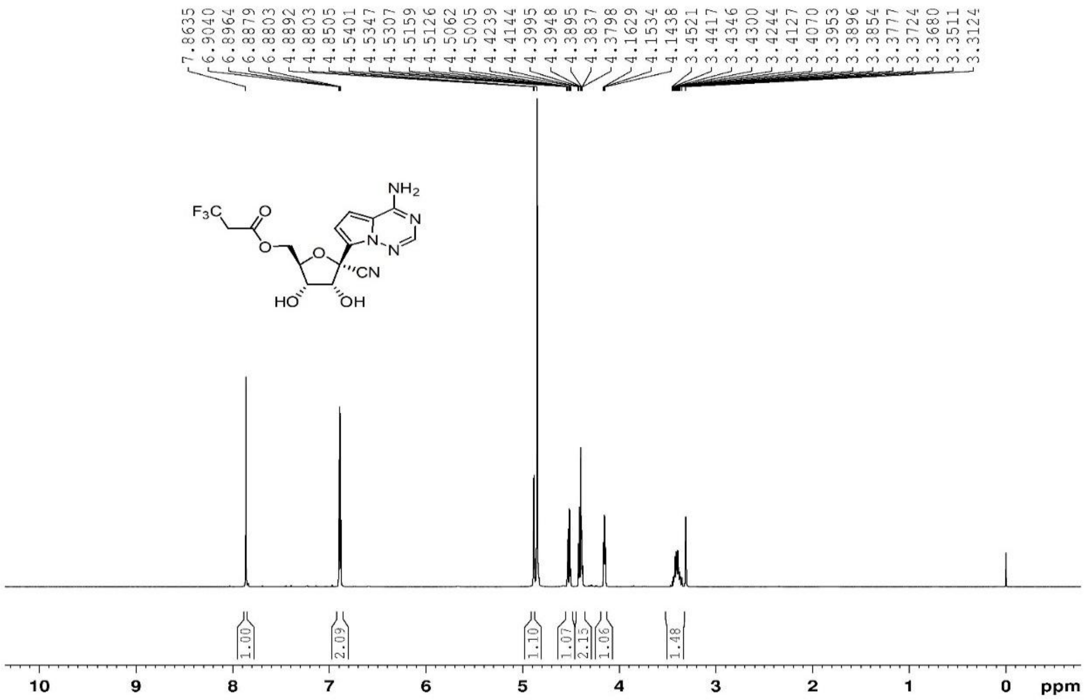

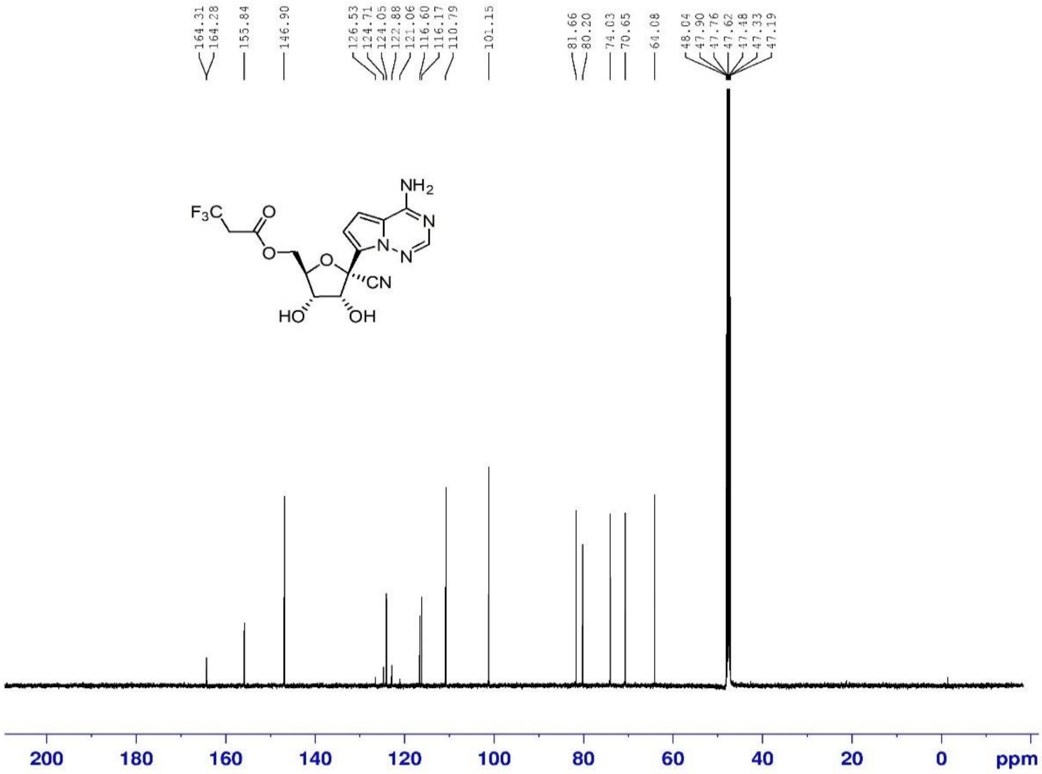

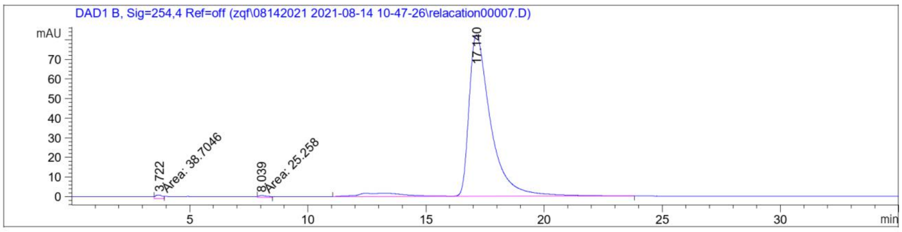

Signal 2: DAD1 B, Sig=254,4 Ref=off

| Peak # | RetTime [min] | Type | Width [min] | Area [mAU*s] | Height [mAU] | Area % |
|---|---|---|---|---|---|---|
| 1 | 3.722 | MM | 0.3557 | 38.70457 | 1.81369 | 0.6994 |
| 2 | 8.039 | MM | 0.4405 | 25.25796 | 9.55747e-1 | 0.4564 |
| 3 | 17.140 | VB R | 0.9499 | 5469.86768 | 82.32418 | 98.8442 |
| Totals : | | | | 5533.83021 | 85.09362 | |

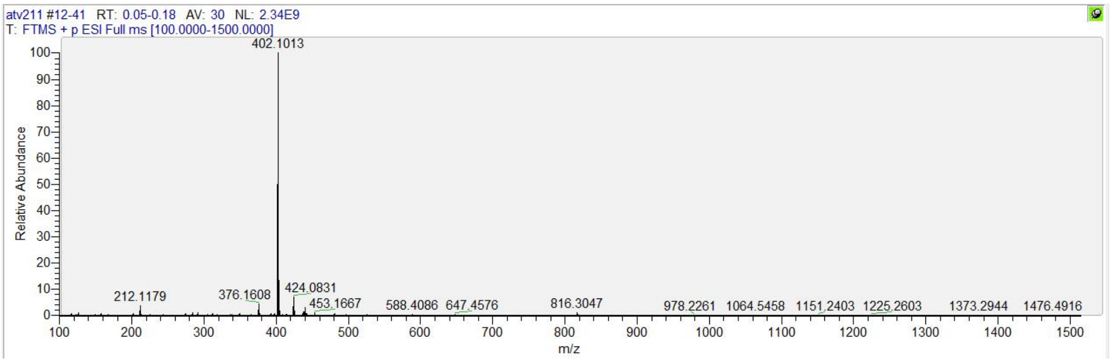

**$^1$H, $^{13}$C NMR, HRMS and HPLC spectrum of ATV017**

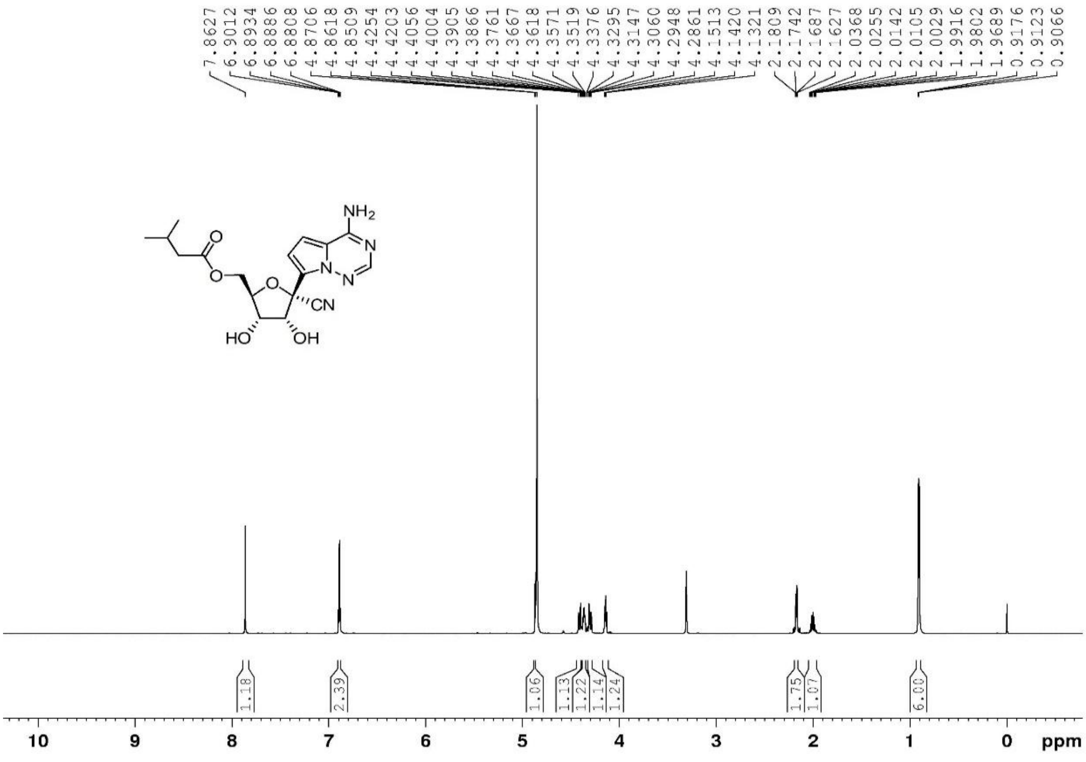

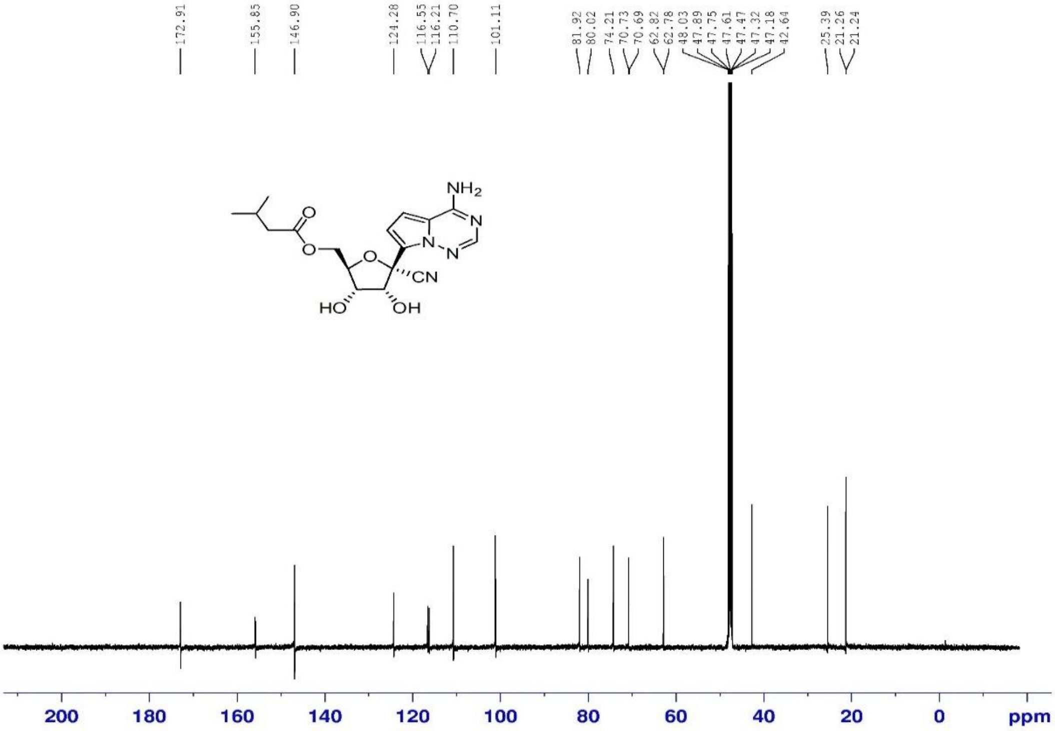

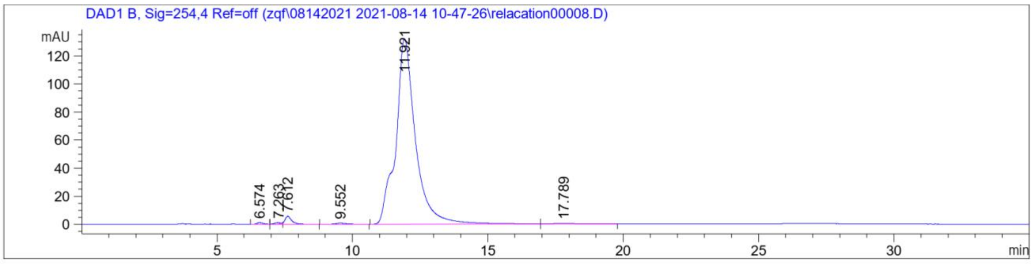

Signal 2: DAD1 B, Sig=254,4 Ref=off

| Peak # | RetTime [min] | Type | Width [min] | Area [mAU*s] | Height [mAU] | Area % |
|---|---|---|---|---|---|---|
| 1 | 6.574 | BV | 0.2480 | 19.11662 | 1.17372 | 0.2779 |
| 2 | 7.263 | VV E | 0.2545 | 15.70843 | 9.82475e-1 | 0.2283 |
| 3 | 7.612 | VB R | 0.2717 | 105.98553 | 5.73313 | 1.5405 |
| 4 | 9.552 | BB | 0.4101 | 23.71460 | 8.18228e-1 | 0.3447 |
| 5 | 11.921 | BB | 0.7231 | 6697.85059 | 132.19508 | 97.3521 |
| 6 | 17.789 | BB | 0.8490 | 17.64965 | 2.46604e-1 | 0.2565 |

Totals : 6880.02542 141.14924

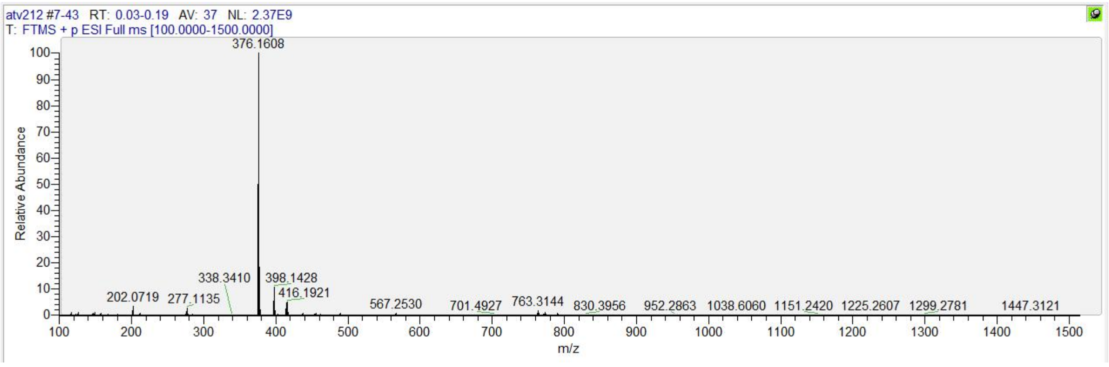

[1]H, [13]C NMR, HRMS and HPLC spectrum of ATV019

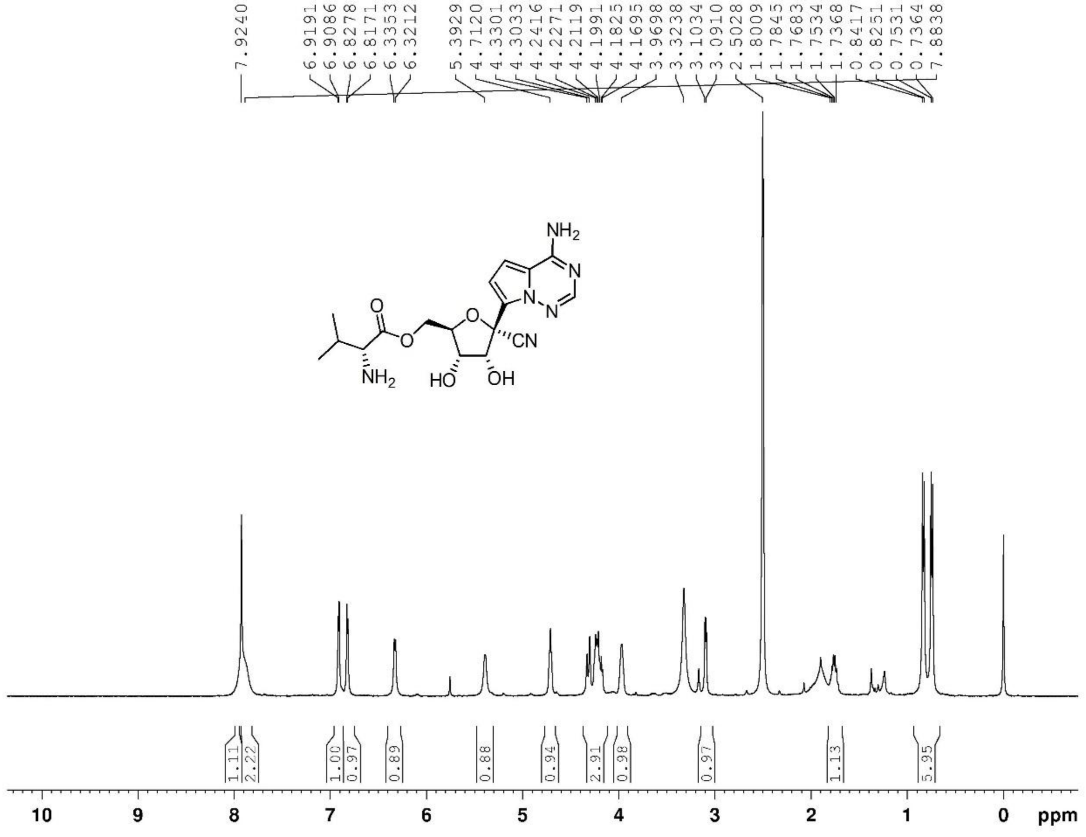

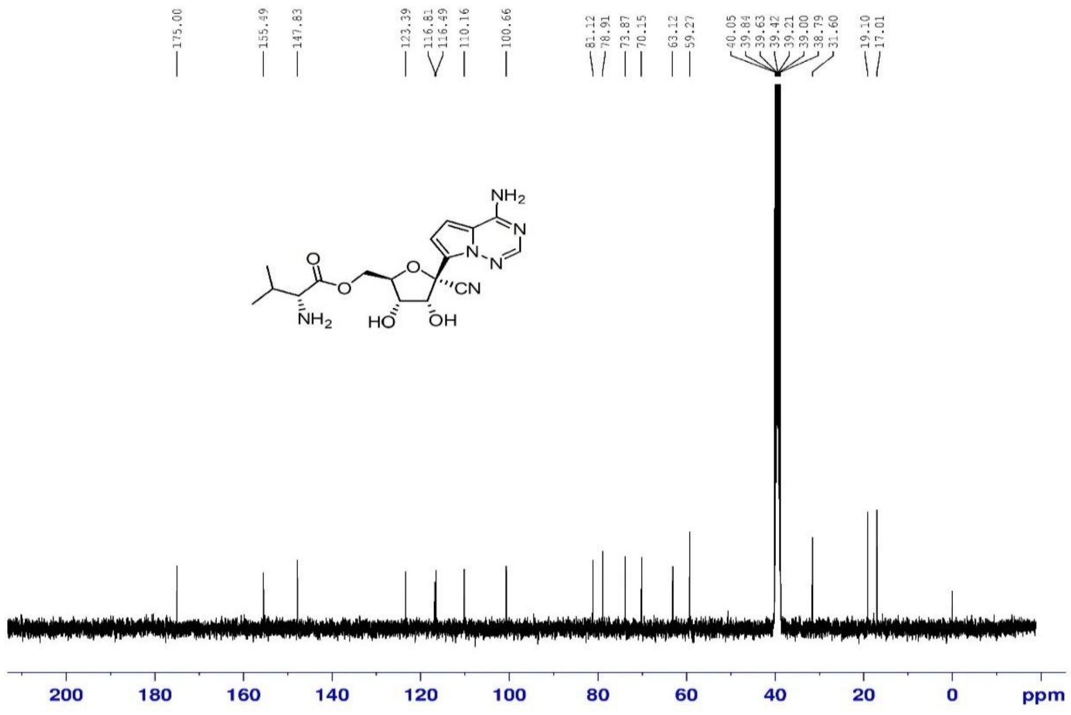

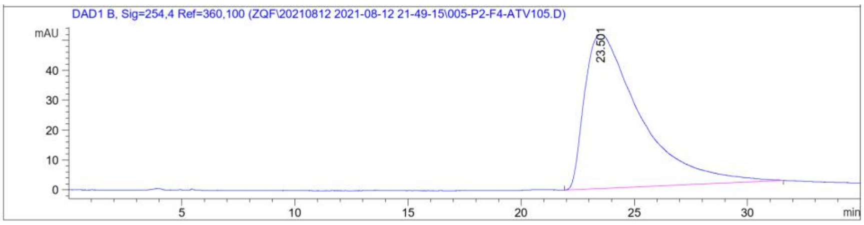

```
Signal 3: DAD1 B, Sig=254,4 Ref=360,100

Peak RetTime Type  Width     Area      Height     Area
  #   [min]        [min]   [mAU*s]     [mAU]       %
----|-------|----|-------|----------|----------|--------|
   1  23.501 BB    2.0173 8807.49219   51.34311 100.0000

Totals :                  8807.49219   51.34311
```

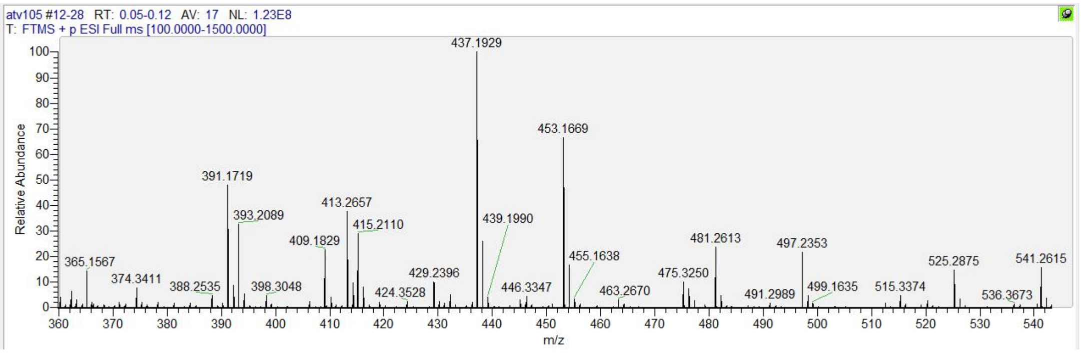

#### $^1H$, $^{13}C$ NMR, HRMS and HPLC spectrum of ATV020

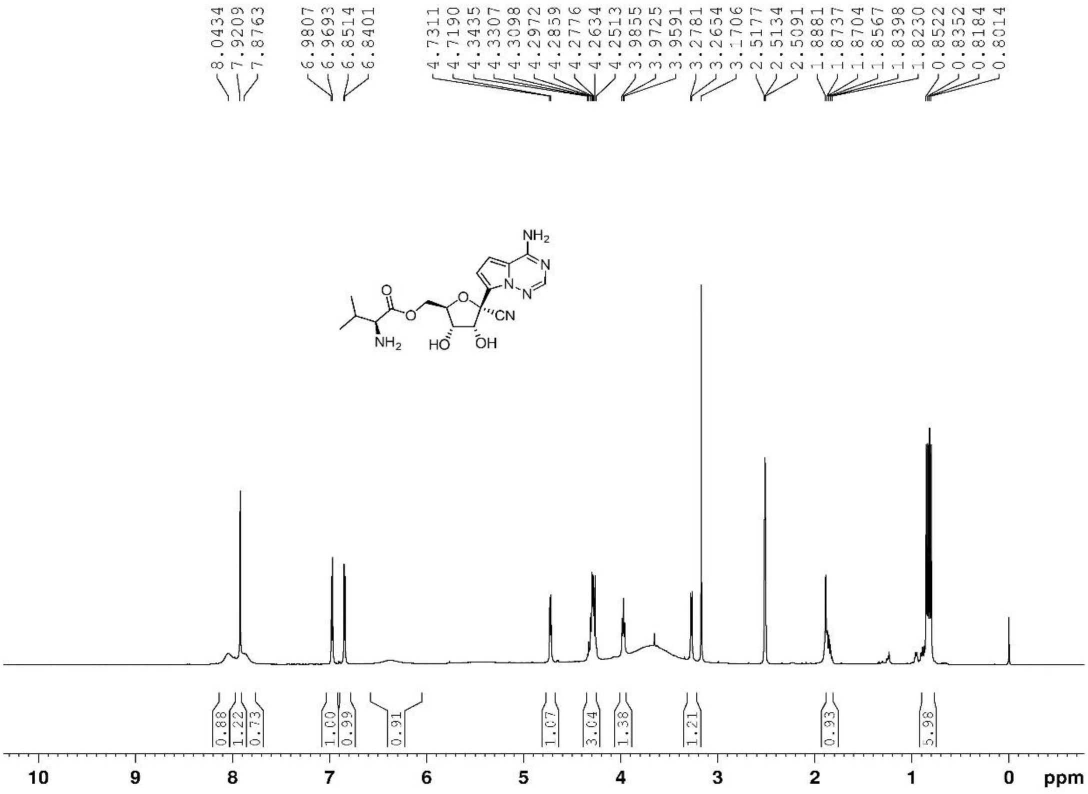

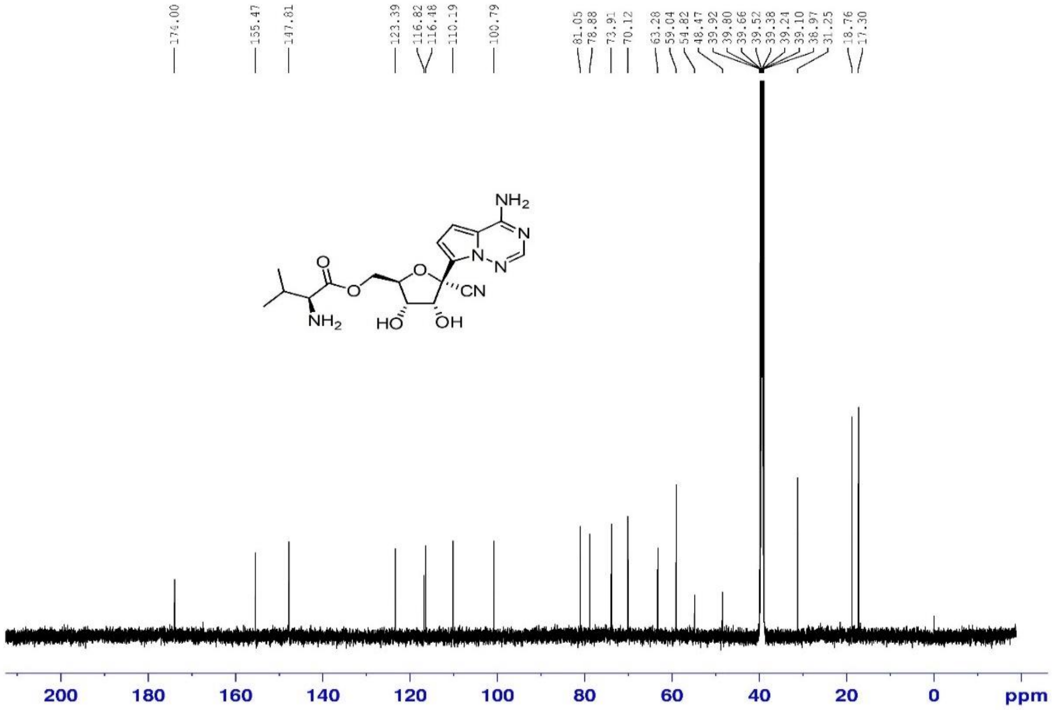

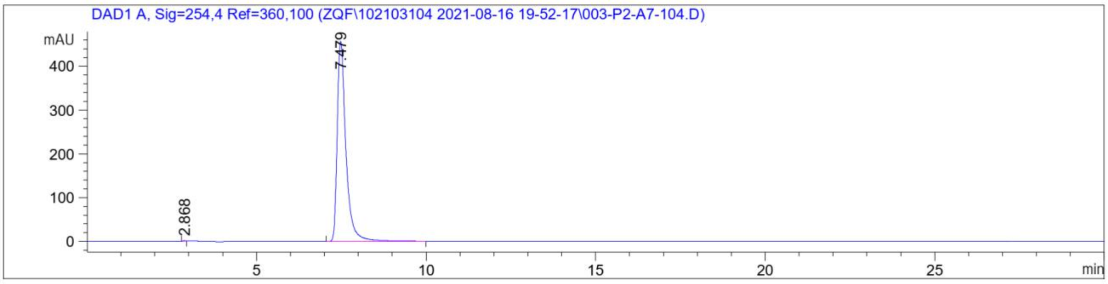

Signal 1: DAD1 A, Sig=254,4 Ref=360,100

| Peak # | RetTime [min] | Type | Width [min] | Area [mAU*s] | Height [mAU] | Area % |
|---|---|---|---|---|---|---|
| 1 | 2.868 | BB | 0.0725 | 5.09173 | 1.13117 | 0.0626 |
| 2 | 7.479 | BB | 0.2658 | 8131.78857 | 456.65549 | 99.9374 |

Totals : 8136.88030 457.78666

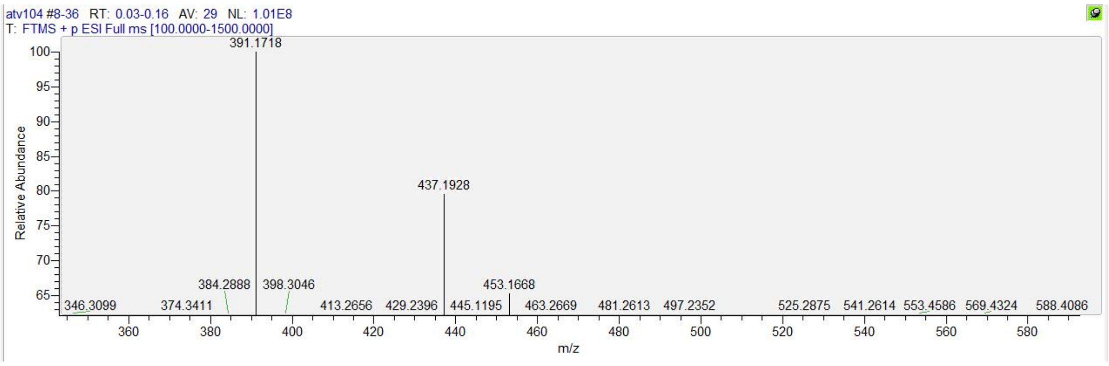

$^{1}H$, $^{13}C$ NMR, HRMS and HPLC spectrum of ATV021

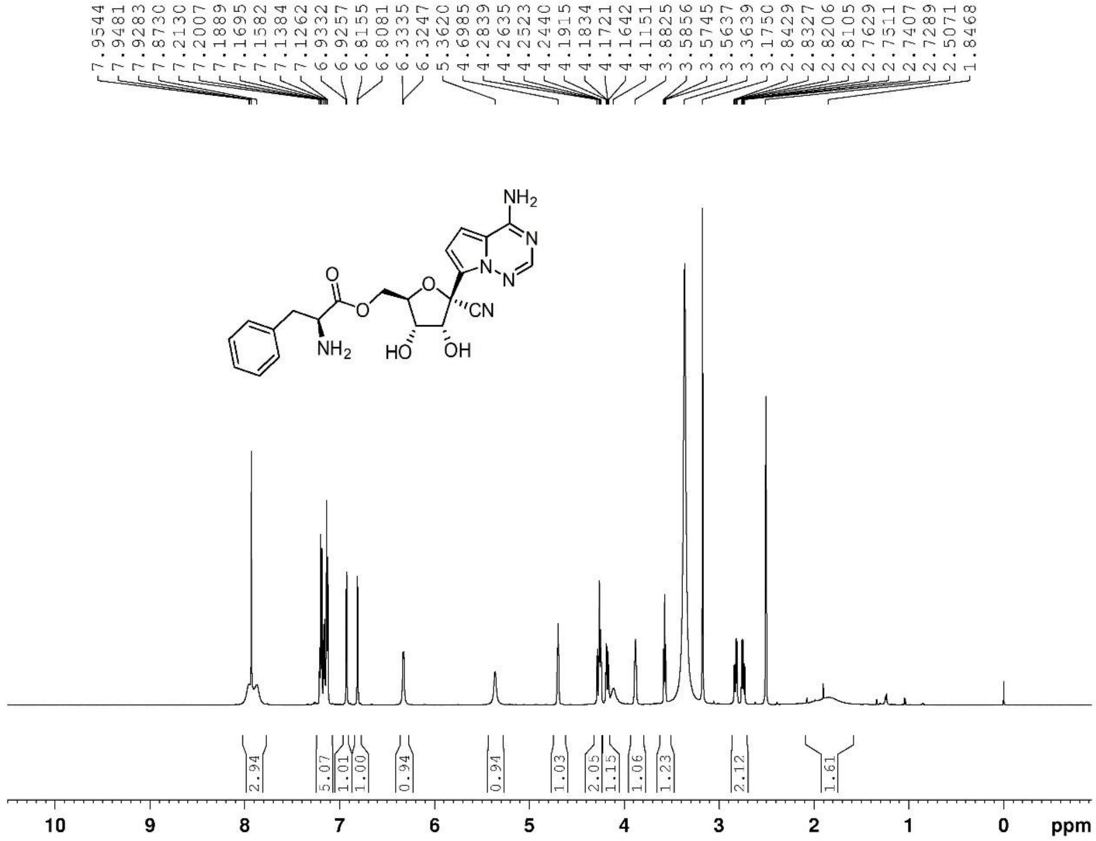

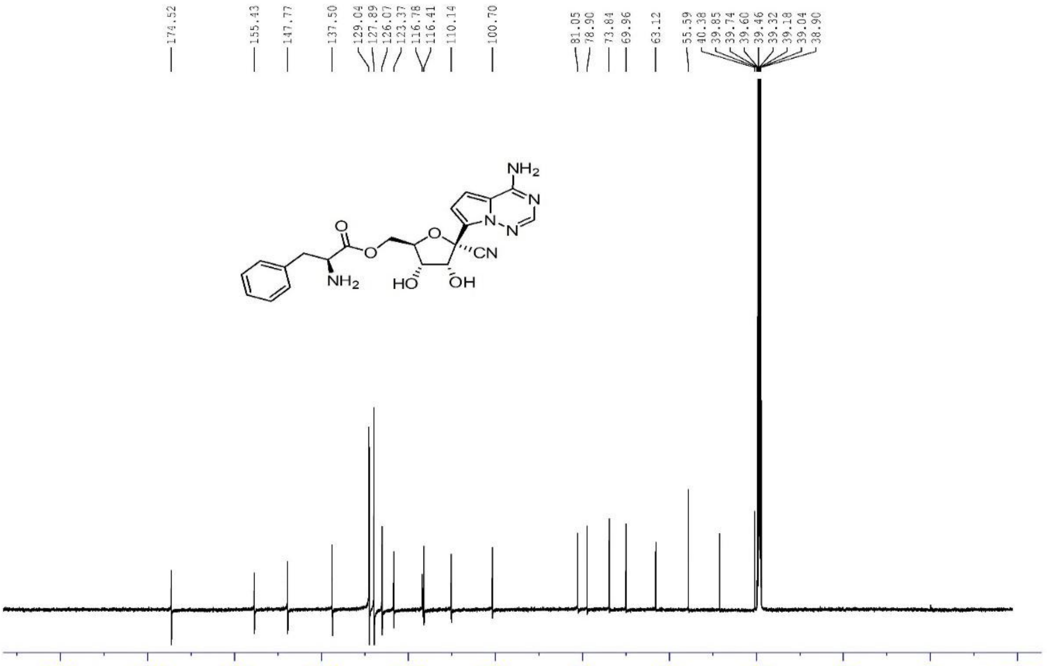

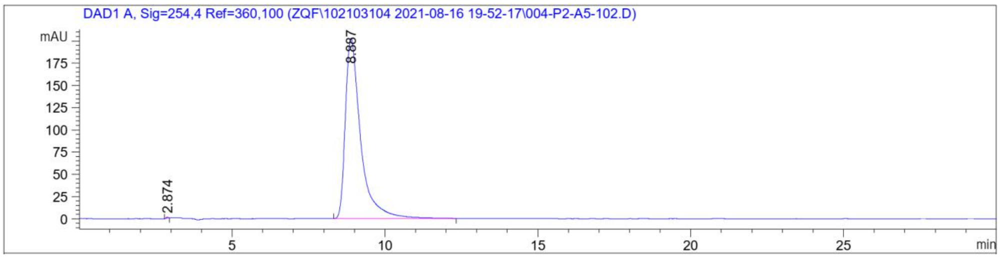

| Peak # | RetTime [min] | Type | Width [min] | Area [mAU*s] | Height [mAU] | Area % |
|---|---|---|---|---|---|---|
| 1 | 2.874 | BB | 0.0776 | 5.40199 | 1.13428 | 0.0766 |
| 2 | 8.887 | BB | 0.5173 | 7049.84961 | 202.91095 | 99.9234 |
| Totals : | | | | 7055.25160 | 204.04523 | |

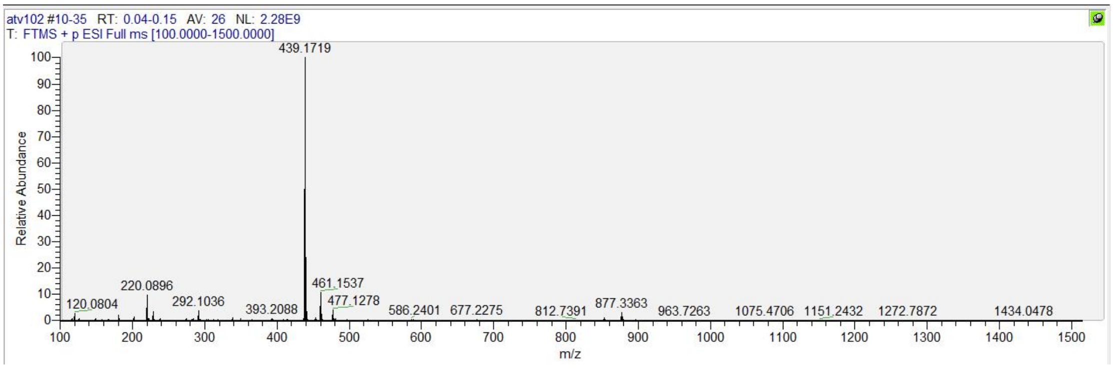

**$^1$H, $^{13}$C NMR, HRMS and HPLC spectrum of ATV022**

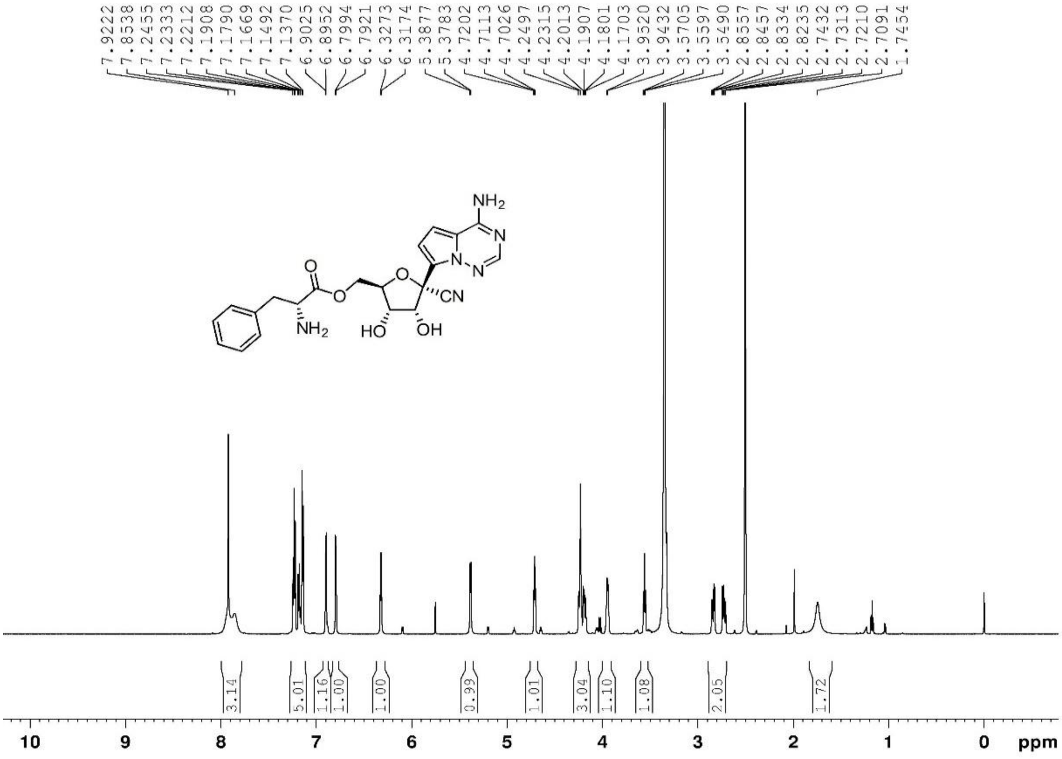

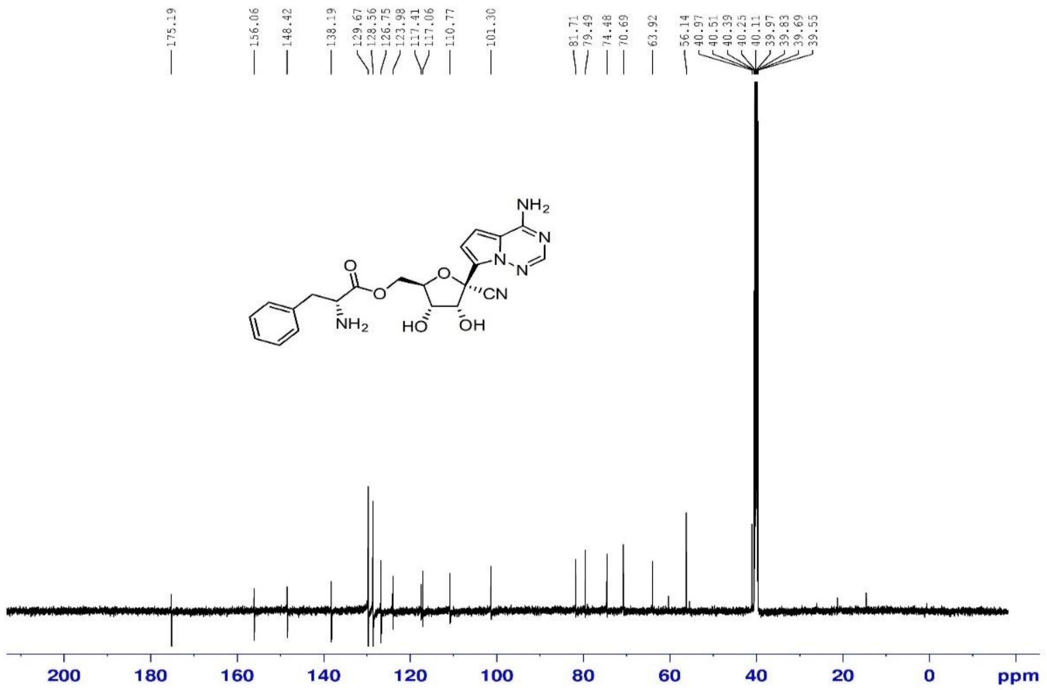

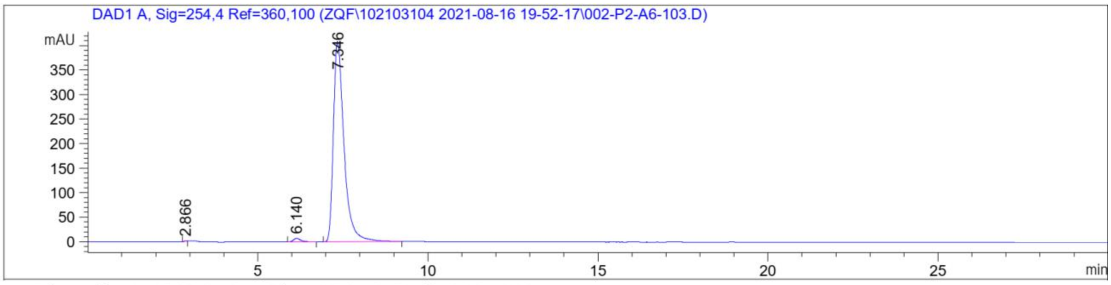

Signal 1: DAD1 A, Sig=254,4 Ref=360,100

| Peak # | RetTime [min] | Type | Width [min] | Area [mAU*s] | Height [mAU] | Area % |
|---|---|---|---|---|---|---|
| 1 | 2.866 | BB | 0.0721 | 6.22332 | 1.39341 | 0.0692 |
| 2 | 6.140 | BB | 0.2198 | 106.34193 | 7.22068 | 1.1817 |
| 3 | 7.346 | BB | 0.3279 | 8886.73242 | 408.36743 | 98.7492 |

Totals : 8999.29767 416.98152

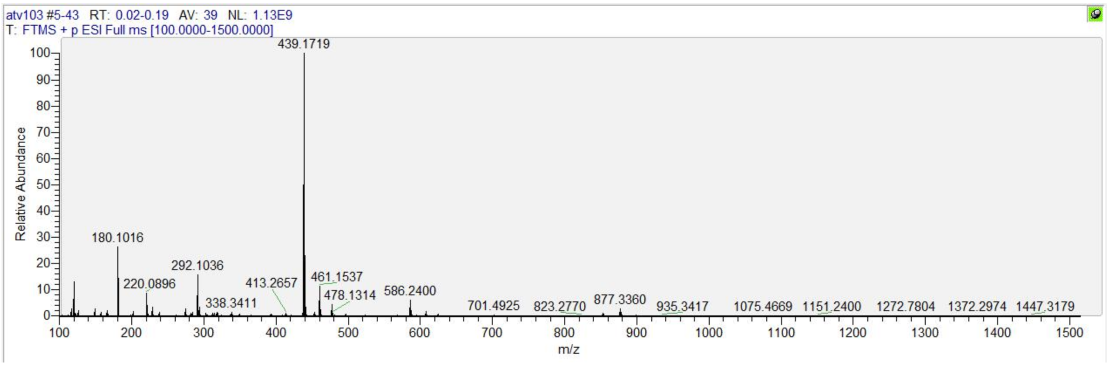

$^{1}H$, $^{13}C$ NMR, HRMS and HPLC spectrum of ATV023

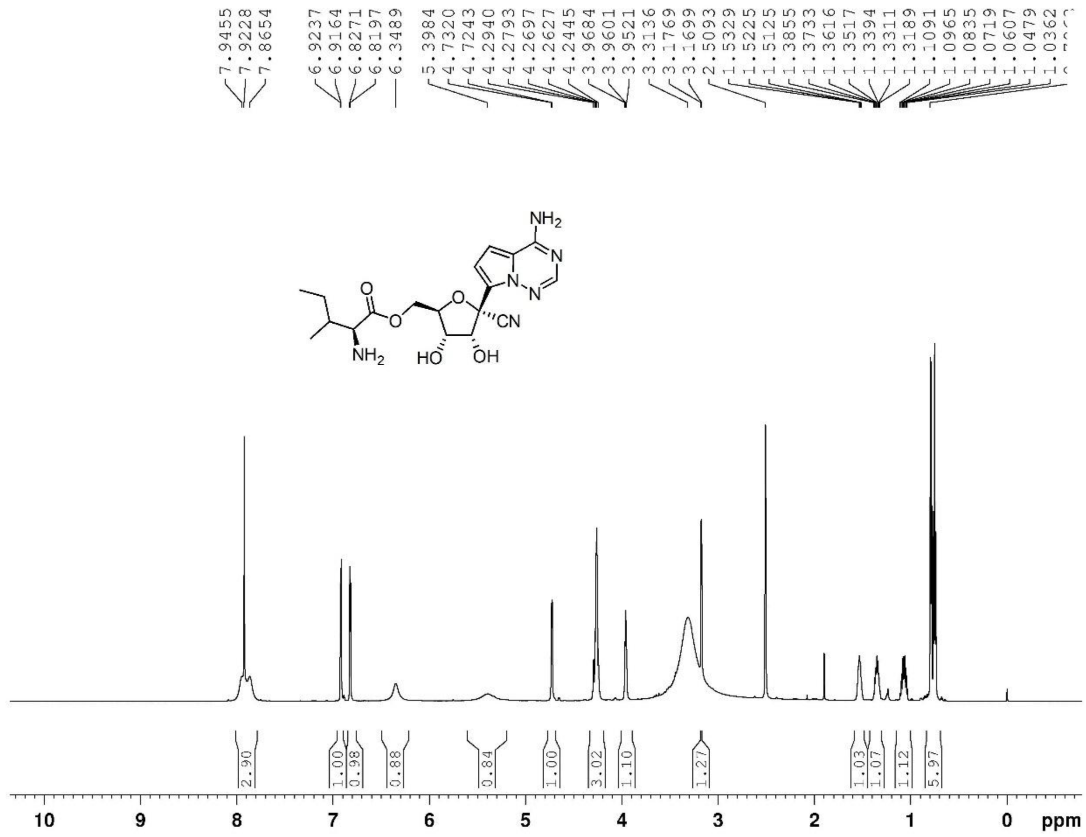

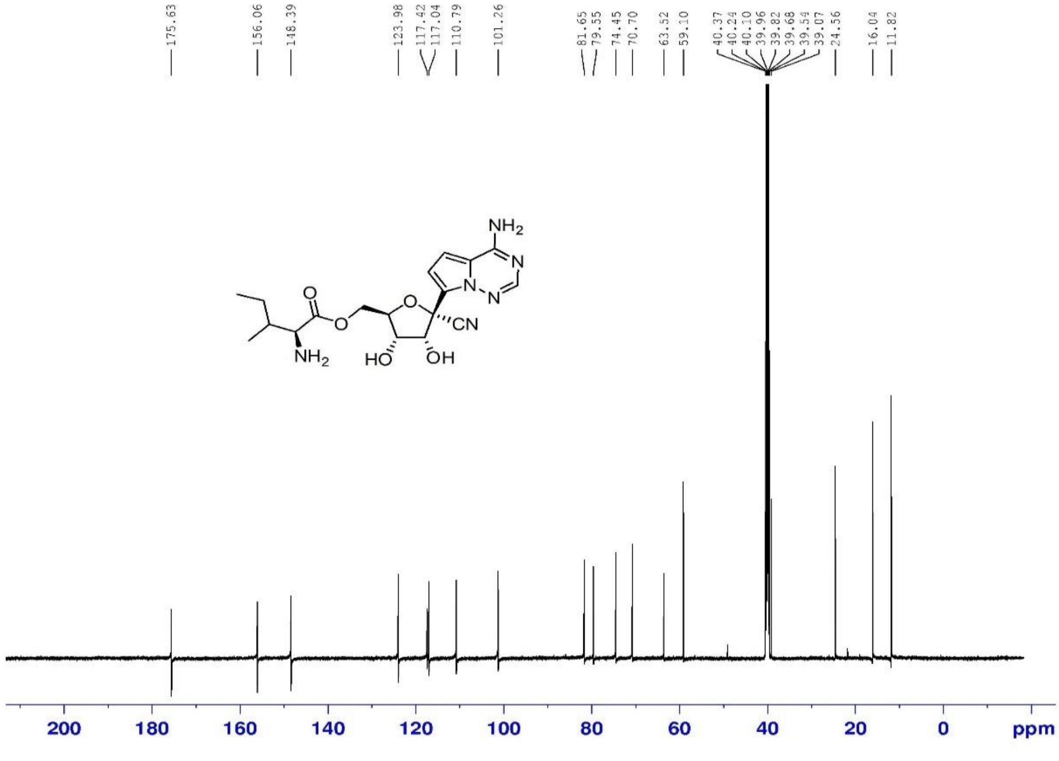

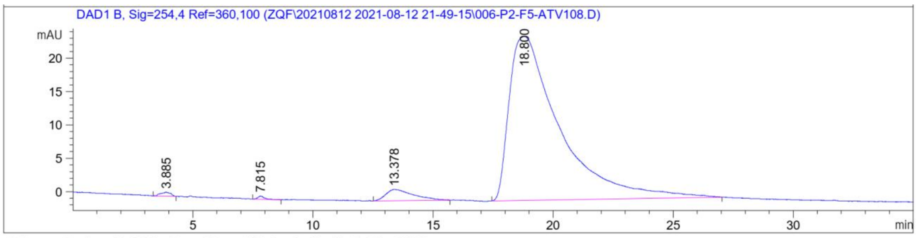

Signal 3: DAD1 B, Sig=254,4 Ref=360,100

| Peak # | RetTime [min] | Type | Width [min] | Area [mAU*s] | Height [mAU] | Area % |
|---|---|---|---|---|---|---|
| 1 | 3.885 | BB | 0.3693 | 17.95588 | 6.06681e-1 | 0.4768 |
| 2 | 7.815 | BB | 0.2672 | 8.88128 | 4.56002e-1 | 0.2358 |
| 3 | 13.378 | BB | 0.9613 | 133.90427 | 1.68837 | 3.5555 |
| 4 | 18.800 | BB | 1.7229 | 3605.35791 | 24.55655 | 95.7319 |

Totals : 3766.09933 27.30761

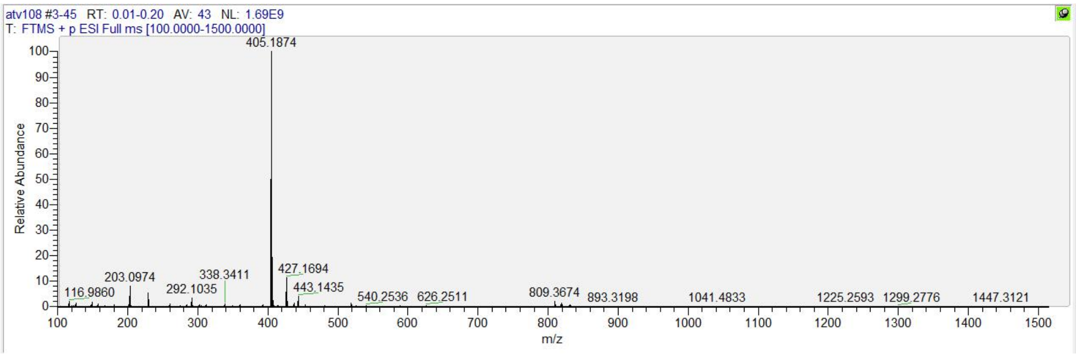

#### $^{1}H$, $^{13}C$ NMR, HRMS and HPLC spectrum of ATV024

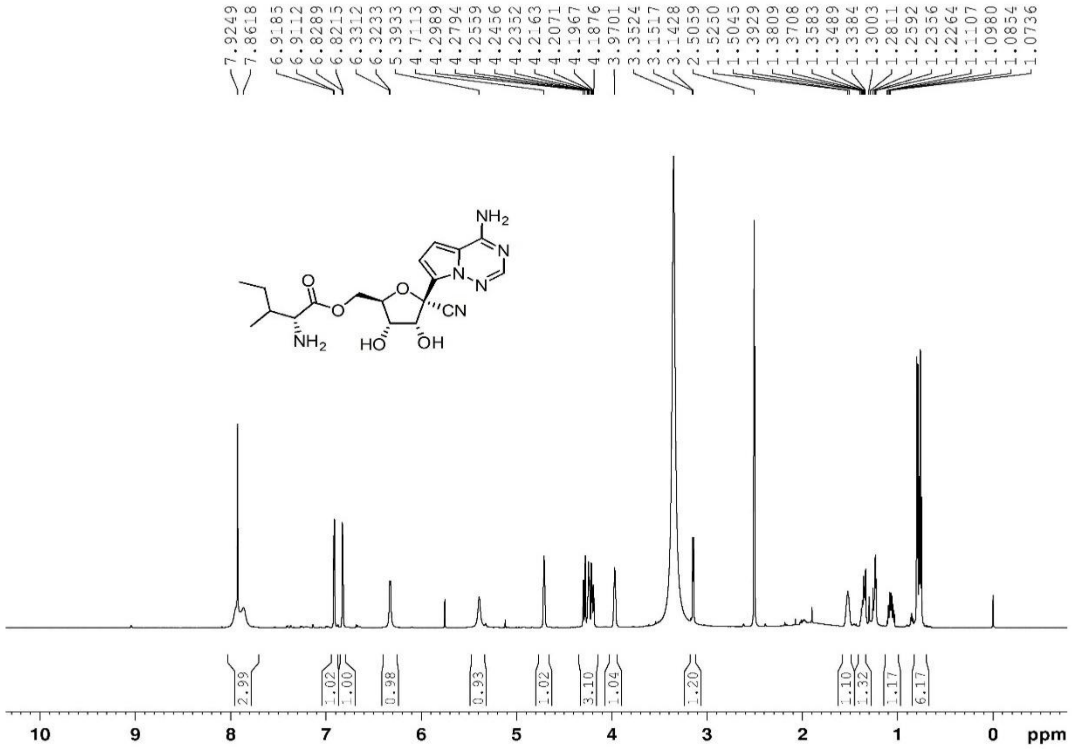

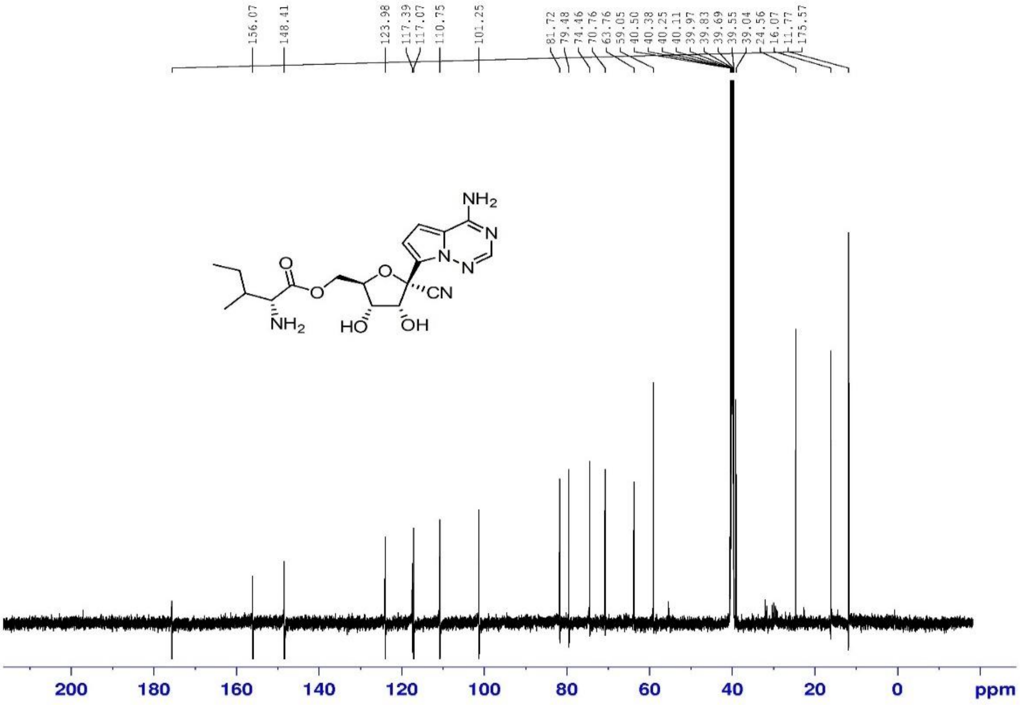

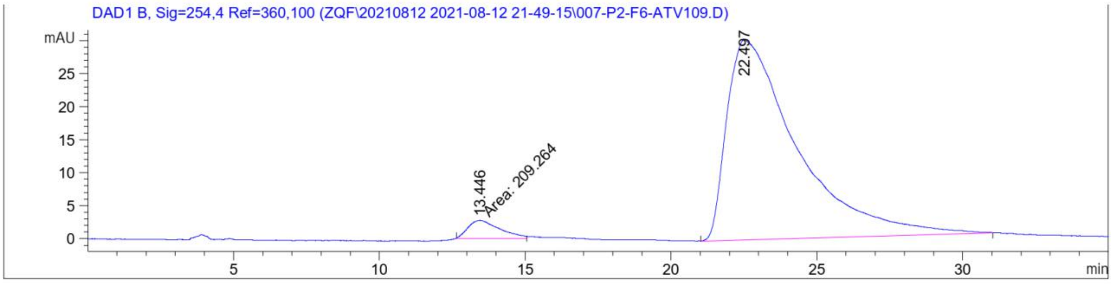

Signal 3: DAD1 B, Sig=254,4 Ref=360,100

| Peak # | RetTime [min] | Type | Width [min] | Area [mAU*s] | Height [mAU] | Area % |
|---|---|---|---|---|---|---|
| 1 | 13.446 | MM | 1.2664 | 209.26396 | 2.75409 | 3.9522 |
| 2 | 22.497 | BB | 2.1710 | 5085.57080 | 30.31805 | 96.0478 |
| Totals : | | | | 5294.83476 | 33.07214 | |

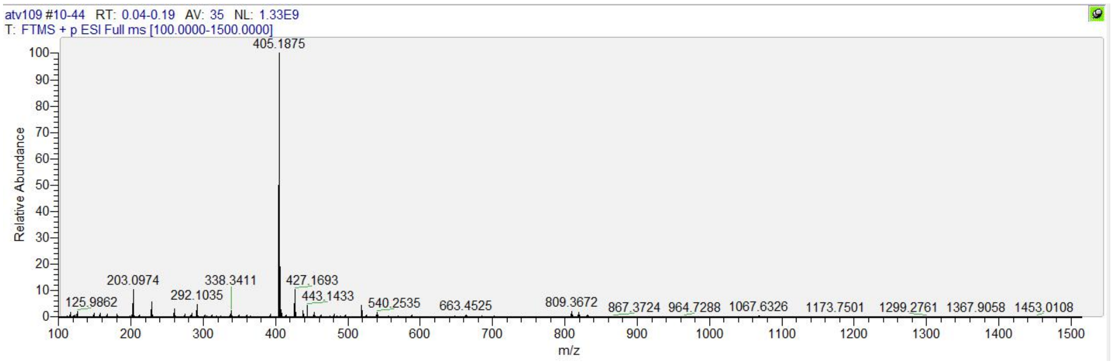
